# Spatial remodeling of bone marrow architecture defines tissue-state signatures of disease activity and therapeutic response in myelodysplastic neoplasms

**DOI:** 10.1101/2025.10.02.680138

**Authors:** Ryan Nachman, Aleksandra Kopacz, Caitlin Unkenholz, Jian Chai, Arvin Ruiz, Itzel Valencia, Jeanne Jiang, Fabio Socciarelli, Jiwoon Park, Christopher E. Mason, Ling Zhang, David Sallman, Gail J. Roboz, Pinkal Desai, Justin Kaner, Joshua Fein, Monica Guzman, Neal Lindeman, Amy Chadburn, Madhu Ouseph, Paul Simonson, Julia Geyer, Giorgio Inghirami, Shahin Rafii, David Redmond, Sanjay S. Patel

## Abstract

Myelodysplastic neoplasms (MDS) disrupt bone marrow hematopoiesis, yet clinical assessment relies largely on blast enumeration and qualitative morphology, which incompletely capture marrow architecture and disease state. We applied whole-slide multiplex immunofluorescence imaging with single-cell phenotyping to map bone marrow microarchitecture in MDS. Diagnostic biopsies (n=36), longitudinal treatment samples (n=29), precursor states (n=13), and normal controls (n=21) were analyzed, comprising >5 million spatially resolved cells. MDS marrow exhibited coordinated, genotype-imprinted architectural remodeling, including altered progenitor composition and spatial patterning, disrupted erythroid island organization, and displacement of hematopoietic stem and progenitor cells from perivascular niches. Interrogation of 82 cellular and spatial features yielded a composite Microarchitectural Perturbation Score (MDS-MAPS), derived from diagnostic samples and fixed prior to longitudinal analyses. In leave-one-patient-out cross-validation, MDS-MAPS discriminated remission from active disease more accurately than blast percentage (AUC 0.883 vs 0.660) and distinguished low-blast MDS from clonal cytopenia of undetermined significance (CCUS) (AUC 0.815). Mixed-effects modeling showed MAPS decreased in remission statistically independent of blast burden, with architectural normalization during remission and re-emergence at relapse. These findings define quantitative bone marrow architecture as a dynamic tissue-state biomarker that complements molecular and blast-based assessment in MDS.

## INTRODUCTION

Myelodysplastic neoplasms (MDS) arise from clonal hematopoietic stem and progenitor cells and are characterized by ineffective hematopoiesis, peripheral cytopenias, and risk of progression to acute myeloid leukemia.^1^ Diagnosis and disease assessment rely heavily on histopathologic evaluation of bone marrow biopsies, including subjective assessment of dysplasia and blast enumeration, yet these approaches provide limited quantitative insight into the spatial organization of the marrow and may incompletely capture the biological state of the disease.^2,3^

Genomic profiling has substantially advanced understanding of the cell-intrinsic drivers of MDS and has improved risk stratification models.^4^ However, most molecular and immunophenotypic data are derived from peripheral blood or aspirated marrow, which lack preservation of tissue architecture and spatial cellular interactions. As a result, while hundreds of thousands of cells can be profiled in suspension, the spatial organization of hematopoiesis within intact marrow—where both cell-intrinsic and microenvironmental influences converge—remains poorly quantified. Recent studies suggest that clonal hematopoiesis may remodel the marrow niche itself, altering interactions between progenitors, stromal elements, and vascular structures,^5^ but these changes have not been systematically quantified in human disease. Reliable models that integrate spatial architecture with the clinical and genetic heterogeneity of MDS are still lacking.^6,7^

We recently performed single-cell–resolved spatial mapping of human bone marrow architecture using multiplex immunofluorescence and whole-slide imaging, revealing conserved structural patterns in normal aging marrow.^8^ Here, we apply and extend this framework to clinically and genetically annotated MDS biopsies, including longitudinal samples obtained during therapy. We hypothesized that quantitative measures of bone marrow microarchitecture capture a tissue-level disease state that complements conventional metrics. To summarize these spatial features, we derived a composite Microarchitectural Perturbation Score (MDS-MAPS) and evaluated whether marrow architecture encodes clinically relevant information across diagnostic and serial samples.

## METHODS

### Study Cohort and Clinical Definitions

Archival posterior iliac crest bone marrow trephine biopsies were collected from individuals evaluated for cytopenia of unknown etiology at our academic medical center. All cases were classified according to World Health Organization (WHO) 5th edition criteria. The study was approved by the Institutional Review Board and conducted in accordance with the Declaration of Helsinki. Comprehensive details are provided in **Supp.**

### Materials

#### Clinical Laboratory Characterization

Standard clinical evaluation included morphology, multiparameter flow cytometry, cytogenetics, and targeted sequencing performed in CLIA/validated clinical workflows.

#### Multiplex Immunofluorescence Imaging

Whole-slide Opal-based multiplex immunofluorescence was performed using a previously established workflow.^8^

#### Image Segmentation and Phenotyping

Cells and structural elements were segmented and phenotyped using previously described deep-learning and image-analysis pipelines.^8^

#### Spatial Analyses

Spatial enrichment, erythroid clustering, and ALIP were quantified using prespecified spatial metrics and permutation-based testing.

#### Construction of the MDS Microarchitectural Perturbation Score (MDS-MAPS)

MDS-MAPS was derived from an interrogation of 82 initial prespecified cellular, morphologic, and spatial features using diagnostic samples only, and the model was locked before longitudinal analyses (see *Results* section for detailed derivation).

#### CCUS/MDS-LB Discrimination and Remission Classification Modeling

Diagnostic and remission classification were evaluated with patient-level leave-one-patient-out cross-validation and compared with blast- and mutation-based comparators.

#### Mixed-Effects Modeling

To evaluate whether MAPS reductions occurred independently of blast burden, a linear mixed-effects model was fit:

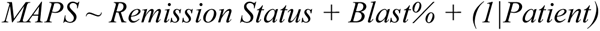

using statsmodels in Python. A random intercept for patient was included; random slopes were not modeled due to limited within-patient sample size.

#### Re-analysis of Public Datasets

Public transcriptomic/proteomic datasets were re-analyzed to evaluate CXCR4 expression in MDS HSPCs.

#### Statistical Analysis

All analyses were performed in Python (v3.11). Non-parametric comparisons used the Mann–Whitney U test, correlations used Spearman’s rank coefficient, and multiple testing correction was applied using Benjamini–Hochberg (FDR < 0.05) where appropriate.

## RESULTS

### Characteristics of patient cohort

We assembled a cohort of clinically and genetically annotated archival bone marrow trephine biopsy samples collected as part of routine diagnostics at our academic medical center **(Figure 1a, Supp. Data Table 1)**. The cohort included 77 samples from 49 unique patients with either MDS (n=36) or the precursor state clonal cytopenia of unknown significance (CCUS, n=13); additionally, for 14 of the MDS patients, 29 serial samples collected along the course of pharmacologic management, with or without eventual allogeneic hematopoietic stem cell transplantation (allo-HCT), were also profiled. 21 normal bone marrow controls (NBMs), including age-matched samples, from a recently published cohort were also analyzed.^8^ All cross-sectional analyses were performed on diagnostic samples unless otherwise stated (Figures 1-6 data). Remission classification analyses included only MDS samples with annotated clinical status (Figure 7 data).

**Figure 1.**
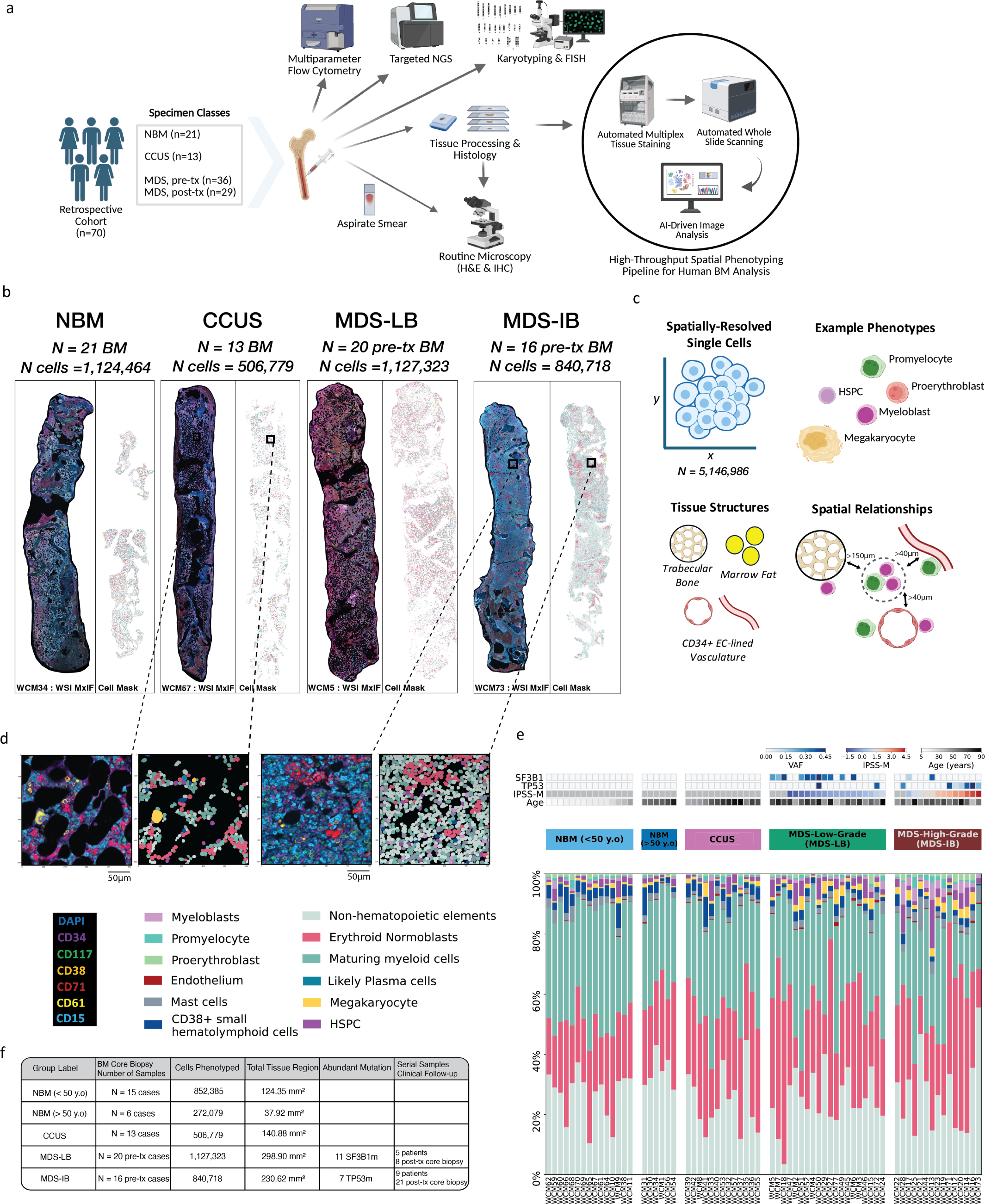
Study design and single-cell spatial profiling of bone marrow trephine biopsies. **(a)** Overview of cohort assembly and multiplex immunofluorescence (MxIF) workflow for whole-slide analysis of bone marrow trephine biopsies. **(b)** Representative whole-slide MxIF images and corresponding single-cell segmentation maps from treatment-naïve samples. MDS cases were categorized as low blast (MDS-LB) or increased blast (MDS-IB). Non-MDS samples included clonal cytopenia of undetermined significance (CCUS) and normal bone marrow (NBM; <50 years or ≥50 years). **(c)** Schematic summary of cellular, morphologic, and spatial features extracted for quantitative analysis. **(d)** Representative region of interest showing multiplex channels and corresponding phenotyped cell segmentation map from two example cases. Antibody channels and phenotype assignments are indicated. **(e)** Distribution of cell-type proportions across treatment-naïve samples. Twelve cell phenotypes were defined based on membrane marker expression and morphologic features. **(f)** Cohort overview including number of cases per group, total cells phenotyped, tissue area analyzed, most frequent mutation per disease group, and number of serial samples included in the longitudinal cohort.

The median age among CCUS and MDS patients was 77, with a M:F of 1:1. MDS samples were categorized into those with low-blasts (MDS-LB, n=20) or with high-blasts (MDS-IB, n=16). International Prognostic Scoring System-Molecular (IPSS-M) values were calculated for all diagnostic samples from MDS patients. IPSS-M scores ranged from –1.61 to 4.39 (median=0.285); cases were subclassified by risk class into Very Low (n=1), Low (n=6), Moderate Low (n=7), Moderate High (n=3), High (n=5), and Very High (n=10).

Fifty-four treatment-naive patient samples underwent targeted next-generation sequencing across the entire cohort, including 6 NBM samples (no mutations detected), all diagnostic MDS patient samples, and samples from other patients >50 years of age; data from the latter subset were used to classify patients as CCUS. The following genes were mutated in ≥10% of these 54 samples: *DNMT3A* (n=15), *SF3B1* (n=14), *ASXL1* (n=10), *TP53* (n=9), *TET2* (n=7), *SRSF2* (n=8) [**Supp. Figure 1**].

### Quantitative and morphologic variations between normal and MDS bone marrows

Following multiplex immunofluorescence (MxIF) staining and whole-slide imaging (WSI) of trephine biopsy tissues, individual cells and structural landmarks were segmented and phenotyped at single-cell resolution as previously described, yielding a dataset of 5,146,986 spatially resolved cells (**Figure 1b-f, Supp. Figure 2a-b**).^8^

**Figure 2.**
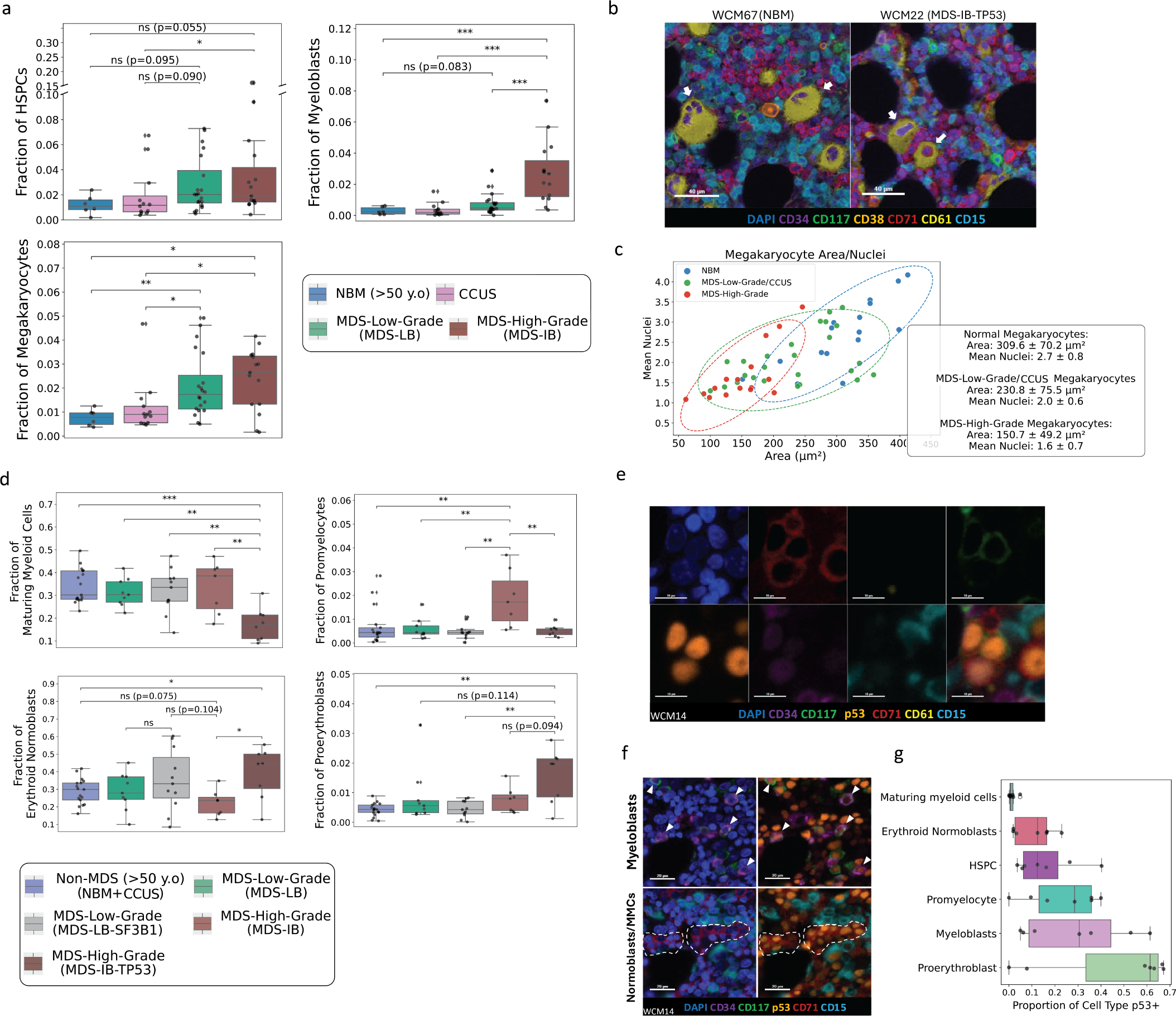
Single-cell compositional and morphologic alterations in MDS. **(a)** Proportions of hematopoietic stem and progenitor cells (HSPCs), myeloblasts, and megakaryocytes across NBM, CCUS, MDS-LB, and MDS-IB samples. Two-sided Mann–Whitney U test (*P ≤ 0.05, **P ≤ 0.01, ***P ≤ 0.001). **(b)** Representative MxIF images illustrating megakaryocyte morphology in NBM and MDS samples (scale bars, 40 µm). **(c)** Quantification of megakaryocyte area and nuclear number across disease groups. Points represent individual samples with standard-deviation ellipses. **(d)** Proportions of maturing myeloid cells (MMCs), promyelocytes, proerythroblasts, and erythroid normoblasts across disease groups, with MDS-IB stratified by *TP53* mutation status (Mann–Whitney U test; *P ≤ 0.05, **P ≤ 0.01, ***P ≤ 0.001). **(e)** Example cluster of CD117⁺CD71⁺ proerythroblasts with high p53 expression from a *TP53*-mutated case (WCM14). Top row: DAPI, CD71, CD61, CD117; bottom row: p53, CD34, CD15, merged image (scale bars, 10 µm). **(f)** Representative MxIF image showing myeloblasts and erythroid precursors with elevated p53 expression in a *TP53*-mutated sample (scale bars, 20 µm). **(g)** Proportion of cells with high p53 expression across *TP53*-mutated MDS samples in a tissue microarray, calculated as the fraction of p53⁺ cells within each phenotype.

Quantitative cell-type proportions derived from MxIF—including myeloblasts, promyelocytes, erythroid normoblasts, and maturing myeloid cells (MMCs)—correlated with orthogonal measurements from manual differential counts and multiparameter flow cytometry (**Supp. Figure 2c**, **Supp. Data Table 2**), supporting the analytical validity of the approach.

Comparative analysis across NBMs and treatment-naïve CCUS, MDS-LB, and MDS-IB samples revealed progressive remodeling of hematopoietic composition and morphology with disease severity (**Figure 2**). HSPCs (CD34+CD117-cells; see **Supp. Methods**), myeloblasts (CD34+CD117+ cells), and megakaryocytes increased commensurate with higher-risk disease (**Figure 2a**, **Supp. Figure 3a**), consistent with prior aspirate-based studies^9^ but now contextualized within intact marrow architecture. Megakaryocyte expansion in MDS-IB was accompanied by enrichment of smaller, frequently mononuclear forms; quantitative assessment of nuclear number and cell area confirmed this shift (**Figure 2b–c**). CD34-expressing megakaryocytes were present in all groups but were most frequent in *TP53*-mutated MDS-IB samples (**Supp. Figure 3d**). Erythroid normoblasts demonstrated increased cell area in MDS relative to age-matched NBMs and CCUS (**Supp. Figure 3e**), whereas early erythroid and myeloid precursors exhibited genotype-dependent size variation, with the smallest proerythroblasts and promyelocytes observed in *TP53*-mutated cases (**Supp. Figure 3f**).

**Figure 3.**
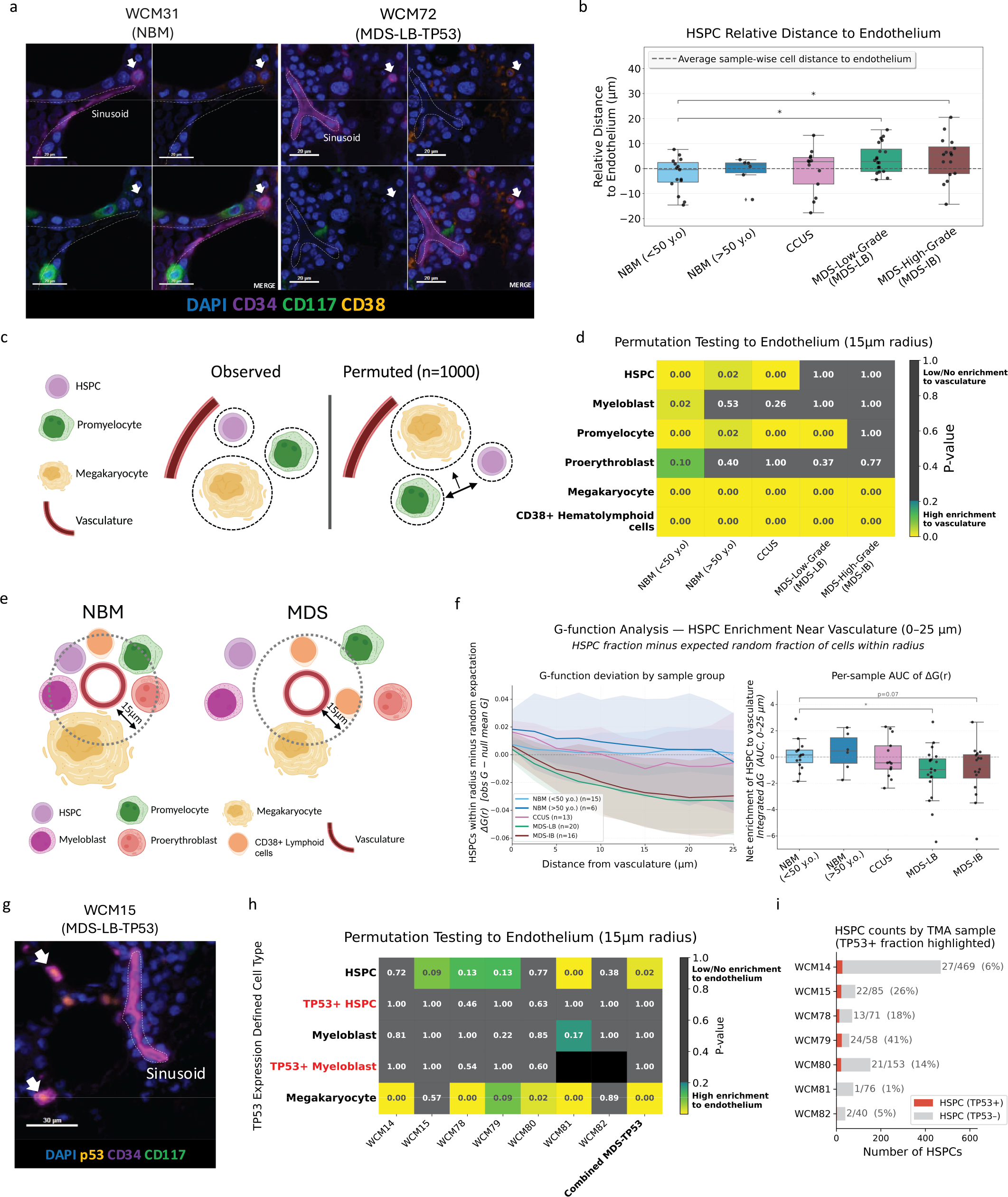
Reduced perivascular enrichment of hematopoietic stem and progenitor cells during progression to MDS. **(a)** Representative MxIF images showing HSPCs (arrows) adjacent to CD34⁺ endothelium in NBM and reduced perivascular localization in MDS (scale bars, 20 µm). **(b)** Boxplot comparing the relative distance of HSPCs to vasculature (defined as HSPC distance / all other cell types distance to vasculature in each sample) across groups. (Mann–Whitney U test; *P ≤ 0.05, **P ≤ 0.01). **(c)** Schematic illustrating the spatial analysis framework used to evaluate cellular enrichment near vascular structures. **(d)** Heatmap summarizing enrichment of indicated cell types within 15 µm of endothelium across disease groups. Significance was assessed using permutation testing with patient-level values combined across cases. Lower values (yellow) indicate significant enrichment relative to randomized spatial distributions. **(e)** Conceptual schematic summarizing alterations in perivascular niche composition between NBM and MDS. **(f)** Line plot and box plot of G function test of HSPC spatial enrichment to vasculature. Line plot value is difference of observed HSPC fraction within radius of vasculature, divided by a random expectation within each sample, as the mean for each sample group. Box plot value is AUC of this curve from radius of 0 to 25 µm representing net spatial enrichment (higher value is greater spatial enrichment to vasculature). **(g)** Representative *TP53*-mutated MDS sample showing HSPCs with elevated p53 expression (arrows) located distal to endothelium (scale bars, 30 µm). **(h)** Permutation analysis of vascular enrichment in the tissue microarray cohort stratified by p53 expression (i.e., *TP53* mutation status). Lower values indicate significant enrichment within 15 µm of endothelium. **(i)** Bar plot quantifying total number of HSPCs and *TP53-*mutated HSPCs in each tissue microarray sample

Genotype-specific remodeling patterns were evident, with *TP53*-mutated MDS exhibiting distinctive alterations in progenitor composition and erythroid differentiation. *TP53*-mutated MDS samples displayed marked depletion of promyelocytes and MMCs, alongside expansion of proerythroblasts and maturing erythroid normoblasts relative to other MDS-IB samples (**Figure 2d**). p53 protein labeling has been well-described as a method for identifying mutant p53 protein-containing cells (putative *TP53*-mutated cells) in tissue sections.^10–12^ Integration of p53 protein labeling with MxIF enabled in situ identification of putative *TP53*-mutated cells, which were enriched within myeloblast and erythroid precursor compartments (**Figure 2e–g**, **Supp. Figure 3c**). In contrast, endothelial cell proportions did not differ significantly across disease states (**Supp. Figure 3b**).

Collectively, these findings demonstrate reproducible, genotype-imprinted alterations in marrow cellular composition and morphology that extend beyond conventional blast enumeration and risk classification.

### HSPCs and early hematopoietic progenitors are progressively displaced from perivascular and para-trabecular zones along progression from clonal hematopoiesis to overt MDS

In normal bone marrow, hematopoietic stem and progenitor cells (HSPCs) localize preferentially to perivascular and para-trabecular niches.^8^ Consistent with our prior findings, HSPCs in young and older NBMs were significantly enriched within 15µm of CD34+ endothelial-lined vasculature (**Figure 3a–f**, **Supp. Figure 4a–b**). This enrichment pattern was largely preserved in CCUS samples. In contrast, HSPCs in both MDS-LB and MDS-IB samples were significantly displaced from vasculature, exhibiting increased average distances relative to NBMs and precursor states (**Figure 3a-f**, **Supp. Figure 4a–c**). Within MDS samples, HSPCs were positioned farther from vasculature than most other hematopoietic cell types (**Supp. Figure 4a**), indicating a selective disruption of the normal perivascular niche. These findings suggest that MDS is accompanied by disruption of canonical vascular niche interactions that govern normal HSPC localization.

**Figure 4.**
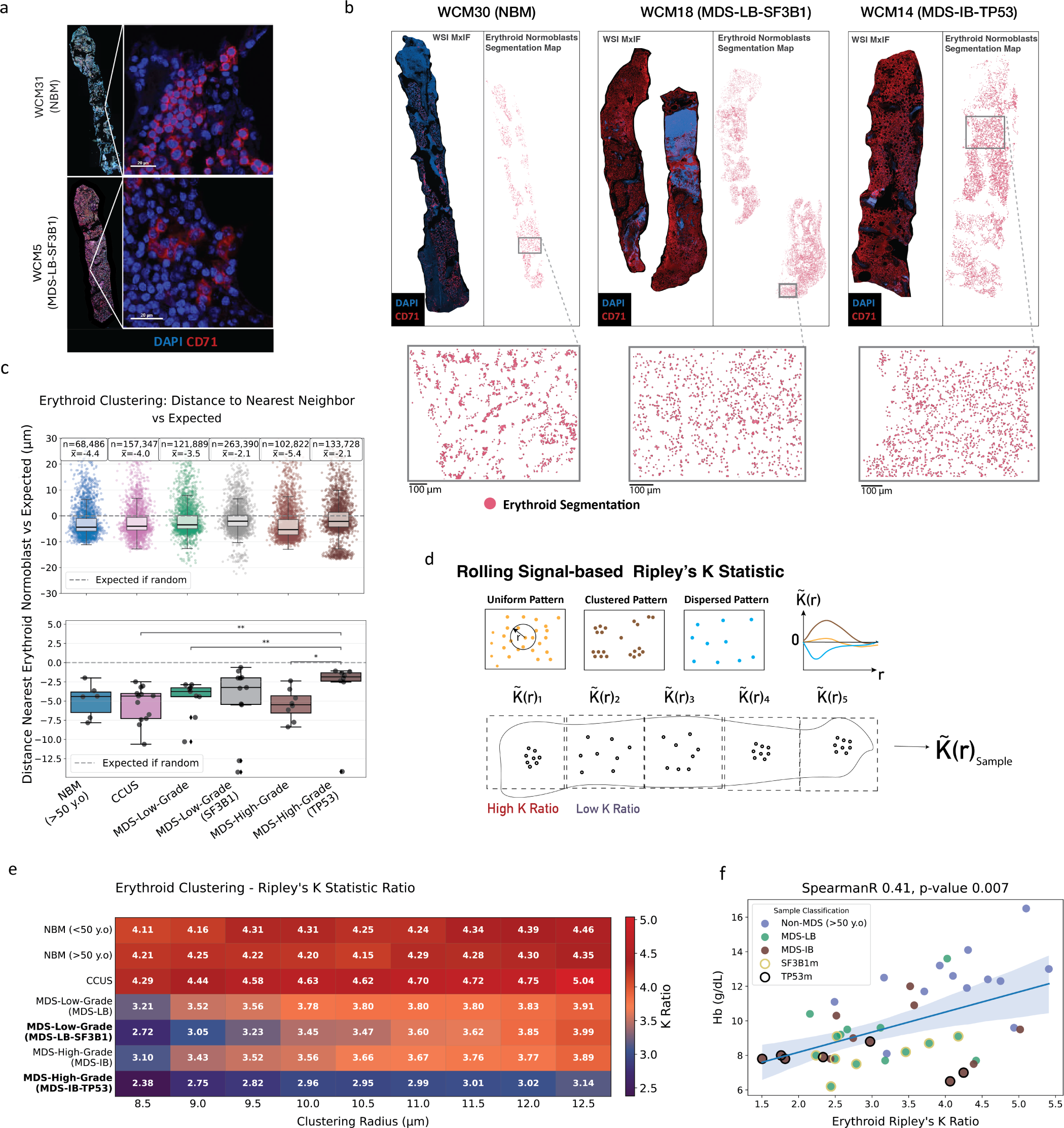
Disruption of erythroid island architecture in MDS. **(a)** Representative MxIF images showing a compact erythroid island in NBM (top) and dispersed erythroid normoblasts in an *SF3B1*-mutated MDS-LB sample (bottom) (scale bars, 20 µm). **(b)** Whole-slide examples showing merged DAPI and CD71 channels (left) and corresponding erythroid segmentation maps (right) for representative NBM, MDS-LB–*SF3B1*, and MDS-IB–*TP53* samples. Insets highlight local erythroid organization. **(c)** Nearest-neighbor distances among erythroid normoblasts normalized to expected distances relative to other cell types. Top, single-cell distributions; bottom, patient-level summaries (Mann–Whitney U test; *P ≤ 0.05, **P ≤ 0.01). **(d)** Schematic illustrating application of Rolling Signal–based Ripley’s K analysis to quantify erythroid clustering across spatial scales. **(e)** Mean Ripley’s K values across genotype-defined groups evaluated over radii of 8.5–12.5 µm. Lower values indicate reduced clustering. **(f)** Relationship between erythroid clustering (mean Ripley’s K) and hemoglobin concentration across MDS and age-matched controls (Spearman r = 0.41, P = 0.007).

A similar exclusionary pattern was observed for myeloblasts, while promyelocytes demonstrated significant displacement primarily in MDS-IB (**Figure 3d**, **Supp. Figure 4a-c**). In contrast, CD38+ small hematolymphoid cells and megakaryocytes remained consistently enriched near vasculature across disease states, with megakaryocytes showing relative enrichment in MDS samples (**Figure 3d**, **Supp. Figure 4a–c**). These findings indicate remodeling of perivascular cellular composition in MDS.

To determine whether clonal genotype influenced spatial localization, we leveraged p53 protein labeling to distinguish putative *TP53*-mutated from wild-type (WT) HSPCs. WT-HSPCs remained relatively proximal to endothelium, whereas *TP53*-mutated HSPCs were significantly displaced (**Figure 3g–i**, **Supp. Figure 4f**). This genotype-specific pattern suggests that clonal HSPCs contribute disproportionately to perivascular niche disruption in *TP53*-mutated disease.

Given the established role of CXCL12–CXCR4 signaling in HSPC homing to the vascular niche,^13^ we examined publicly available transcriptomic and proteomic datasets of MDS samples.^14–17^ Re-analysis revealed consistently reduced CXCR4 expression within HSPC populations from MDS patients relative to healthy controls (**Supp. Figure 5**). Across four independent datasets (3 transcriptomic, 1 proteomic), *CXCR4* was the only gene or protein significantly and reproducibly downregulated in HSPC compartments. These findings provide a potential molecular correlate for impaired perivascular localization in MDS.

**Figure 5.**
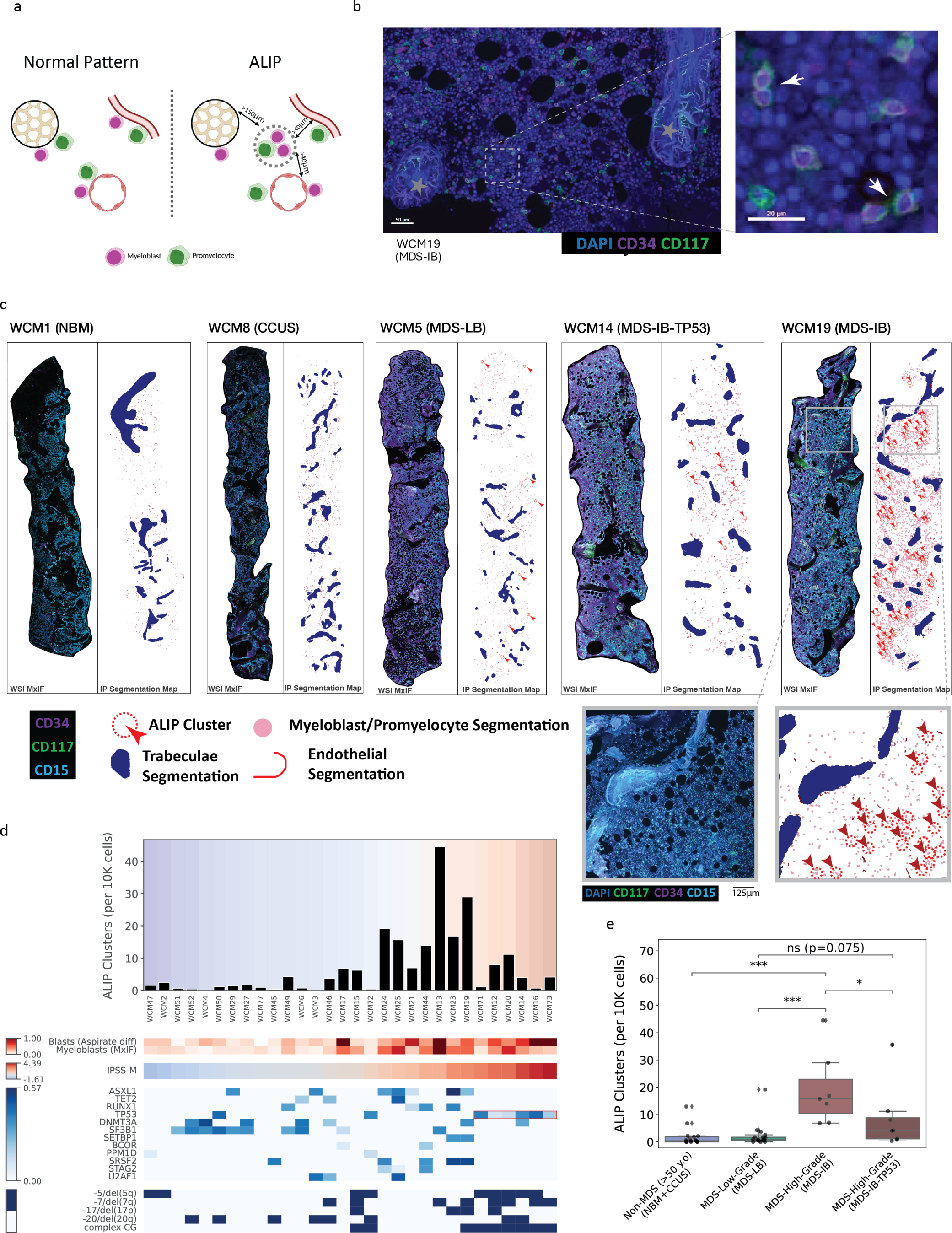
Atypical localization of immature precursors (ALIP) associates with disease severity and distinguishes TP53-mutated high-risk MDS. **(a)** Schematic illustrating computational identification of ALIP clusters, defined as ≥2 myeloblasts or promyelocytes located >150 µm from trabeculae and >40 µm from vasculature. **(b)** Representative MxIF image from an MDS-IB sample showing clusters of immature precursors distant from bone and endothelium (scale bar, 50 µm). Insets highlight ALIP clusters (scale bar, 20 µm). **(c)** Whole-slide examples showing merged CD117/CD34/CD15 channels and corresponding segmentation maps. Trabeculae, endothelium, and immature precursors are indicated. Representative cases include NBM, CCUS, MDS-LB, MDS-IB–*TP53*, and MDS-IB–*TP53*-WT. **(d)** Sample-level ALIP frequency normalized per 10,000 nucleated cells. Heatmaps show corresponding blast percentage, MxIF-derived myeloblast proportion, IPSS-M risk class, selected mutation variant allele frequency, and cytogenetic abnormalities. **(e)** ALIP cluster frequency across disease groups demonstrating highest levels in *TP53*-wild-type MDS-IB samples (Mann–Whitney U test; *P ≤ 0.05, **P ≤ 0.01, ***P ≤ 0.001).

In contrast to vascular displacement, HSPC enrichment near bone trabeculae was largely preserved across disease states, although average distances increased modestly in MDS-LB and MDS-IB samples (**Supp. Figure 4d–e**). Promyelocytes and myeloblasts, however, exhibited more widespread disruption of normal para-trabecular localization in MDS (**Supp. Figure 4d–e**).

Collectively, these analyses demonstrate selective displacement of early hematopoietic progenitors from perivascular niches in MDS, with pronounced effects in *TP53*-mutated disease and accompanying alterations in *CXCR4* expression, highlighting spatial remodeling of key regulatory microenvironments.

### Atypical erythroid island formation in MDS

Erythroid normoblasts in normal bone marrow are organized into well-formed maturation “islands,” a structural feature readily appreciated by light microscopy. Disaggregation of these islands has been described in advanced MDS,^18–21^ but systematic, quantitative assessment of erythroid spatial organization and its relationship to genotype and clinical parameters has not been previously performed.

Whole-slide erythroid phenotype mapping revealed large, cohesive erythroid aggregates in NBMs, with progressive disaggregation in MDS samples, particularly in MDS-IB (**Figure 4a–b**, **Supp. Figure 6a**). At the single-cell level, erythroid normoblasts in MDS were separated by significantly greater nearest-neighbor distances compared with NBMs and CCUS samples (**Figure 4c**), indicating reduced clustering.

**Figure 6.**
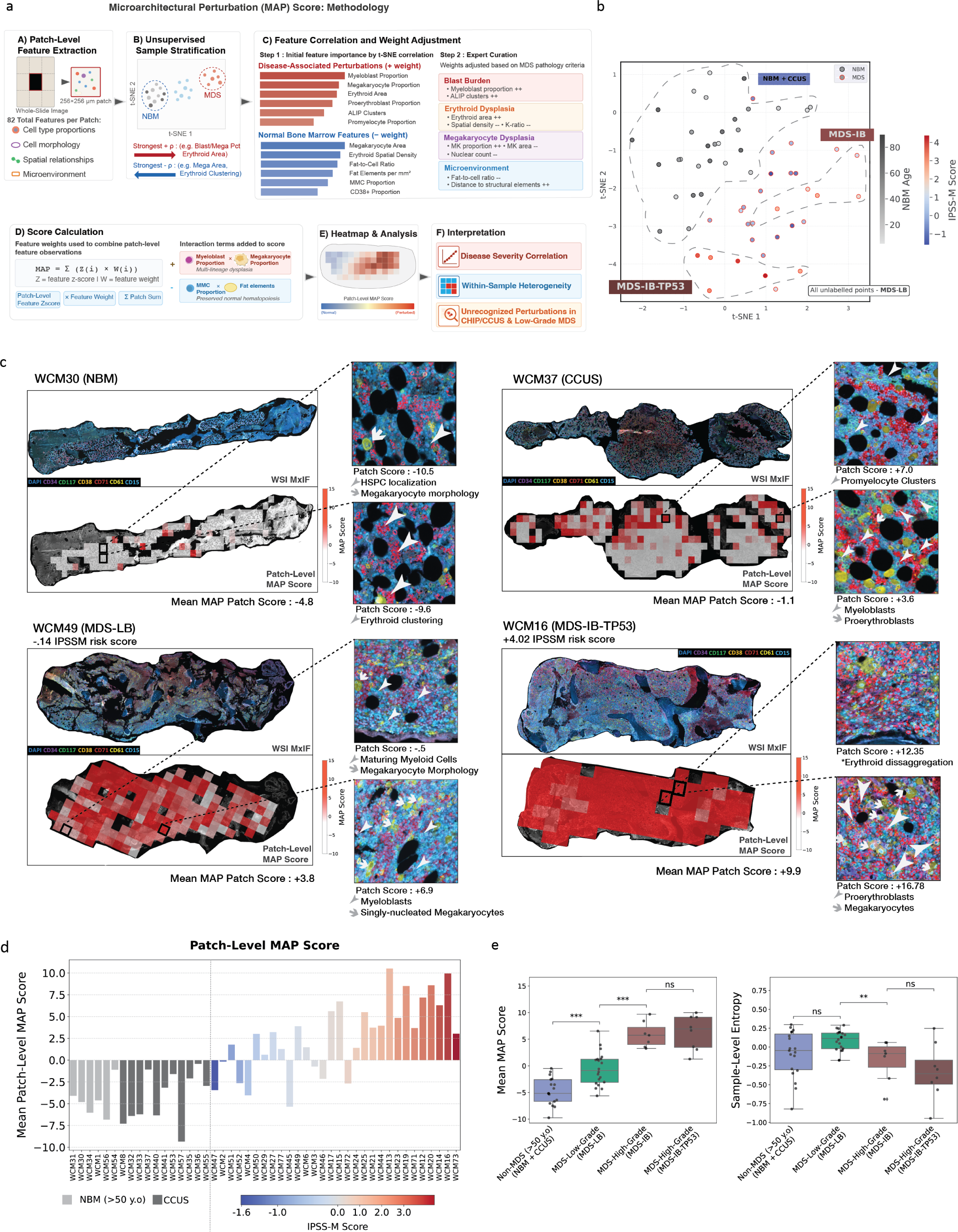
Derivation and spatial visualization of the MDS Microarchitectural Perturbation Score (MDS-MAPS). **(a)** Overview of MAPS derivation. Whole-slide images were divided into 256 × 256 µm patches, from which 82 predefined cellular and spatial features were extracted and used to develop a weighted composite score. **(b)** t-distributed stochastic neighbor embedding (t-SNE) of sample-level feature vectors showing separation of MDS samples from NBM and CCUS. NBM samples are color-scaled by age; MDS samples are color-scaled by IPSS-M score. **(c)** Representative whole-slide MxIF images and corresponding patch-level MAPS heatmaps for NBM, CCUS, MDS-LB, and *TP53*-mutated MDS-IB samples. Red indicates higher perturbation scores. Insets highlight representative patches and features contributing to elevated MAPS. **(d)** Mean MAPS values across the cohort. NBM and CCUS samples are shown in gray; MDS samples are colored by IPSS-M risk. **(e)** Comparison of mean MAPS and normalized entropy of patch-level MAPS distributions, reflecting spatial heterogeneity (Mann–Whitney U test; *P ≤ 0.05, **P ≤ 0.01, ***P ≤ 0.001).

To quantify erythroid spatial organization across treatment-naïve samples, we applied a Rolling Signal–based Ripley’s K analysis, which measures the degree of spatial clustering relative to a random distribution at progressively increasing radii. This approach captures both the magnitude of erythroid clustering (reflecting intact erythroid island formation) and its regional heterogeneity across the biopsy section (**Figure 4d**). NBMs exhibited the highest K-ratios, reflecting robust clustering, whereas the lowest K-ratios were observed in *TP53*-mutated MDS-IB samples (**Figure 4e**). Notably, *SF3B1*-mutated MDS-LB samples also demonstrated reduced clustering relative to mutation-negative MDS-LB and MDS-IB cases, indicating genotype-specific distortion of erythroid topology independent of blast burden. Erythroid cluster density analyses yielded concordant results (**Supp. Figure 6b–c**).

Importantly, the K-ratio correlated positively with peripheral blood hemoglobin levels (**Figure 4f**), linking quantitative disruption of erythroid island architecture with clinical anemia severity. Together, these findings demonstrate that erythroid island disaggregation is a measurable and genotype-imprinted feature of MDS microarchitecture and that quantitative spatial metrics capture clinically relevant aspects of ineffective hematopoiesis not discernible by conventional morphology alone.

### Atypical localization of immature precursors (ALIP) correlates with disease severity and discriminates TP53-mutated and -WT subtypes of high-risk MDS

In normal bone marrow, immature granulocytic precursors localize preferentially near bone trabeculae and vasculature, with progressive maturation occurring toward medullary regions. Atypical localization of immature precursors (ALIP) describes the abnormal clustering of myeloblasts and promyelocytes distant from these regulatory niches and has historically been associated with adverse cytogenetic features and inferior outcomes.^22,23^ However, ALIP has largely been assessed qualitatively, without systematic whole-slide quantification.

Using multiplex imaging and spatial mapping, we identified discrete foci of ALIP in a subset of MDS samples (**Figure 5a–b**). We operationally defined ALIP as clusters of two or more myeloblasts and/or promyelocytes within 12µm of one another located >150µm from trabeculae and >40µm from vasculature. Whole-slide analyses revealed marked variation in ALIP frequency and distribution across NBMs, CCUS, and MDS samples (**Figure 5c**).

Among treatment-naïve MDS cases, ALIP cluster density increased with disease severity, with the highest densities observed in many MDS-IB samples (**Figure 5d–e**, **Supp. Figure 7**). Notably, within the MDS-IB group, *TP53*–wild-type higher-risk cases exhibited greater ALIP cluster density than *TP53*-mutated counterparts (**Figure 5d–e**), revealing spatial heterogeneity within clinically similar high-risk disease categories. These patterns were robust to correction for cellularity and alternative cluster definitions.

**Figure 7.**
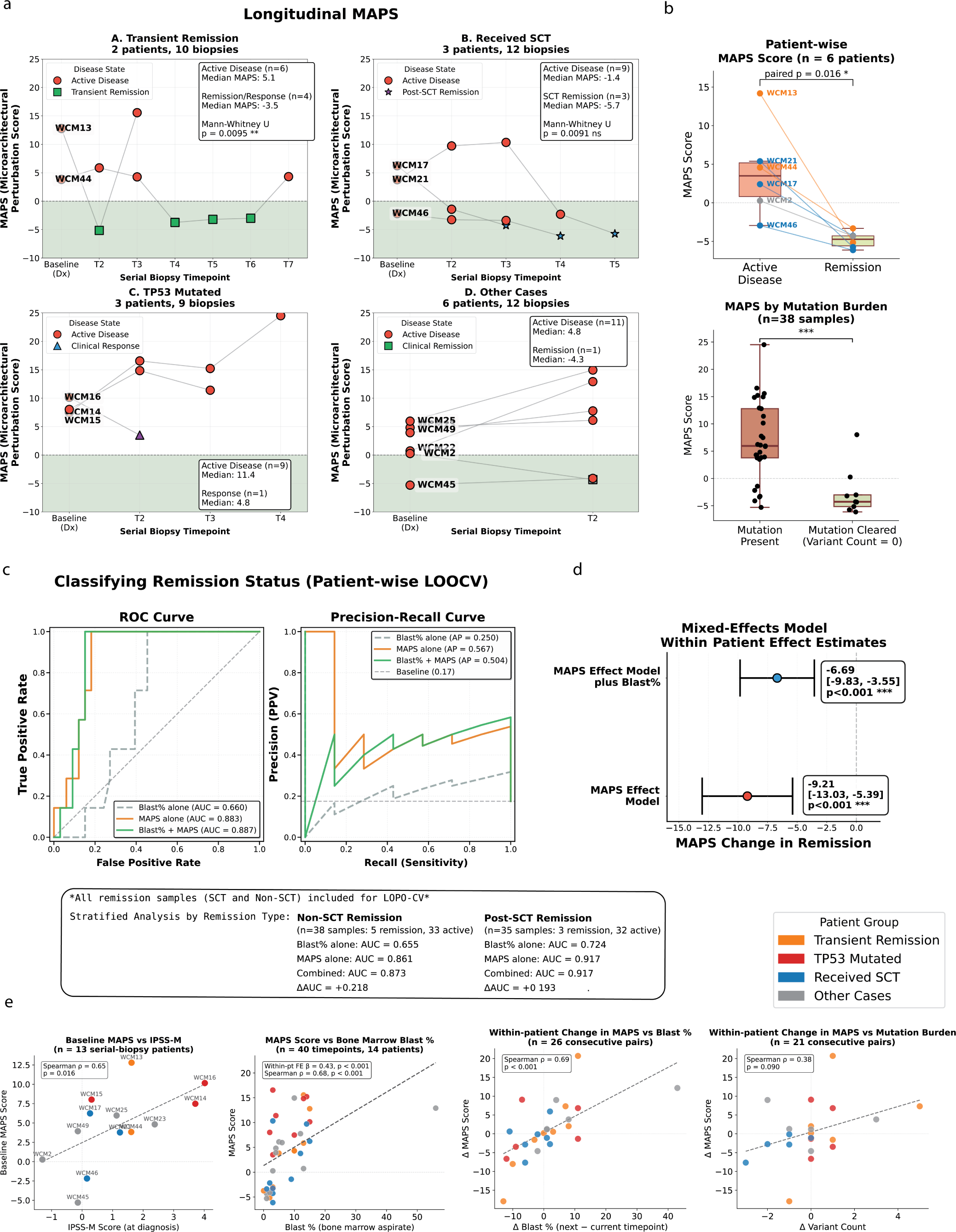
MDS-MAPS tracks therapeutic response and discriminates remission. **(a)** Longitudinal MAPS trajectories in patients with serial biopsies. Lines connect sequential samples; dashed line indicates MAPS = 0. Transient remission cases (WCM13, WCM44; n = 10 biopsies) showed MAPS reduction at remission (median 5.1 vs −3.5, P = 0.0095). Post-transplant remission cases (WCM17, WCM21, WCM46; n = 12 biopsies) showed similar decreases (median −1.4 vs −5.7, P = 0.0091). *TP53*-mutated cases (WCM14–16; n = 9 biopsies) exhibited persistently elevated MAPS. **(b)** Paired within-patient comparison (n = 6) demonstrating lower MAPS in remission (median 5.5 vs −4.6; Wilcoxon P = 0.016). Mutation-cleared samples also exhibited lower MAPS than mutation-persistent samples (median −3.2 vs 7.5; ***P < 0.001). **(c)** Leave-one-patient-out cross-validation (LOPO-CV) for remission classification (40 samples, 14 patients). ROC AUC: blast% = 0.660, MAPS = 0.883, combined = 0.887. Precision–recall average precision: MAPS = 0.567, blast% = 0.250. **(d)** Linear mixed-effects modeling (43 biopsies, 14 patients) demonstrating independent association of MAPS with remission after adjustment for blast percentage (β = −6.69, 95% CI −9.83 to −3.55; P < 0.001). **(e)** MAPS correlated with baseline IPSS-M score (n = 13; ρ = 0.65, P = 0.016) and with blast percentage across serial samples (within-patient ρ = 0.43; between-patient ρ = 0.68; both P < 0.001). Changes in MAPS correlated with changes in blast percentage (ρ = 0.69, P < 0.001).

These findings demonstrate that whole-slide quantification of ALIP uncovers genetically imprinted spatial heterogeneity in MDS that may not be captured by conventional risk stratification frameworks, reinforcing the concept that microarchitectural organization provides complementary information to blast enumeration and clinical scoring systems.

### Integration of single cell quantitative features and microarchitectural patterns to generate a spatially informed Microarchitectural Perturbation Score (MAPS)

To integrate the diverse architectural perturbations observed across MDS samples into a unified representation of tissue state, we developed a composite metric termed the MDS Microarchitectural Perturbation Score (MDS-MAPS) (**Figure 6a**). MAPS is computed within 256μm x 256μm patch regions across tissue sections by the summation of locally observed spatial and cellular features in patch regions according to feature weights representing the degree of disease-associated architectural perturbation. To derive feature weights, all 82 features from treatment-naive samples were projected onto a t-SNE dimensionality reduction plot, which revealed clustering of MDS samples along progressive gradations of disease severity. Features used spanned four domains: (1) cell type frequencies; (2) cell morphology; (3) cell-to-structure spatial enrichment; (4) cell-to-cell spatial enrichment. Individual features were then ranked by their correlation with disease severity along both t-SNE dimensions. Expert curation refined these weights based on known MDS biology resulting in 29 weighted features **(Supp. Data Table 6**). 3 combinatorial parameters were added, scored by their co-occurrence in patch regions. Once weights were finalized based on the observations for treatment-naive samples, they were locked and applied without modification to all remaining samples, including remission cases, to assess whether the composite spatial score captured clinically meaningful changes in marrow architecture across disease states.

Integration of spatial features by t-SNE dimensionality reduction revealed clear separation between normal marrow and MDS samples (**Figure 6b, Supp. Figure 8a-c**). Within MDS cases, *TP53*-mutated samples formed a distinct cluster independent of blast category, whereas *TP53*–wild-type MDS-LB and MDS-IB samples distributed along a continuum corresponding broadly to IPSS-M–defined risk strata. Feature importance analyses identified contributions from multiple domains—including progenitor displacement, erythroid clustering disruption, megakaryocyte morphometrics, and ALIP density (**Supp. Figure 8d**).

Patch-level MAPS values were derived using the locked weights described above, enabling visualization of localized perturbations across whole-slide sections (**Figure 6c**). Heatmap overlays revealed marked intra-tissue heterogeneity in many MDS samples. Mean MDS-MAPS values increased with morphologic severity, with the highest scores observed in *TP53*-mutated cases (**Figure 6d–e, Supp. Figure 8e**). Notably, several patients categorized as lower risk by IPSS-M exhibited elevated MDS-MAPS values (WCM6, WCM27, WCM29, WCM49, WCM50, WCM51, WCM77), and these discordant samples displayed focal features typically associated with higher-risk disease. MDS-LB samples demonstrated the greatest entropy of patch-level MAPS values (**Figure 6e**), indicating substantial spatial heterogeneity at earlier disease stages.

In addition, focal regions with elevated MAPS were detected in selected CCUS samples. These localized perturbations exhibited architectural features characteristic of overt MDS, suggesting that quantitative spatial mapping may detect early microarchitectural remodeling not readily apparent by conventional histopathology. These observations remain hypothesis-generating and warrant prospective validation.

Notably, global MAPS values in CCUS samples were not significantly different from age-matched normal bone marrow controls, indicating that the presence of clonal hematopoiesis and cytopenias alone is insufficient to produce the diffuse architectural remodeling characteristic of overt MDS. This distinction suggests that MAPS captures spatial perturbations specific to the transition from CCUS to MDS rather than to clonal hematopoiesis itself. However, the detection of focal elevated-MAPS regions in several CCUS biopsies that were globally near-normal suggests that spatial remodeling may begin as a localized process — detectable only through patch-level analysis — before progressing to the tissue-wide disruption observed in established MDS.

Collectively, these analyses establish MDS-MAPS as a quantitative tissue-level metric that integrates cellular composition, morphology, and spatial topology to represent the degree and heterogeneity of marrow architectural perturbation across genetic and clinical strata.

To determine whether MDS-MAPS could discriminate precursor states from overt low-risk disease, we performed leave-one-patient-out cross-validation comparing treatment-naïve CCUS (n=13) and MDS-LB (n=20) samples. MDS-MAPS demonstrated strong discrimination (AUC=0.815; average precision=0.890), outperforming mutation burden–based metrics including maximum and summed variant allele frequency (**Supp. Figure 9**). These findings indicate that quantitative microarchitectural perturbation distinguishes CCUS from morphologically defined low-blast MDS and captures tissue-level disease features not reflected by clonal burden alone.

### Correlation of cell morphologic and spatial features with blast progression

Because blast percentage remains central to clinical prognostication models such as IPSS-R and IPSS-M,^4,24^ we examined whether quantitative spatial and morphologic parameters derived from multiplex imaging were associated with subsequent blast progression in the subset of diagnostic MDS samples from patients who later received disease-modifying therapies (n=16) (**Supp. Data Table 3**).

Using modeling approaches appropriate for limited sample sizes, we evaluated the relationship between extracted spatial features and subsequent blast count increase. Models incorporating multiplex imaging–derived parameters demonstrated improved discrimination of blast progression relative to models based solely on standard clinical risk variables (**Supp. Figure 10a**). While MDS-MAPS broadly tracked with IPSS-M scores, multiple samples exhibited discordant risk estimates between these frameworks (**Supp. Figure 10b**), suggesting that spatial architecture captures complementary dimensions of disease biology.

Among individual features, displacement of promyelocytes from perivascular niches emerged as one of the most strongly associated spatial parameters with blast progression, demonstrating greater rank-based significance than conventional clinical variables (**Supp. Figure 10c–d**). These findings indicate that microarchitectural remodeling—particularly disruption of early progenitor niche localization—may reflect dynamic disease biology relevant to blast evolution.

Given the limited cohort size, these analyses are exploratory and hypothesis-generating. However, they suggest that quantifiable spatial features derived from whole-slide imaging may provide additive prognostic information beyond conventional blast enumeration and established risk scoring systems, warranting prospective validation in larger cohorts.

### Multimodal longitudinal assessment in the course of disease modifying therapies

Longitudinal biopsies were analyzed from 14 MDS patients treated with growth factor therapy and/or hypomethylating agents, with or without venetoclax; three patients additionally underwent allo-HCT. Among pharmacologically treated patients, three achieved clinical remission (WCM2, WCM13, WCM44), and three additional patients achieved remission following transplantation (WCM17, WCM21, WCM46).

Across serial samples, MDS-MAPS values were consistently reduced in remission biopsies relative to active disease (**Figure 7a**). Paired within-patient analyses demonstrated significant decreases in MAPS at remission timepoints, including in cases with molecular remission (**Figure 7b**), indicating that architectural normalization accompanies clinical and genetic response.

To evaluate whether MAPS could discriminate remission status in unseen patients, we performed logistic regression with leave-one-patient-out cross-validation (LOPO-CV), thereby avoiding within-patient data leakage. Across serial samples, MAPS demonstrated superior discrimination of remission versus active disease compared with blast percentage, as measured by both area under the receiver operating characteristic curve (AUC) and average precision (AP) in precision–recall analysis (**Figure 7c**). Notably, inclusion of blast percentage did not materially improve model performance, suggesting that MAPS captures dominant and partially orthogonal information regarding disease state.

Mixed-effects modeling confirmed that MAPS reductions in remission remained statistically significant after adjustment for blast percentage as a covariate (**Figure 7d**), indicating that the architectural changes captured by MAPS are not fully explained by changes in blast count alone. While MAPS correlated with established clinical risk metrics, discordance in several samples underscores that it quantifies distinct aspects of marrow organization (**Figure 7e**).

Representative longitudinal trajectories from patients WCM13 and WCM44 illustrate dynamic remodeling of marrow architecture across treatment courses (**Supp. Figure 11a)**. Remission samples embedded closer to NBMs in reduced-dimensional feature space and returned toward an MDS-associated profile at relapse (**Supp. Figure 11b**). In patient WCM44, transient re-localization of HSPCs to perivascular niches was observed at remission (**Supp. Figure 11c**); remission in both patients was accompanied by normalization of ALIP density, erythroid clustering, and megakaryocyte morphometrics (**Supp. Figure 11d**).

Although limited by sample size, these findings provide proof-of-principle that quantitative spatial architecture represents a dynamic and clinically informative tissue-state variable that tracks therapeutic response and complements conventional blast-based assessment.

## DISCUSSION

The spatial microarchitecture of hematopoiesis in the bone marrow remains incompletely characterized, particularly in human disease states.^25^ While recent in situ imaging studies have delineated the spatial organization of murine hematopoiesis under steady-state and stress conditions,^26,27^ comparable resolution in the human system has been limited. Building on our prior work establishing a single-cell, whole-slide framework for mapping normal human marrow architecture,^8^ we applied this approach to MDS to test whether quantitative spatial organization encodes clinically meaningful disease states.

Our findings demonstrate that MDS is characterized by coordinated and genotype-imprinted architectural remodeling. Increased proportions of hematopoietic stem and progenitor cells (HSPCs), myeloblasts, and megakaryocytes intensified with disease severity, but these quantitative shifts were accompanied by spatial reorganization not captured by conventional morphology. *TP53*-mutated cases exhibited particularly distinct architectural profiles, including altered erythroid and granulocytic progenitor composition and pronounced megakaryocyte abnormalities. Megakaryocytes are recognized regulators of the marrow niche, influencing thrombopoiesis and HSPC maintenance through paracrine signaling,^28,29^ and the marked morphologic perturbations observed in *TP53*-mutated disease suggest genotype-specific niche remodeling. These observations suggest that *TP53*-mutated disease may reshape the marrow niche in ways distinct from other MDS subtypes.

A central architectural alteration in MDS was displacement of HSPCs from perivascular niches. In normal marrow, HSPCs preferentially localize near trabecular bone and CD34-positive endothelial structures, consistent with prior demonstrations of perisinusoidal stem cell enrichment.^30^ Endothelial cells play a critical role in regulating hematopoietic homeostasis through angiocrine signaling,^31–38^ and disruption of these interactions has been implicated in hematologic disease.^33,34^ In MDS, particularly in *TP53*-mutated cases, HSPCs and early progenitors were selectively displaced from vascular niches. Re-analysis of publicly available transcriptomic and proteomic datasets^14–17^ revealed reproducibly reduced CXCR4 expression in MDS HSPCs relative to healthy controls. Given established CXCL12–CXCR4–mediated niche retention mechanisms and reported modulation of CXCR4 by p53,^39^ these findings provide a plausible molecular correlate linking cell-intrinsic genetic alterations with large-scale spatial reorganization. Although causality cannot be inferred, the convergence of genotype, gene expression, and spatial displacement supports a model in which clonal evolution is accompanied by niche remodeling.

Quantitative spatial analysis also enabled systematic reassessment of historically described morphologic abnormalities in MDS. Disruption of erythroid islands and abnormal progenitor clustering have long been recognized features of dysplastic marrow,^18,22^ yet comprehensive spatial quantification has been lacking. We demonstrate that erythroid island disaggregation correlates with anemia severity and occurs in genotype-specific patterns, including prominent disruption in *SF3B1*-mutated low-blast cases. SF3B1 plays a critical role in erythropoiesis through regulation of mRNA splicing,^40^ and mutation-associated mis-splicing contributes to ineffective erythroid differentiation and anemia.^41,42^ Taken together, these observations suggest that spatial disorganization of erythropoiesis represents a structural manifestation of ineffective hematopoiesis in MDS. Similarly, quantitative assessment of atypical localization of immature precursors (ALIP) revealed spatial heterogeneity within clinically similar high-risk MDS cases, reinforcing the concept that marrow topology reflects underlying genetic programs.

Integration of cellular composition, morphologic features, and spatial topology into the MDS Microarchitectural Perturbation Score (MDS-MAPS) provided a unified representation of tissue-level disease state. While MDS-MAPS broadly aligned with IPSS-M categories, discordant cases—particularly among lower-risk patients—suggest that spatial architecture captures dimensions of disease biology not incorporated into current prognostic models. Patch-level analyses further revealed substantial intra-tissue heterogeneity, especially in low-blast MDS, indicating that early disease may involve focal architectural remodeling that could be overlooked in aspirate-based or purely morphologic assessments. Notably, in leave-one-patient-out cross-validation comparing treatment-naïve CCUS and MDS-LB samples, MDS-MAPS demonstrated strong discrimination and outperformed mutation burden–based metrics, including maximum and summed variant allele frequency. This finding suggests that quantitative microarchitectural perturbation distinguishes precursor states from morphologically defined low-blast MDS and captures tissue-level disease biology not reflected by clonal burden alone.

Longitudinal analyses underscore the dynamic nature of spatial remodeling. MDS-MAPS decreased consistently in remission samples and discriminated active disease from remission using leave-one-patient-out cross-validation, outperforming blast percentage in both ROC and precision–recall analyses. Mixed-effects modeling confirmed that MAPS reductions remained significant after adjustment for blast burden, supporting the interpretation that quantitative architecture represents an orthogonal tissue-state metric. In representative patients, normalization of HSPC localization and progenitor clustering accompanied clinical and genetic remission, with re-emergence of spatial perturbations at relapse. These findings provide proof-of-principle that intact marrow architecture tracks therapeutic response. While MAPS substantially outperformed blast percentage in classifying remission status, we note that the composite score incorporates features related to progenitor and blast-like populations as well as spatial parameters that may covary with blast expansion. Accordingly, the improved performance of MAPS relative to blast percentage alone likely reflects both the integration of non-blast spatial features — such as erythroid island organization, lineage distribution, and niche localization — and the richer quantification of blast-related biology afforded by spatial context (e.g., ALIP detection/quantification). Disentangling the mechanisms of blast-related versus non-blast spatial alterations across therapeutic trajectories is an important direction for future work.

Our findings highlight the importance of considering the bone marrow as a spatially organized tissue ecosystem in myeloid malignancies. While genomic alterations initiate clonal hematopoiesis, disease progression appears accompanied by coordinated remodeling of cellular composition, microenvironmental interactions, and niche localization. Quantitative spatial approaches may therefore provide complementary insight into disease biology beyond molecular profiling alone.

Exploratory analyses of blast progression further suggest that specific spatial parameters—particularly progenitor displacement from vascular niches—may provide additive prognostic information beyond conventional clinical variables. Although limited by cohort size, these observations support prospective evaluation of spatial biomarkers in larger, clinically annotated cohorts.

This study has limitations. The use of a 7-plex immunofluorescence panel constrains the granularity with which specific cell populations can be resolved. For example, megakaryocytes are identified using CD61 alone; incorporation of additional markers such as CD41 and CD42 would enable discrimination between megakaryocyte progenitors, immature forms, and mature cells. Similarly, CD71 identifies erythroid precursors broadly but does not resolve individual stages of erythroid differentiation. We note that our pipeline distinguishes CD34⁺/CD117⁻ cells (HSPCs) from CD34⁺/CD117⁺ cells (myeloblasts) by combinatorial phenotyping; nevertheless, the CD34⁺/CD117⁻ population likely includes both primitive stem cells and early committed progenitors not further resolved by this panel. However, the 7-plex design reflects a deliberate trade-off: by limiting the panel to markers compatible with standard pathology laboratory instrumentation and the Opal tyramide signal amplification platform, we prioritized whole-slide coverage of a large cohort and clinical scalability over molecular depth. Higher-plex spatial platforms — including CODEX, imaging mass cytometry, multiplexed ion beam imaging, COMET (SeqIF), and spatial transcriptomics — would enable finer resolution of differentiation states and more precise niche characterization, and application of such platforms to this cohort represents an important direction for future work. Importantly, despite these constraints, the 7-plex panel was sufficient to detect reproducible spatial perturbations that distinguished disease states and tracked treatment response, suggesting that the architectural features captured by MAPS reflect biologically meaningful organizational changes rather than artifacts of limited phenotypic resolution. In addition, the sample size, particularly for progression and remission analyses, constrains definitive prognostic conclusions, and mechanistic interpretations of niche disruption remain correlative. Validation in independent and prospectively collected cohorts will be essential to establish clinical utility. Nevertheless, interrogation of millions of spatially resolved cells across diagnostic and longitudinal biopsies provides a high-resolution framework for understanding marrow organization in MDS.

In summary, we demonstrate that quantitative spatial architecture of the human bone marrow encodes genotype, disease severity, and therapeutic response in MDS. By integrating multiplex imaging with computational modeling, we define a tissue-level metric that complements blast-based and molecular assessments. These findings support incorporation of spatially resolved analysis into future diagnostic and translational studies and provide a foundation for evaluating microarchitectural biomarkers in hematologic malignancies.

## DATA AVAILABILITY

All data and code will be deposited online (https://github.com/BoneMarrowMxIFImaging/MDS.Manuscript) following formal manuscript review.

## AUTHOR CONTRIBUTIONS

*Conceptualization*: S.P. *Methodology*: R.N., D.R., S.P. *Formal analysis*: R.N., D.R. *Investigation*: R.N., A.K., J.C., A.R., I.V., C.U., J.J., F.S., J.P., C.M., L.Z., D.S., G.R., P.D., J.K., J.F., M.G., N.L., A.C., M.O., P.S., J.G., G.I., S.R., D.R., S.P., *Visualization*: R.N. *Resources*: N.L., G.I., S.R., S.P. *Writing*: R.N., S.P. *Supervision*: D.R., S.P. *Funding acquisition*: S.R., S.P.

## ACKNOWLEDGEMENTS

This work was supported by the Department of Pathology and Laboratory Medicine, Weill Cornell Medical College (start-up funding to S.S.P.). Tissue staining and imaging was performed in the Multiparametric In Situ Imaging (MISI) Laboratory of the Department of Pathology and Laboratory Medicine, Weill Cornell Medical College.

