## Supplementary Materials for "Spatial remodeling of bone marrow architecture defines tissue-state signatures of disease activity and therapeutic response in myelodysplastic neoplasms"

### METHODS

#### *Clinicopathologic and Laboratory Data*

Electronic medical records were reviewed for all patients to extract clinical data concurrent with the evaluated biopsy, including time of collection, body mass index (BMI), complete blood count parameters (hemoglobin, mean corpuscular volume, absolute neutrophil count, platelet count), and marrow blast percentage. Diagnostic hematopathology reports were re-reviewed by board-certified hematopathologists (SSP, JTG) to confirm diagnostic classification (NBM, CCUS, MDS) and to extract aspirate differential counts, including myeloid-to-erythroid (M:E) ratio and lineage-specific proportions.

International Prognostic Scoring System–Molecular (IPSS-M) scores were calculated for all treatment-naïve MDS samples. Cytogenetic and molecular findings were incorporated into diagnostic categorization as described in the main Methods. Clinical and laboratory parameters, flow cytometry and NGS data, as well as longitudinal clinical/treatment response data are presented in **Supplemental Data Tables 1-5**.

#### *Study Cohort and Clinical Definitions*

The cohort comprised 49 unique patients with myelodysplastic neoplasms (MDS; n=36) or clonal cytopenia(s) of undetermined significance (n=13), contributing 77 trephine biopsy samples. Twenty-nine serial biopsies from 14 MDS patients (range 2–7 per patient) were obtained after the diagnostic biopsy. Twenty-one normal bone marrow (NBM) controls from a previously published cohort were included. A tissue microarray (TMA) incorporating 7 additional *TP53*-mutated MDS cases and 5 NBMs was included in analyses corresponding to main text Figures 2–3.

Clinical definitions were prespecified and adjudicated prior to development of the MDS Microarchitectural Perturbation Score (MDS-MAPS), and therefore without knowledge of MAPS results. Diagnostic (treatment-naïve) samples were those establishing diagnosis prior to therapy. Serial samples were those obtained after diagnosis. Disease-modifying therapies primarily included hypomethylating agents (azacytidine or decitabine) with or without venetoclax **(Supplemental Data Tables 1 and 3)**.

##### *Remission versus Active Disease Classification of Individual Biopsies*

Remission versus active disease status for serial biopsies was determined through integration of 2023 International Working Group (IWG) response criteria,[1] contemporaneous pathology reports, and concurrent cytogenetic and molecular findings. Biopsies were classified as remission if they met IWG criteria for complete remission (CR) or CR with incomplete hematologic recovery (CRi), including marrow blasts <5%, and were interpreted morphologically as remission without persistent dysplasia or excess blasts. Cases were classified as active disease if driver mutations and/or clonal cytogenetic abnormalities consistent with residual MDS persisted, even when blasts were <5%. Molecular remission was defined as absence of detectable mutations to a sensitivity of 5% variant allele frequency (VAF) using clinically validated non–minimal residual disease targeted sequencing. Blast progression was defined as any increase in marrow blast percentage relative to baseline. Clinical laboratory data were abstracted from medical records.

##### *Clinical Laboratory Characterization*

All diagnostic samples underwent standard clinical evaluation including histomorphologic review, multiparameter flow cytometry (MFC), cytogenetic analysis, and targeted next-generation sequencing (NGS).

MFC was performed using a 10-color EuroFlow[2] panel with acquisition of ≥200,000 events per sample and analyzed using BD FACSDiva. Blast populations were defined as CD34+/CD117+ cells. Flow-derived proportions were used for orthogonal validation of multiplex immunofluorescence–derived cell types.

Conventional cytogenetic analysis was performed on 20 metaphases per case and reported according to International System for Human Cytogenetic Nomenclature guidelines. FISH was performed when clinically indicated using probes targeting recurrent MDS-associated abnormalities.

Targeted NGS was performed using either a 45-gene panel (analytical sensitivity ~2% VAF for SNVs, 1% for indels) or the TruSight Oncology 500 assay (~3% VAF sensitivity) on Illumina platforms, with analysis using clinically validated pipelines. Mutation status was derived from diagnostic reports.

***Multiplex Immunofluorescence Tissue Staining***

Four-micron-thick Bouin-fixed, decalcified, paraffin-embedded trephine sections were prepared and stained using the Opal multiplex immunofluorescence system (Akoya Biosciences) on a Bond RX automated stainer (Leica Biosystems).[3] Slides were deparaffinized and subjected to EDTA-based antigen retrieval (Leica ER2, 20 minutes). Cyclical horseradish peroxidase–mediated tyramide signal amplification was performed, with antibody stripping between cycles using citrate-based retrieval (Leica ER1).

The primary antibody panel (applied sequentially) included:

1. CD71 (Opal 480; clone 10F11; 1:80)
2. CD61 (Opal 520; clone 2F2; ready-to-use)
3. CD117 (Opal 570; clone D3W6Y; 1:100)
4. CD38 (Opal 620; clone 38C03; 1:50)
5. CD34 (Opal 690; clone QBEND/10; 1:100)
6. CD15 (Opal 780; clone MMA; 1:50)

For tissue microarray (TMA) analyses, CD38 was replaced with p53 (clone DO-7) [examples below].

<u>Representative single channel and merge immunofluorescence images from validation of p53-containing panel applied to TMA</u>. Scale bars shown are 100µm (tonsil, 11.2x magnification), and both 100µm (top row, 11.2x magnification) and 20µm (bottom row, 48.2x magnification) for all other case examples.

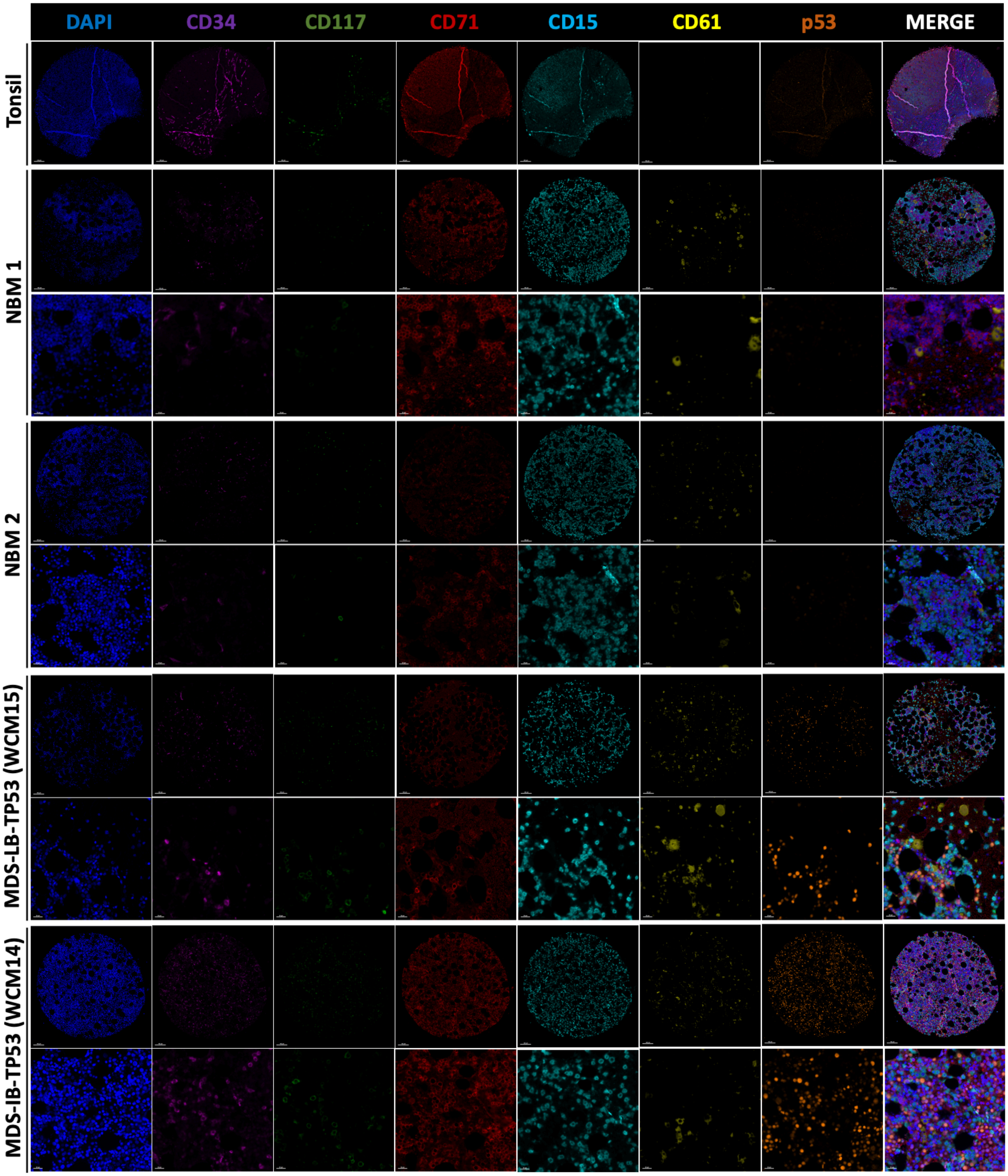

For both the stained whole trephine biopsy sections and the TMA slides, whole-slide images (WSIs) were acquired at 20× magnification using the Vectra Polaris / PhenoImager platform (Akoya Biosciences). Spectral unmixing was performed in InForm (v2.4.8), tiles were fused in HALO (v3.3.2541.231), and multispectral image stacks were exported as multi-layer TIFF files for computational analysis.

##### *Image Processing and Segmentation Pipeline*

Image analysis was performed in Python (v3.11) using open-source libraries.

Whole-cell and nuclear segmentation was performed using Mesmer (DeepCell v0.12.9),[4] a convolutional neural network–based segmentation framework optimized for multiplex immunofluorescence images. The Mesmer architecture consists of a feature pyramid network coupled to a ResNet50 backbone, with dual semantic heads that predict inner-distance transforms and pixel-wise classification maps to delineate individual cells.

The model was pretrained on an augmented TissueNet dataset and subsequently fine-tuned using manually annotated 256 × 256 pixel bone marrow image patches derived from this cohort. Fine-tuning employed the default Mesmer parameter configuration with a four-channel semantic head output (inner distance, outer distance, foreground–background classification, and pixel-wise transformations).

Segmentation employed a watershed-based postprocessing pipeline with an interior probability threshold of 0.3 and a centroid threshold of 0.1. A deep watershed transform was applied to extract cell masks, using parameters maxima_threshold = 1, interior_threshold = 0.3, and interior_smoothing = 1. The minimum nuclear and whole-cell area was set to 15 pixels, with no imposed maximum area constraint. Edge-touching cells were retained and not excluded from downstream analyses.

Whole-slide segmentation was performed by tiling images into overlapping 256 × 256 pixel patches. Segmentation masks were subsequently merged across tile boundaries using a Dask-image–based patch stitching algorithm to generate continuous whole-slide masks. Segmentation accuracy was validated against manually annotated masks, with performance metrics reported in **Supplementary Table 5**.

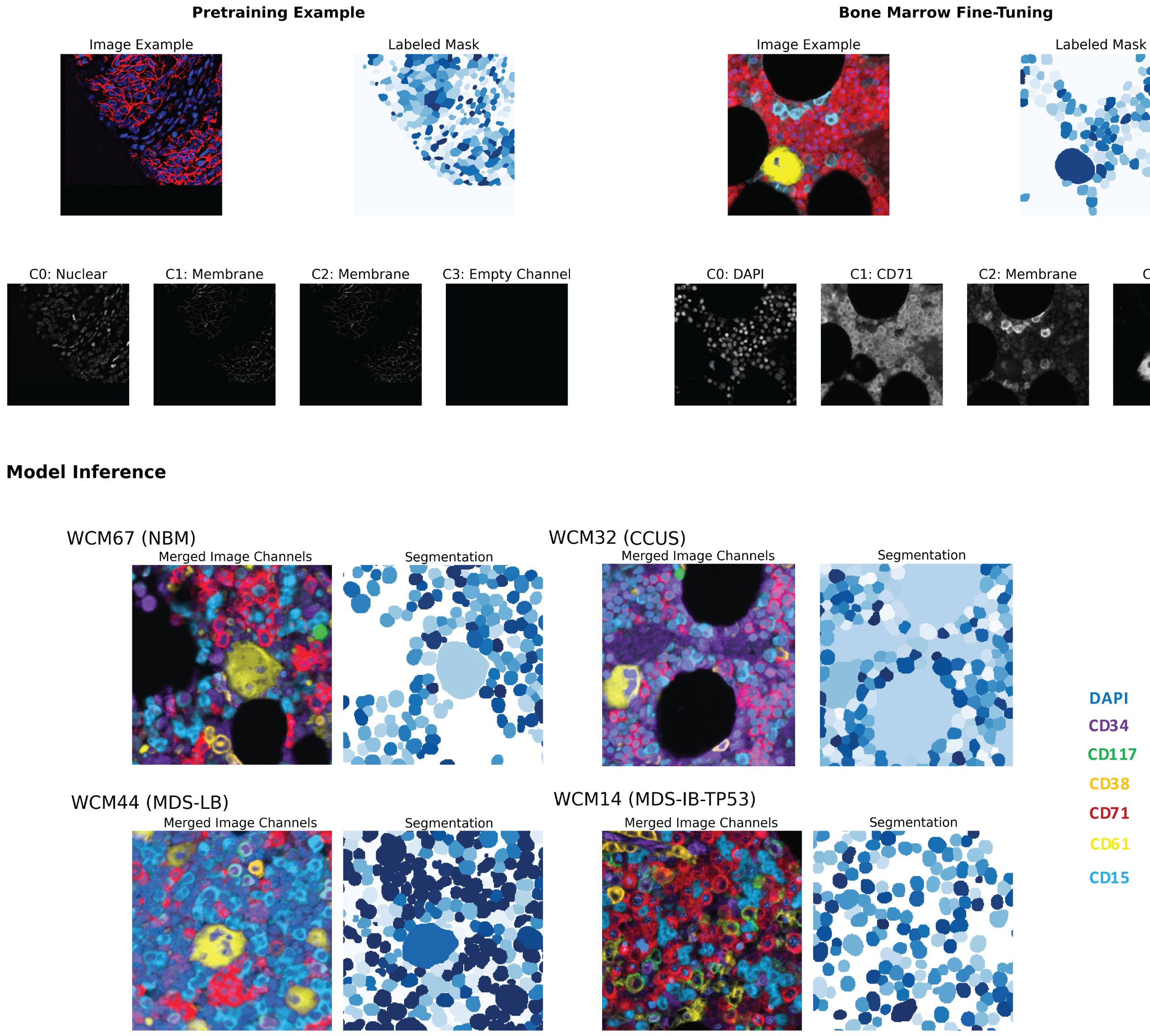

##### *Cell Phenotyping and Classification*

Single-cell membrane marker classification was performed using a ResNet18 convolutional neural network initialized with ImageNet pretrained weights. The neural network was fine-tuned on a dataset of expert hematopathologist annotations, including examples of all 6 membrane markers, which is detailed in Supplementary Table 5. For each segmented cell, the classifier received a three-channel cropped image centered on the cell of interest, consisting of the binary whole-cell segmentation mask, the DAPI nuclear channel, and the individual membrane marker channel being evaluated. Marker positivity classifications were validated against expert hematopathologist annotations on a held-out set of image examples, with performance metrics reported in Supplemental Table 5. The classifier was applied separately to each membrane marker. .

Each classifier generated a continuous probability score (range 0–1) representing the likelihood of marker positivity. Marker intensities were normalized using a slide-wise normalization approach prior to classification. Positivity thresholds were determined using manual gating aligned with expert hematopathologist review. Unless otherwise specified, cells were considered positive for a given marker at a probability threshold of >0.1. For CD34 and CD61, more stringent thresholds were applied (>0.25 probability) to minimize false-positive assignments in structurally complex regions.

Following marker-level classification, cell identities were assigned using a vector-based approach. For each cell, the predicted marker probability vector was compared to predefined lineage signature vectors using cosine similarity, and the phenotype corresponding to the closest matching signature was assigned as the final cell identity. The lineage signature vectors were constructed based on canonical immunophenotypic profiles established in clinical hematopathology, with each vector specifying a binary pattern of expected marker positivity and negativity for a given cell type (e.g., myeloblast [$CD34^+/CD117^+$]: [1,1,0,0,0,0]; proerythroblast [$CD117^+/CD71^+$]: [0,1,0,1,0,0]). Cosine similarity was selected for vector matching because it emphasizes the pattern of marker expression rather than absolute classifier confidence magnitude, providing robustness to inter-slide staining intensity variation inherent in multiplex immunofluorescence imaging.

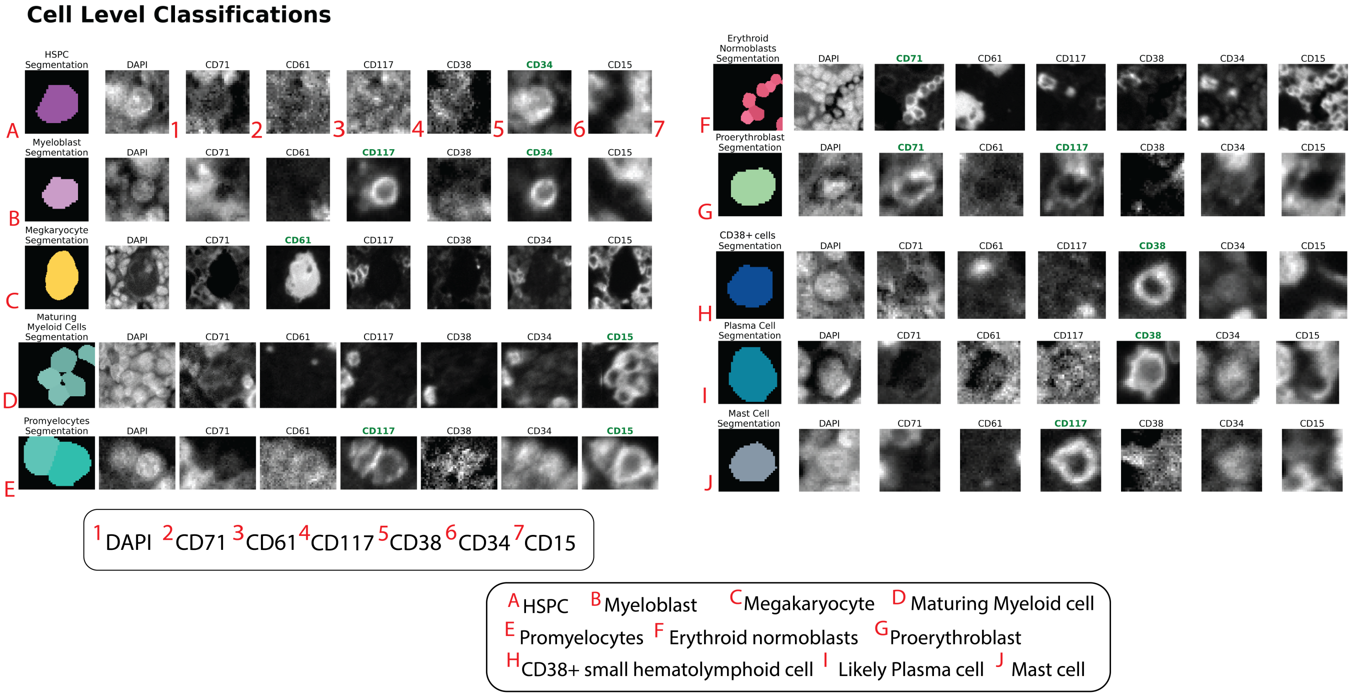

**Phenotyping Maps**

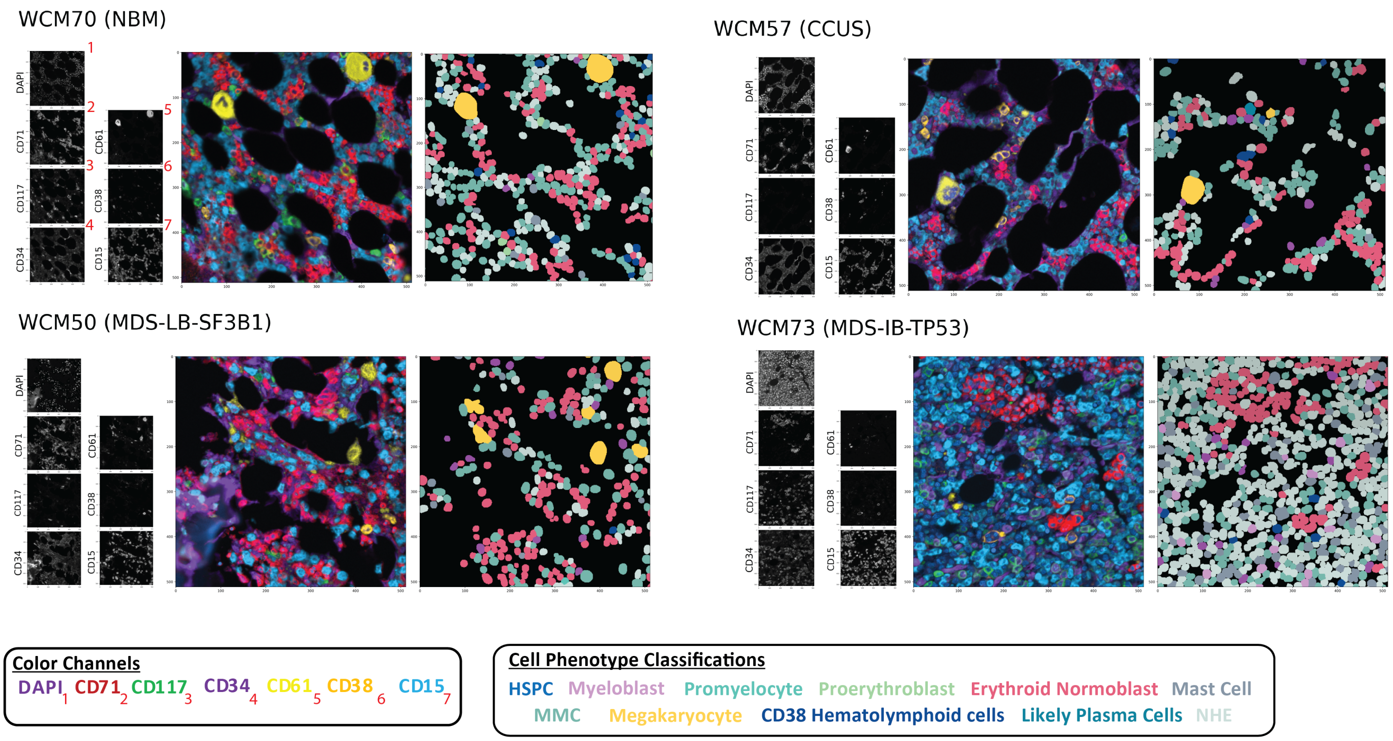

***Discrimination of CD34+ hematopoietic cells from CD34+ endothelial cells***

Endothelial cells and CD34⁺ hematopoietic cells are distinguished in our pipeline through two complementary approaches. First, sinusoidal vasculature is identified using a dedicated structural segmentation mask that detects vessel-like structures based on morphologic features (elongated, branching architecture) distinct from round hematopoietic cells. Second, CD34 mean fluorescence intensity (MFI) is used as a discriminating feature: endothelial cells exhibit higher CD34 MFI than hematopoietic progenitors. Specifically, $CD34^+$ nuclei with >50% spatial overlap with the vascular segmentation mask are classified as endothelial-associated and excluded from hematopoietic cell quantification; remaining $CD34^+$ cells are available for hematopoietic cell type classification. This dual criterion — morphology-based structural masking combined with intensity-based and spatial overlap thresholding — minimizes misclassification between the two $CD34^+$ populations.

***p53 Classification (TMA)***

To enable in situ identification of putative TP53-mutated cells, a separate convolutional neural network classifier was developed to categorize nuclear p53 protein expression levels. A ResNet18 architecture initialized with ImageNet-pretrained weights was trained to classify individual nuclei into three expression categories: no expression, low expression, and moderate/high expression. Cells classified as moderate/high were operationally designated as p53-positive for downstream spatial analyses.

A total of 76 manually annotated single-cell nuclear p53 images were used for model development. Annotations were performed by expert review and stratified into the three expression categories described above. Images were randomly partitioned into training (80%) and validation (20%) sets, with 25 annotated single-cell images used for validation.

On the held-out validation set, the classifier achieved an overall accuracy of 0.95, precision of 1.00, recall of 0.86, and an F1 score of 0.92 for identification of moderate/high p53 expression. These performance metrics supported the use of the model for in situ detection of p53-overexpressing cells within multiplex immunofluorescence datasets.

A comprehensive summary of model architecture, preprocessing steps, training parameters, and validation procedures is provided in **Supplementary Table 5**.

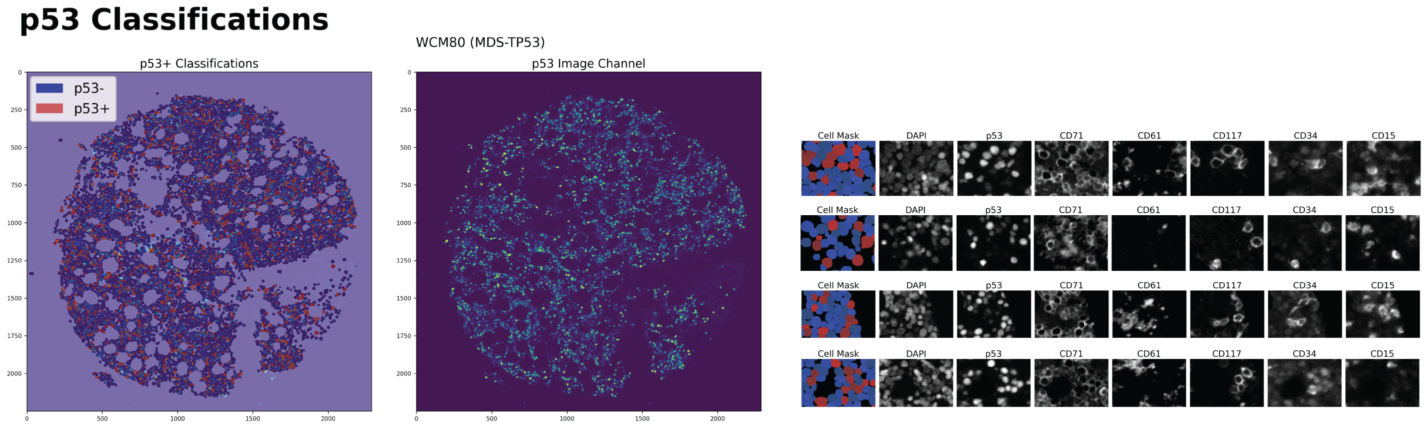

##### *Structural Masks Segmentation*

CNN-based segmentation was used to generate binary masks for endothelium (CD34+ elongated structures), bone trabeculae, and adipose tissue, as described in Sarachakov et al. Cell-to-structure distances were calculated by subtracting estimated cell radius from centroid-to-boundary distance using OpenCV functions (findContours, pointPolygonTest).

##### *Morphologic Feature Extraction*

Morphologic features were computed directly from the whole-cell and nuclear segmentation masks generated during image processing. For each segmented cell, total cell area was calculated by converting the pixel area of the whole-cell mask into square microns ($\mu m^2$) using the known image resolution. Nuclear area was similarly computed from the corresponding nuclear mask.

Cell shape was quantified using eccentricity, defined as the ratio of the major axis to the minor axis of an ellipse fitted to the segmentation mask, providing a measure of deviation from circularity. For megakaryocytes, multinucleation was quantified by counting the number of discrete nuclear mask objects contained within the corresponding whole-cell boundary.

All morphologic computations, including area measurements, ellipse fitting, and nuclear enumeration, were performed using the OpenCV Python library.

#### Spatial Statistics Implementation

##### *Permutation Testing*

Spatial enrichment analyses were performed using a permutation-based framework to assess whether specific cell phenotypes were preferentially localized relative to structural elements. For each whole-slide image, cell phenotype labels were randomly permuted 1,000 times while preserving the original spatial coordinates, thereby generating a null distribution for enrichment statistics.

Distances between cells and structural elements were calculated using the Euclidean distance metric in two-dimensional space. Enrichment was evaluated within predefined radii of 15 μm for vascular proximity and 30 μm for trabecular proximity, selected based on the spatial scale of these structural components within bone marrow tissue.

P-values were calculated as the proportion of permuted datasets demonstrating equal or greater enrichment than observed. No Benjamini–Hochberg false discovery rate correction was applied to permutation-derived p-values in these analyses.

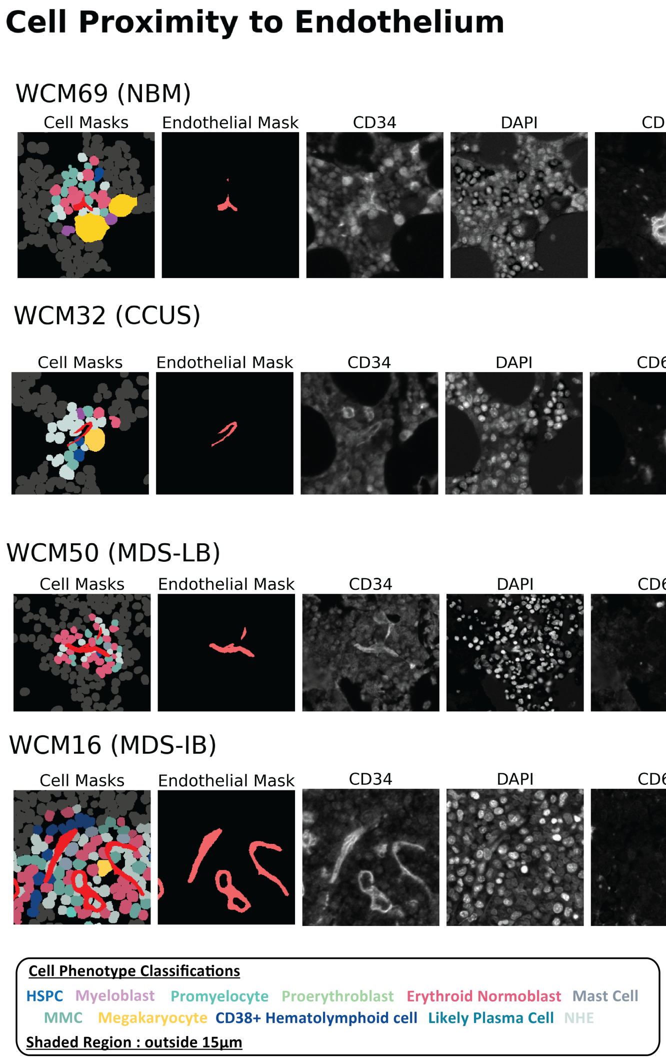

##### *Ripley's K (Rolling Signal)*

Spatial clustering of erythroid normoblasts was quantified using a rolling-window implementation of Ripley's K statistic.[5] Whole-slide images were subdivided into overlapping windows measuring 256 × 256 µm, with 50% overlap between adjacent windows to ensure continuous spatial coverage while preserving local resolution.

Within each window, Ripley's K was computed across a range of radii spanning 8.5 to 12.5 µm in 0.5 µm increments, capturing clustering behavior at distances relevant to erythroid island organization. A border edge-correction method was applied to account for truncated neighborhoods at window boundaries.

Whole-slide K values were calculated as the mean of the window-level K estimates, providing a summary measure of erythroid clustering across the entire tissue section.

##### *HDBSCAN*

Erythroid clustering used HDBSCAN, with min_cluster_size = 5, min_samples = 3, cluster detection method = eom, and Euclidean values as the distance metric. Cluster density was computed using convex hull area. Cluster density was computed using convex hull area.

##### *Atypical Localization of Immature Precursors (ALIP) Analysis*

Atypical localization of immature precursors (ALIP) clusters were defined as ≥2 myeloblasts and/or promyelocytes within 12 µm of one another and located >150 µm from trabeculae and >40 µm from vasculature.

#### MDS-MAPS Detailed Implementation

##### *Patch Definition*

WSIs were subdivided into non-overlapping 256 × 256 µm patches. Exclusion criteria included patches with <100 cells or >95% non-hematopoietic elements.

##### *Feature Scaling*

Features were scaled using z-score normalization derived exclusively from diagnostic samples. Scaling parameters were locked prior to longitudinal analysis.

##### *Weight Derivation*

Weights were derived using diagnostic samples only and locked prior to remission modeling. For each 256×256 μm patch, 82 features spanning cell-type proportions, cell morphology, spatial proximity metrics, and microenvironmental localization were computed and z-score normalized across the diagnostic cohort. Feature weights were derived in two steps. First, features were ranked by their correlation with disease severity through unsupervised sample stratification, identifying disease-associated perturbations receiving positive weights (e.g., myeloblast proportion, ALIP clusters) and normal bone marrow features receiving negative weights (e.g., erythroid spatial density, fat-to-cell ratio). Second, weights were adjusted through expert curation by a board-certified hematopathologist (SSP) according to established MDS pathological criteria, organized into four biological domains: blast burden, erythroid dysplasia, megakaryocyte dysplasia, and microenvironmental features. Biologically-motivated interaction terms were additionally incorporated as products of z-scored feature pairs reflecting known pathological co-occurrences, including myeloblast–megakaryocyte proportion (multi-lineage dysplasia), myeloblast proportion–trabecular distance (ALIP pattern), and MMC proportion–fat elements (preserved normal hematopoiesis). This amounted to 32 total weighted features and 4 interaction terms (**Supplementary Table 6**). The patch-level MAPS was then computed as:

$$MAP = \Sigma\, (Z(i) \times W(i)) + \Sigma\, (Z(j) \times Z(k) \times W(jk))$$

where Z(i) is the z-score of feature i, W(i) is its assigned weight, and Z(j) × Z(k) × W(jk) represents weighted interaction terms between feature pairs j and k.

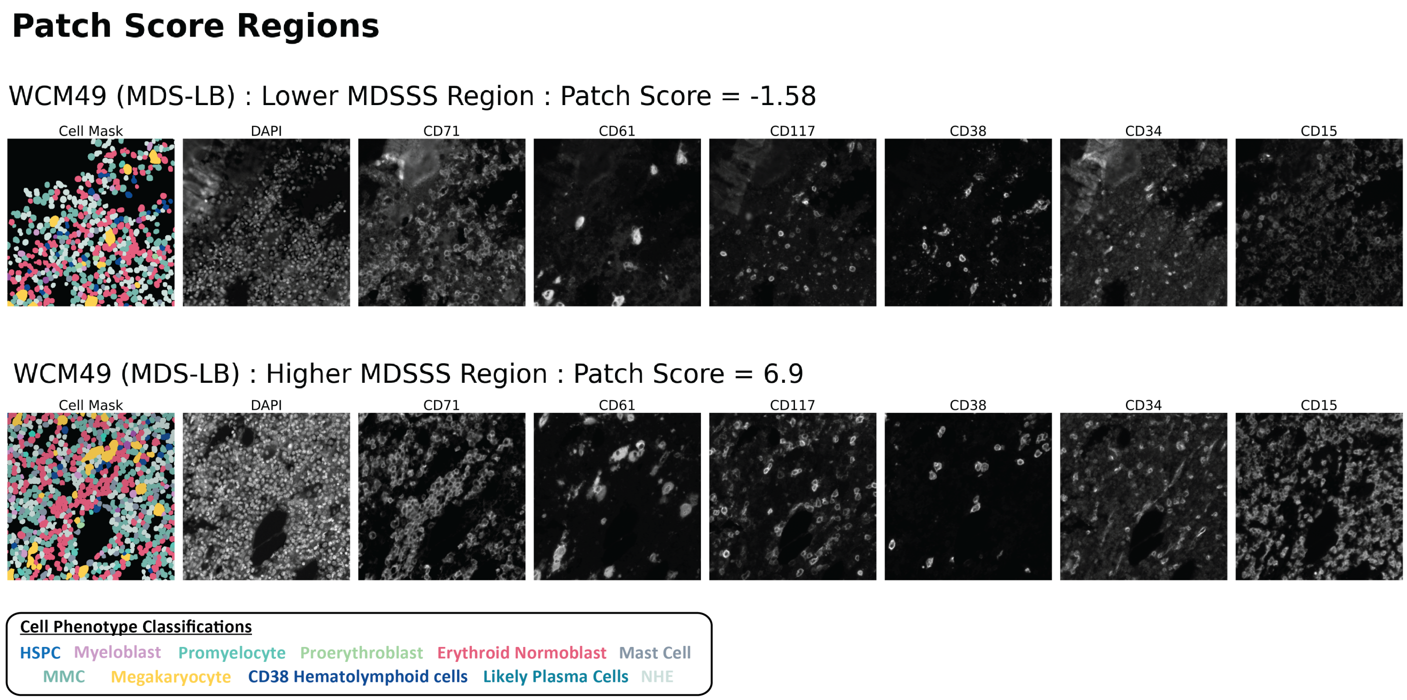

**CCUS/MDS-LB Discrimination and Remission Classification Modeling**

To evaluate the ability of MAPS to discriminate CCUS from MDS-LB and separately, remission from active disease, logistic regression models were implemented using leave-one-out cross-validation (LOOCV) for CCUS/MDS-LB discrimination or a leave-one-patient-out cross-validation (LOPO-CV) framework for remission classification. L2-regularized logistic regression was applied with a regularization strength of C = 1.0, using the lbfgs solver. To prevent data leakage, feature standardization was performed independently within each training fold, and the derived scaling parameters were applied to the corresponding held-out patient samples (or individual samples in LOOCV analysis).

Statistical significance of incremental discrimination (ΔAUC) was assessed using permutation testing with 1,000 iterations, in which MAPS values were randomly permuted and the LOOCV or LOPO-CV procedures repeated to generate a null distribution. Bootstrap confidence intervals were not calculated. Specifically for remission classification, sensitivity analyses were performed through stratified evaluation separating stem cell transplantation (SCT)–associated remission samples from non-SCT remission samples, as presented in the corresponding figure.

**Blast Progression Modeling**

To evaluate the relative contributions of clinical and spatial features to blast progression, analyses were performed in a subset of 16 patients for whom follow-up data were available, including 8 patients who experienced an increase in marrow blast percentage and 8 who remained stable.

Two L2-regularized logistic regression models were constructed with a regularization strength of C = 0.5 to mitigate overfitting in this limited cohort. The clinical model incorporated established clinical and laboratory variables, including IPSS-M score, mean corpuscular volume (MCV), age at diagnosis, hemoglobin level, absolute neutrophil count (ANC), and platelet count. The spatial model included mean MAPS along with the top five multiplex immunofluorescence (MxIF)–derived features ranked by univariate non-parametric AUC.

Feature importance was evaluated using SHAP (SHapley Additive exPlanations) values, calculated as the product of each model coefficient and the corresponding standardized feature value. SHAP values were sign-corrected based on coefficient directionality to facilitate interpretation, such that positive values indicated association with blast increase and negative values indicated association with stability.

Bootstrap confidence intervals were generated using 1,000 resamples with replacement, refitting the model in each iteration. Model discrimination was quantified using a non-parametric AUC computed as the Mann–Whitney U statistic normalized by group sizes, representing the fraction of correctly ranked stable versus blast-increase patient pairs.

**Mixed-Effects Modeling**

To assess whether changes in MAPS were associated with remission status independent of blast burden while accounting for repeated measurements within patients, a linear mixed-effects model was fitted. The model specified MAPS as the continuous outcome and included fixed effects for remission status and blast percentage, with a random intercept for each patient to account for within-patient correlation across serial biopsies.

An identity link function was used, as the outcome variable (MAPS) was continuous. Only random intercepts were included; random slopes were not modeled due to the limited number of serial samples per patient. Statistical significance of fixed effects was assessed using Wald tests. Formal residual diagnostics were not performed.

Linear mixed-effects model:

$$MAPS \sim Remission + Blast\% + (1|Patient)$$

**Public Dataset Re-Analysis**

To evaluate transcriptional correlates of spatial niche remodeling observed in our imaging cohort, we re-analyzed publicly available bulk and single-cell transcriptomic datasets of $CD34^+$ hematopoietic progenitor populations from patients with MDS and healthy controls. Datasets included GSE136816, GSE111085, and EGAS00001007568. In addition, protein-level expression data were examined from the mass cytometry dataset reported by Behbehani *et al.*

For bulk RNA sequencing datasets (GSE136816 and GSE111085), raw gene-level count matrices were obtained when available. Expression values were analyzed in count space and normalized using the trimmed mean of M-values (TMM) method. Differential expression testing was performed using generalized linear modeling with likelihood ratio tests. Multiple hypothesis testing correction was applied using the Benjamini–Hochberg procedure, and genes were considered significantly differentially expressed at a false discovery rate (FDR) < 0.05 (adjusted $P < 0.05$).

For the single-cell RNA sequencing dataset (EGAS00001007568), normalized expression matrices were used as provided by the original study authors. Cell-type annotations were retained from the published metadata. Gene expression comparisons between MDS and healthy $CD34^+$ progenitor populations were performed using rank-based or model-based testing as appropriate to the dataset structure, with multiple testing correction applied using the Benjamini–Hochberg method (FDR < 0.05).

Mass cytometry data from Behbehani *et al.* were analyzed at the sample level to quantify relative CXCR4 protein expression across hematopoietic cell populations. Protein abundance values were used as reported in the original publication, and comparisons between diagnostic categories were performed using nonparametric statistical testing.

Across all datasets, statistical significance was defined as an adjusted *P* value < 0.05.

**Code and Data Availability**

Custom scripts were written in Python (v3.11) using Scikit-learn, HDBSCAN, NetworkX, Scipy, Statsmodels, Scimap, and SHAP. Derived feature matrices are available upon request. Whole-slide images are available under institutional data use agreement due to protected health information.

***Abbreviations Used in Manuscript***

| **Term** | **Abbreviation** |
|---|---|
| Artifical Intelligence | AI |
| Atypical Localization of Immature Precursors | ALIP |
| Clonal Cytopenia of Undetermined Significance | CCUS |
| Clonal Hematopoiesis of Indeterminate Potential | CHIP |
| C-X-C motif chemokine ligand 12 | CXCL12 |
| C-X-C motif chemokine receptor 4 | CXCR4 |
| Disease-Modifying Therapy | DMT |
| Hematopoietic Stem Cell Transplantation | HSCT |
| Hematopoietic Stem and Progenitor Cells | HSPCs |
| International Consensus Classification | ICC |
| International Prognostic Scoring System-Molecular | IPSS-M |
| Myelodysplastic Neoplasms | MDS |
| MDS with Increased Blasts | MDS-IB |
| MDS with Low Blasts | MDS-LB |
| MDS Severity Score | MDSSS |
| Multiparameter Flow Cytometry | MFC |
| Maturing Myeloid Cells | MMCs |
| Multiplex Immunofluorescence | MxIF |
| Non-Hematopoietic Elements | NHEs |
| Normal Bone Marrow | NBM |
| t-distributed Stochastic Neighbor Embedding | t-SNE |
| Weill Cornell Medicine/NewYork-Presbyterian Hospital | WCM/NYP |
| World Health Organization | WHO |

| | |
|---|---|
| Whole Slide Images | WSIs |

**Supplemental Table 2. Cell type quantifications by flow cytometric immunophenotyping (FCM).**

| Case ID | %lymphocytes | %granulocytes | %monocytes | %monos+grans | %CD34+/CD117+ myeloblasts | %CD117+/CD15+ promyelocytes | %CD15+ | %CD71+ | %CD117+/CD71+ proerytherythroblasts | %B-cell precursors | %plasma cells | %mast cells |
|---|---|---|---|---|---|---|---|---|---|---|---|---|
| WCM2 | 9.400% | 84.600% | 2.300% | 86.900% | 2.200% | N/A | N/A | 2.100% | 1.200% | 0.400% | N/A | 0.007% |
| WCM2_2 | 20.100% | 69.500% | 5.300% | 74.800% | 0.200% | N/A | N/A | 0.800% | 0.200% | 1.070% | N/A | 0.007% |
| WCM3 | 16.000% | 59.100% | 6.400% | 65.500% | 5.000% | 4.800% | 65.800% | 9.700% | 3.100% | 1.200% | 0.600% | 0.050% |
| WCM4 | 13.200% | 81.100% | 3.400% | 84.500% | 0.200% | N/A | N/A | N/A | N/A | 0.042% | N/A | 0.017% |
| WCM5 | 15.200% | 75.000% | 4.500% | 79.500% | 0.800% | N/A | N/A | 4.500% | 1.800% | 0.210% | 0.200% | 0.120% |
| WCM6 | 15.000% | 77.300% | 6.300% | 83.600% | 0.100% | N/A | N/A | 0.200% | 0.037% | 0.017% | 0.000% | 0.002% |
| WCM7 | 14.700% | 66.100% | 3.900% | 70.000% | 0.600% | N/A | N/A | 6.700% | 1.400% | 0.180% | N/A | 0.016% |
| WCM8 | 23.700% | 66.200% | 6.900% | 73.100% | N/A | N/A | N/A | N/A | N/A | 0.130% | N/A | N/A |
| WCM9 | 10.400% | 79.000% | 2.800% | 81.800% | 0.700% | N/A | N/A | N/A | N/A | 1.400% | 0.058% | 0.000% |
| WCM10 | 8.500% | 86.100% | 2.000% | 88.100% | N/A | N/A | N/A | N/A | N/A | 0.049% | N/A | N/A |
| WCM11 | 16.000% | 75.400% | 5.700% | 81.100% | N/A | N/A | N/A | N/A | N/A | 0.200% | N/A | N/A |
| WCM12 | 3.100% | 69.200% | 2.600% | 71.800% | 5.400% | N/A | N/A | 15.400% | 2.800% | 0.100% | 0.200% | 0.100% |
| WCM13 | 20.800% | 52.200% | 6.600% | 58.800% | 7.000% | N/A | N/A | N/A | N/A | 0.100% | 0.130% | 0.000% |
| WCM13_2 | 3.000% | 89.500% | 2.600% | 92.100% | 0.800% | N/A | N/A | 1.600% | 0.500% | 0.004% | N/A | 0.016% |
| WCM13_3 | 33.000% | 27.900% | 5.900% | 33.800% | 18.000% | N/A | N/A | N/A | N/A | 0.300% | N/A | 0.037% |
| WCM14 | 24.800% | 61.900% | 2.500% | 64.400% | 4.500% | N/A | N/A | 5.600% | 2.300% | 0.876% | N/A | 0.010% |
| WCM14_2 | 38.700% | 48.300% | 0.900% | 49.200% | 4.000% | N/A | N/A | 7.900% | 4.100% | 0.300% | 0.052% | 0.020% |
| WCM14_4 | 90.600% | 1.100% | 0.800% | 1.900% | 4.000% | N/A | N/A | N/A | N/A | 0.012% | N/A | 0.000% |
| WCM15 | 12.000% | 71.300% | 7.100% | 78.400% | 1.700% | N/A | N/A | 3.900% | 2.400% | 0.240% | 0.550% | 0.012% |
| WCM15_2 | 13.000% | 58.000% | 18.200% | 76.200% | 1.000% | N/A | N/A | 0.680% | 0.278% | 0.029% | N/A | 0.004% |
| WCM15_3 | 19.000% | 64.600% | 7.800% | 72.400% | 0.600% | 0.040% | 67.800% | N/A | N/A | 0.019% | 0.008% | 0.010% |
| WCM16 | 9.200% | 50.200% | 1.900% | 52.100% | 16.000% | N/A | N/A | 10.800% | 5.000% | 0.210% | N/A | 0.080% |
| WCM16_2 | 2.500% | 86.800% | 2.500% | 89.300% | 1.000% | N/A | N/A | 5.200% | 1.100% | 0.030% | N/A | 0.006% |
| WCM17 | 10.100% | 74.400% | 3.300% | 77.700% | 7.200% | 1.300% | 79.000% | 1.400% | 0.600% | 0.069% | 0.074% | 0.000% |
| WCM17_2 | 38.900% | 1.400% | 39.100% | 40.500% | 12.000% | N/A | N/A | 4.300% | 1.500% | 0.022% | N/A | 0.023% |
| WCM17_3 | 0.400% | 93.600% | 0.700% | 94.300% | 0.119% | N/A | N/A | 0.400% | 0.100% | 2.150% | N/A | 0.000% |
| WCM18 | 10.200% | 75.200% | 2.200% | 77.400% | 0.400% | 0.900% | 76.300% | N/A | N/A | 0.100% | 0.200% | 0.030% |
| WCM19 | 22.900% | 60.500% | 3.700% | 64.200% | 5.000% | N/A | N/A | 1.900% | 0.320% | 0.002% | 0.100% | 0.070% |
| WCM20 | 46.300% | 41.300% | 4.500% | 45.800% | 6.000% | 1.100% | 45.500% | N/A | N/A | 0.175% | 0.100% | 0.400% |
| WCM21 | 21.600% | 62.400% | 8.400% | 70.800% | 2.000% | 0.300% | 71.700% | 0.300% | 0.074% | 0.000% | 0.014% | 0.005% |
| WCM21_2 | 44.200% | 37.200% | 3.300% | 40.500% | 9.700% | N/A | N/A | N/A | N/A | 0.004% | N/A | 0.040% |
| WCM21_3 | 32.700% | 41.900% | 2.900% | 44.800% | 6.000% | N/A | N/A | N/A | N/A | 0.000% | N/A | 0.300% |
| WCM21_4 | 46.500% | 51.300% | 0.000% | 51.300% | 0.040% | N/A | N/A | N/A | N/A | 0.000% | N/A | 0.000% |
| WCM21_5 | 0.200% | 96.100% | 2.500% | 98.600% | 0.032% | N/A | N/A | 0.400% | 0.044% | 0.004% | N/A | 0.000% |
| WCM22 | 15.000% | 79.400% | 2.500% | 81.900% | 0.500% | N/A | N/A | 1.600% | 0.300% | 0.100% | 0.100% | 0.032% |
| WCM22_2 | 11.300% | 73.900% | 4.900% | 78.800% | 0.900% | 0.500% | 74.200% | 12.000% | 5.100% | 0.010% | 0.100% | 0.000% |
| WCM23 | 27.400% | 56.100% | 7.100% | 63.200% | 4.800% | N/A | N/A | 0.880% | 0.560% | 0.015% | 0.095% | 0.018% |
| WCM23_2 | 19.600% | 60.400% | 11.300% | 71.700% | 4.000% | N/A | N/A | 1.500% | 1.300% | 0.014% | 0.010% | 0.028% |
| WCM24 | 29.200% | 65.000% | 2.800% | 67.800% | 1.700% | N/A | N/A | 0.500% | 0.100% | 0.000% | 0.029% | 0.000% |
| WCM25 | 6.500% | 86.200% | 3.100% | 89.300% | 2.300% | N/A | N/A | N/A | N/A | 0.195% | N/A | 0.005% |
| WCM25_2 | 6.900% | 69.200% | 5.900% | 75.100% | 12.000% | N/A | N/A | 1.900% | 0.700% | 1.220% | N/A | 0.017% |
| WCM26 | 10.700% | 84.500% | 1.500% | 86.000% | 1.200% | N/A | N/A | 1.900% | 0.300% | 0.500% | 0.026% | 0.010% |
| WCM27 | 9.400% | 72.000% | 2.800% | 74.800% | 0.900% | N/A | N/A | 9.700% | 1.800% | 0.240% | 0.300% | 0.700% |
| WCM28 | 27.000% | 22.000% | 13.400% | 35.400% | 21.300% | N/A | N/A | 5.800% | 1.300% | 0.000% | 1.200% | 0.400% |
| WCM29 | 14.100% | 70.400% | 5.400% | 75.800% | 1.400% | N/A | N/A | 0.400% | 0.100% | 0.300% | 0.025% | 0.023% |
| WCM30 | 10.900% | 80.800% | 3.400% | 84.200% | N/A | N/A | N/A | N/A | N/A | 0.616% | N/A | N/A |
| WCM31 | 3.500% | 88.200% | 2.600% | 90.800% | N/A | N/A | N/A | N/A | N/A | 0.910% | N/A | N/A |
| WCM32 | 29.400% | 58.300% | 8.100% | 66.400% | 0.200% | N/A | N/A | N/A | N/A | 0.440% | 0.600% | 0.100% |
| WCM33 | 8.000% | 84.700% | 3.400% | 88.100% | N/A | N/A | N/A | N/A | N/A | 0.198% | 0.500% | N/A |
| WCM34 | 12.900% | 76.900% | 7.200% | 84.100% | N/A | N/A | N/A | N/A | N/A | 1.000% | 0.005% | N/A |
| WCM35 | 14.100% | 78.000% | 2.900% | 80.900% | 0.300% | N/A | N/A | 2.800% | 0.300% | 0.100% | N/A | 0.000% |
| WCM36 | 9.900% | 84.200% | 3.800% | 88.000% | 0.300% | N/A | N/A | 0.600% | 0.200% | 0.030% | 0.100% | 0.008% |
| WCM37 | 2.100% | 82.900% | 2.900% | 85.800% | 0.400% | N/A | N/A | 0.700% | 0.100% | 0.000% | N/A | 0.012% |
| WCM38 | 5.500% | 86.600% | 1.300% | 87.900% | 0.200% | N/A | N/A | N/A | N/A | 0.646% | N/A | 0.022% |
| WCM39 | 21.600% | 70.500% | 4.500% | 75.000% | 0.070% | 0.200% | 72.900% | 1.600% | 0.200% | 0.796% | 0.012% | 0.007% |
| WCM40 | 1.600% | 94.600% | 1.600% | 96.200% | 0.100% | N/A | N/A | 0.400% | 0.100% | 0.420% | 0.019% | 0.100% |
| WCM41 | 4.000% | 84.100% | 5.300% | 89.400% | 0.400% | N/A | N/A | 2.100% | 0.700% | 0.058% | N/A | 0.009% |
| WCM42 | 13.900% | 75.900% | 3.500% | 79.400% | 0.600% | 1.900% | 78.300% | N/A | N/A | 0.631% | 0.260% | 0.048% |
| WCM44 | 38.900% | 58.400% | 0.200% | 58.600% | 0.700% | N/A | N/A | 0.200% | 0.022% | 0.000% | N/A | 0.000% |
| WCM44_2 | 29.100% | 55.300% | 3.800% | 59.100% | 9.700% | 2.300% | 56.500% | 0.900% | 0.039% | 0.165% | 0.027% | 0.002% |
| WCM44_3 | 40.300% | 48.700% | 1.700% | 50.400% | 1.800% | 4.500% | 52.200% | 1.600% | 0.100% | 0.000% | 0.047% | 0.007% |
| WCM44_4 | 73.800% | 17.800% | 3.000% | 20.800% | 0.017% | N/A | N/A | 0.300% | 0.024% | N/A | N/A | 0.000% |
| WCM44_5 | 61.000% | 29.500% | 3.800% | 33.300% | 0.100% | 0.100% | 29.600% | 0.800% | 0.100% | N/A | 0.039% | 0.005% |
| WCM44_6 | 25.400% | 65.800% | 5.100% | 70.900% | 0.100% | N/A | N/A | 0.500% | 0.100% | N/A | N/A | 0.009% |
| WCM44_7 | 63.900% | 14.400% | 1.000% | 15.400% | 2.000% | N/A | N/A | 5.100% | 0.700% | N/A | N/A | 0.009% |
| WCM45 | 21.000% | 58.700% | 6.800% | 65.500% | 0.600% | 1.200% | 66.800% | 4.100% | 0.500% | 0.482% | 0.400% | 0.044% |
| WCM45_2 | 11.000% | 72.100% | 8.600% | 80.700% | 0.560% | N/A | N/A | 2.500% | 0.400% | 0.200% | N/A | 0.000% |
| WCM46 | 14.800% | 76.800% | 3.400% | 80.200% | 0.265% | N/A | N/A | 1.700% | 0.300% | 0.145% | N/A | 0.025% |

| | | | | | | | | | | | | |
|---|---|---|---|---|---|---|---|---|---|---|---|---|
| WCM46_2 | 17.000% | 71.700% | 4.400% | 76.100% | 0.900% | N/A | N/A | 1.400% | 0.300% | 0.116% | N/A | 0.042% |
| WCM46_3 | 30.400% | 53.300% | 3.400% | 56.700% | 0.500% | N/A | N/A | 2.900% | 0.700% | 0.900% | N/A | 0.100% |
| WCM46_4 | 4.300% | 87.000% | 2.700% | 89.700% | 0.815% | N/A | N/A | 0.800% | 0.200% | 1.080% | N/A | 0.006% |
| WCM47 | 17.900% | 71.400% | 4.800% | 76.200% | 0.600% | 0.800% | 75.200% | 0.500% | 0.200% | 0.312% | 0.037% | 0.007% |
| WCM49 | 30.000% | 54.100% | 3.000% | 57.100% | 0.950% | N/A | N/A | 1.400% | 0.200% | 0.718% | N/A | 0.008% |
| WCM49_2 | 22.600% | 60.500% | 6.300% | 66.800% | 9.000% | N/A | N/A | 2.800% | 0.600% | 0.023% | N/A | 0.030% |
| WCM50 | 13.000% | 74.000% | 3.800% | 77.800% | 0.209% | 0.400% | 74.600% | 5.300% | 0.300% | 0.008% | 0.100% | 0.100% |
| WCM51 | 49.700% | 36.200% | 8.500% | 44.700% | 0.200% | N/A | N/A | 0.100% | 0.009% | 0.200% | 0.017% | 0.012% |
| WCM52 | 3.700% | 89.900% | 2.900% | 92.800% | 0.137% | | | 1.600% | 0.400% | 0.024% | N/A | 0.018% |
| WCM53 | 6.300% | 79.600% | 4.400% | 84.000% | 0.575% | N/A | N/A | N/A | N/A | 0.910% | N/A | 0.000% |
| WCM54 | 8.600% | 81.100% | 4.900% | 86.000% | N/A | N/A | N/A | N/A | N/A | 0.092% | 0.200% | N/A |
| WCM55 | 12.100% | 80.800% | 4.100% | 84.900% | 0.400% | N/A | N/A | N/A | N/A | 0.009% | N/A | 0.009% |
| WCM56 | 16.200% | 77.300% | 3.100% | 80.400% | N/A | N/A | N/A | N/A | N/A | 0.380% | 0.020% | N/A |
| WCM57 | 6.700% | 90.000% | 2.400% | 92.400% | N/A | N/A | N/A | N/A | N/A | 0.032% | 0.030% | N/A |
| WCM60 | 12.500% | 77.200% | 4.300% | 81.500% | N/A | N/A | N/A | N/A | N/A | N/A | N/A | N/A |
| WCM62 | 7.300% | 73.200% | 2.900% | 76.100% | N/A | N/A | N/A | N/A | N/A | 1.040% | N/A | N/A |
| WCM64 | 3.800% | 89.800% | 3.500% | N/A | N/A | N/A | N/A | N/A | N/A | 0.238% | N/A | N/A |
| WCM65 | 7.200% | 87.000% | 3.500% | 90.500% | 0.600% | N/A | N/A | N/A | N/A | 0.825% | 0.100% | 0.000% |
| WCM66 | 10.800% | 83.200% | 3.300% | 86.500% | 0.200% | N/A | N/A | 1.100% | 0.100% | 0.336% | 0.169% | 0.001% |
| WCM67 | 5.100% | 87.700% | 5.000% | 92.700% | N/A | N/A | N/A | N/A | N/A | 0.415% | 0.046% | N/A |
| WCM70 | 2.900% | 90.900% | 1.900% | 92.800% | 0.414% | N/A | N/A | N/A | N/A | 0.589% | N/A | 0.000% |
| WCM71 | 4.600% | 10.000% | 8.400% | 18.400% | 4.900% | 3.100% | 21.700% | 57.400% | 17.300% | N/A | 0.200% | 0.300% |
| WCM72 | 34.600% | 51.000% | 8.800% | 59.800% | 2.000% | N/A | N/A | N/A | N/A | 0.200% | 0.300% | 0.100% |
| WCM77 | 23.700% | 63.000% | 4.900% | 67.900% | 1.000% | N/A | N/A | 0.400% | 0.100% | 0.368% | N/A | 0.021% |

**Supplemental Table 2. Cell type quantifications by flow cytometric immunophenotyping (FCM).**

| Case ID | %lymphocytes | %granulocytes | %monocytes | %monos+grans | %CD34+/CD117+ myeloblasts | %CD117+/CD15+ promyelocytes | %CD15+ | %CD71+ | %CD117+/CD71+ proerytherythroblasts | %B-cell precursors | %plasma cells | %mast cells |
|---|---|---|---|---|---|---|---|---|---|---|---|---|
| WCM1 | N/A | N/A | N/A | N/A | N/A | N/A | N/A | N/A | N/A | N/A | N/A | N/A |
| WCM2 | 9.400% | 84.600% | 2.300% | 86.900% | 2.200% | N/A | N/A | 2.100% | 1.200% | 0.400% | N/A | 0.007% |
| WCM2_2 | 20.100% | 69.500% | 5.300% | 74.800% | 0.200% | N/A | N/A | 0.800% | 0.200% | 1.070% | N/A | 0.007% |
| WCM3 | 16.000% | 59.100% | 6.400% | 65.500% | 5.000% | 4.800% | 65.800% | 9.700% | 3.100% | 1.200% | 0.600% | 0.050% |
| WCM4 | 13.200% | 81.100% | 3.400% | 84.500% | 0.200% | N/A | N/A | N/A | N/A | 0.042% | N/A | 0.017% |
| WCM5 | 15.200% | 75.000% | 4.500% | 79.500% | 0.800% | N/A | N/A | 4.500% | 1.800% | 0.210% | 0.200% | 0.120% |
| WCM6 | 15.000% | 77.300% | 6.300% | 83.600% | 0.100% | N/A | N/A | 0.200% | 0.037% | 0.017% | 0.000% | 0.002% |
| WCM7 | 14.700% | 66.100% | 3.900% | 70.000% | 0.600% | N/A | N/A | 6.700% | 1.400% | 0.180% | N/A | 0.016% |
| WCM8 | 23.700% | 66.200% | 6.900% | 73.100% | N/A | N/A | N/A | N/A | N/A | 0.130% | N/A | N/A |
| WCM9 | 10.400% | 79.000% | 2.800% | 81.800% | 0.700% | N/A | N/A | N/A | N/A | 1.400% | 0.058% | 0.000% |
| WCM10 | 8.500% | 86.100% | 2.000% | 88.100% | N/A | N/A | N/A | N/A | N/A | 0.049% | N/A | N/A |
| WCM11 | 16.000% | 75.400% | 5.700% | 81.100% | N/A | N/A | N/A | N/A | N/A | 0.200% | N/A | N/A |
| WCM12 | 3.100% | 69.200% | 2.600% | 71.800% | 5.400% | N/A | N/A | 15.400% | 2.800% | 0.100% | 0.200% | 0.100% |
| WCM13 | 20.800% | 52.200% | 6.600% | 58.800% | 7.000% | N/A | N/A | N/A | N/A | 0.100% | 0.130% | 0.000% |
| WCM13_2 | 3.000% | 89.500% | 2.600% | 92.100% | 0.800% | N/A | N/A | 1.600% | 0.500% | 0.004% | N/A | 0.016% |
| WCM13_3 | 33.000% | 27.900% | 5.900% | 33.800% | 18.000% | N/A | N/A | N/A | N/A | 0.300% | N/A | 0.037% |
| WCM14 | 24.800% | 61.900% | 2.500% | 64.400% | 4.500% | N/A | N/A | 5.600% | 2.300% | 0.876% | N/A | 0.010% |
| WCM14_2 | 38.700% | 48.300% | 0.900% | 49.200% | 4.000% | N/A | N/A | 7.900% | 4.100% | 0.300% | 0.052% | 0.020% |
| WCM14_3 | N/A | N/A | N/A | N/A | N/A | N/A | N/A | N/A | N/A | N/A | N/A | N/A |
| WCM14_4 | 90.600% | 1.100% | 0.800% | 1.900% | 4.000% | N/A | N/A | N/A | N/A | 0.012% | N/A | 0.000% |
| WCM15 | 12.000% | 71.300% | 7.100% | 78.400% | 1.700% | N/A | N/A | 3.900% | 2.400% | 0.240% | 0.550% | 0.012% |
| WCM15_2 | 13.000% | 58.000% | 18.200% | 76.200% | 1.000% | N/A | N/A | 0.680% | 0.278% | 0.029% | N/A | 0.004% |
| WCM15_3 | 19.000% | 64.600% | 7.800% | 72.400% | 0.600% | 0.040% | 67.800% | N/A | N/A | 0.019% | 0.008% | 0.010% |
| WCM16 | 9.200% | 50.200% | 1.900% | 52.100% | 16.000% | N/A | N/A | 10.800% | 5.000% | 0.210% | N/A | 0.080% |
| WCM16_2 | 2.500% | 86.800% | 2.500% | 89.300% | 1.000% | N/A | N/A | 5.200% | 1.100% | 0.030% | N/A | 0.006% |
| WCM17 | 10.100% | 74.400% | 3.300% | 77.700% | 7.200% | 1.300% | 79.000% | 1.400% | 0.600% | 0.069% | 0.074% | 0.000% |
| WCM17_2 | 38.900% | 1.400% | 39.100% | 40.500% | 12.000% | N/A | N/A | 4.300% | 1.500% | 0.022% | N/A | 0.023% |
| WCM17_3 | 0.400% | 93.600% | 0.700% | 94.300% | 0.119% | N/A | N/A | 0.400% | 0.100% | 2.150% | N/A | 0.000% |
| WCM18 | 10.200% | 75.200% | 2.200% | 77.400% | 0.400% | 0.900% | 76.300% | N/A | N/A | 0.100% | 0.200% | 0.030% |
| WCM19 | 22.900% | 60.500% | 3.700% | 64.200% | 5.000% | N/A | N/A | 1.900% | 0.320% | 0.002% | 0.100% | 0.070% |
| WCM20 | 46.300% | 41.300% | 4.500% | 45.800% | 6.000% | 1.100% | 45.500% | N/A | N/A | 0.175% | 0.100% | 0.400% |
| WCM21 | 21.600% | 62.400% | 8.400% | 70.800% | 2.000% | 0.300% | 71.700% | 0.300% | 0.074% | 0.000% | 0.014% | 0.005% |
| WCM21_2 | 44.200% | 37.200% | 3.300% | 40.500% | 9.700% | N/A | N/A | N/A | N/A | 0.004% | N/A | 0.040% |
| WCM21_3 | 32.700% | 41.900% | 2.900% | 44.800% | 6.000% | N/A | N/A | N/A | N/A | 0.000% | N/A | 0.300% |
| WCM21_4 | 46.500% | 51.300% | 0.000% | 51.300% | 0.040% | N/A | N/A | N/A | N/A | 0.000% | N/A | 0.000% |
| WCM21_5 | 0.200% | 96.100% | 2.500% | 98.600% | 0.032% | N/A | N/A | 0.400% | 0.044% | 0.004% | N/A | 0.000% |
| WCM22 | 15.000% | 79.400% | 2.500% | 81.900% | 0.500% | N/A | N/A | 1.600% | 0.300% | 0.100% | 0.100% | 0.032% |
| WCM22_2 | 11.300% | 73.900% | 4.900% | 78.800% | 0.900% | 0.500% | 74.200% | 12.000% | 5.100% | 0.010% | 0.100% | 0.000% |
| WCM23 | 27.400% | 56.100% | 7.100% | 63.200% | 4.800% | N/A | N/A | 0.880% | 0.560% | 0.015% | 0.095% | 0.018% |
| WCM23_2 | 19.600% | 60.400% | 11.300% | 71.700% | 4.000% | N/A | N/A | 1.500% | 1.300% | 0.014% | 0.010% | 0.028% |
| WCM24 | 29.200% | 65.000% | 2.800% | 67.800% | 1.700% | N/A | N/A | 0.500% | 0.100% | 0.000% | 0.029% | 0.000% |
| WCM25 | 6.500% | 86.200% | 3.100% | 89.300% | 2.300% | N/A | N/A | N/A | N/A | 0.195% | N/A | 0.005% |
| WCM25_2 | 6.900% | 69.200% | 5.900% | 75.100% | 12.000% | N/A | N/A | 1.900% | 0.700% | 1.220% | N/A | 0.017% |
| WCM26 | 10.700% | 84.500% | 1.500% | 86.000% | 1.200% | N/A | N/A | 1.900% | 0.300% | 0.500% | 0.026% | 0.010% |
| WCM27 | 9.400% | 72.000% | 2.800% | 74.800% | 0.900% | N/A | N/A | 9.700% | 1.800% | 0.240% | 0.300% | 0.700% |
| WCM28 | 27.000% | 22.000% | 13.400% | 35.400% | 21.300% | N/A | N/A | 5.800% | 1.300% | 0.000% | 1.200% | 0.400% |
| WCM29 | 14.100% | 70.400% | 5.400% | 75.800% | 1.400% | N/A | N/A | 0.400% | 0.100% | 0.300% | 0.025% | 0.023% |
| WCM30 | 10.900% | 80.800% | 3.400% | 84.200% | N/A | N/A | N/A | N/A | N/A | 0.616% | N/A | N/A |
| WCM31 | 3.500% | 88.200% | 2.600% | 90.800% | N/A | N/A | N/A | N/A | N/A | 0.910% | N/A | N/A |
| WCM32 | 29.400% | 58.300% | 8.100% | 66.400% | 0.200% | N/A | N/A | N/A | N/A | 0.440% | 0.600% | 0.100% |
| WCM33 | 8.000% | 84.700% | 3.400% | 88.100% | N/A | N/A | N/A | N/A | N/A | 0.198% | 0.500% | N/A |
| WCM34 | 12.900% | 76.900% | 7.200% | 84.100% | N/A | N/A | N/A | N/A | N/A | 1.000% | 0.005% | N/A |
| WCM35 | 14.100% | 78.000% | 2.900% | 80.900% | 0.300% | N/A | N/A | 2.800% | 0.300% | 0.100% | N/A | 0.000% |
| WCM36 | 9.900% | 84.200% | 3.800% | 88.000% | 0.300% | N/A | N/A | 0.600% | 0.200% | 0.030% | 0.100% | 0.008% |
| WCM37 | 2.100% | 82.900% | 2.900% | 85.800% | 0.400% | N/A | N/A | 0.700% | 0.100% | 0.000% | N/A | 0.012% |
| WCM38 | 5.500% | 86.600% | 1.300% | 87.900% | 0.200% | N/A | N/A | N/A | N/A | 0.646% | N/A | 0.022% |
| WCM39 | 21.600% | 70.500% | 4.500% | 75.000% | 0.070% | 0.200% | 72.900% | 1.600% | 0.200% | 0.796% | 0.012% | 0.007% |
| WCM40 | 1.600% | 94.600% | 1.600% | 96.200% | 0.100% | N/A | N/A | 0.400% | 0.100% | 0.420% | 0.019% | 0.100% |
| WCM41 | 4.000% | 84.100% | 5.300% | 89.400% | 0.400% | N/A | N/A | 2.100% | 0.700% | 0.058% | N/A | 0.009% |
| WCM42 | 13.900% | 75.900% | 3.500% | 79.400% | 0.600% | 1.900% | 78.300% | N/A | N/A | 0.631% | 0.260% | 0.048% |
| WCM44 | 38.900% | 58.400% | 0.200% | 58.600% | 0.700% | N/A | N/A | 0.200% | 0.022% | 0.000% | N/A | 0.000% |
| WCM44_2 | 29.100% | 55.300% | 3.800% | 59.100% | 9.700% | 2.300% | 56.500% | 0.900% | 0.039% | 0.165% | 0.027% | 0.002% |
| WCM44_3 | 40.300% | 48.700% | 1.700% | 50.400% | 1.800% | 4.500% | 52.200% | 1.600% | 0.100% | 0.000% | 0.047% | 0.007% |
| WCM44_4 | 73.800% | 17.800% | 3.000% | 20.800% | 0.017% | N/A | N/A | 0.300% | 0.024% | N/A | N/A | 0.000% |
| WCM44_5 | 61.000% | 29.500% | 3.800% | 33.300% | 0.100% | 0.100% | 29.600% | 0.800% | 0.100% | N/A | 0.039% | 0.005% |
| WCM44_6 | 25.400% | 65.800% | 5.100% | 70.900% | 0.100% | N/A | N/A | 0.500% | 0.100% | N/A | N/A | 0.009% |

| | | | | | | | | | | | | |
|---|---|---|---|---|---|---|---|---|---|---|---|---|
| WCM44_7 | 63.900% | 14.400% | 1.000% | 15.400% | 2.000% | N/A | N/A | 5.100% | 0.700% | N/A | N/A | 0.009% |
| WCM45 | 21.000% | 58.700% | 6.800% | 65.500% | 0.600% | 1.200% | 66.800% | 4.100% | 0.500% | 0.482% | 0.400% | 0.044% |
| WCM45_2 | 11.000% | 72.100% | 8.600% | 80.700% | 0.560% | N/A | N/A | 2.500% | 0.400% | 0.200% | N/A | 0.000% |
| WCM46 | 14.800% | 76.800% | 3.400% | 80.200% | 0.265% | N/A | N/A | 1.700% | 0.300% | 0.145% | N/A | 0.025% |
| WCM46_2 | 17.000% | 71.700% | 4.400% | 76.100% | 0.900% | N/A | N/A | 1.400% | 0.300% | 0.116% | N/A | 0.042% |
| WCM46_3 | 30.400% | 53.300% | 3.400% | 56.700% | 0.500% | N/A | N/A | 2.900% | 0.700% | 0.900% | N/A | 0.100% |
| WCM46_4 | 4.300% | 87.000% | 2.700% | 89.700% | 0.815% | N/A | N/A | 0.800% | 0.200% | 1.080% | N/A | 0.006% |
| WCM47 | 17.900% | 71.400% | 4.800% | 76.200% | 0.600% | 0.800% | 75.200% | 0.500% | 0.200% | 0.312% | 0.037% | 0.007% |
| WCM49 | 30.000% | 54.100% | 3.000% | 57.100% | 0.950% | N/A | N/A | 1.400% | 0.200% | 0.718% | N/A | 0.008% |
| WCM49_2 | 22.600% | 60.500% | 6.300% | 66.800% | 9.000% | N/A | N/A | 2.800% | 0.600% | 0.023% | N/A | 0.030% |
| WCM50 | 13.000% | 74.000% | 3.800% | 77.800% | 0.209% | 0.400% | 74.600% | 5.300% | 0.300% | 0.008% | 0.100% | 0.100% |
| WCM51 | 49.700% | 36.200% | 8.500% | 44.700% | 0.200% | N/A | N/A | 0.100% | 0.009% | 0.200% | 0.017% | 0.012% |
| WCM52 | 3.700% | 89.900% | 2.900% | 92.800% | 0.137% | | | 1.600% | 0.400% | 0.024% | N/A | 0.018% |
| WCM53 | 6.300% | 79.600% | 4.400% | 84.000% | 0.575% | N/A | N/A | N/A | N/A | 0.910% | N/A | 0.000% |
| WCM54 | 8.600% | 81.100% | 4.900% | 86.000% | N/A | N/A | N/A | N/A | N/A | 0.092% | 0.200% | N/A |
| WCM55 | 12.100% | 80.800% | 4.100% | 84.900% | 0.400% | N/A | N/A | N/A | N/A | 0.009% | N/A | 0.009% |
| WCM56 | 16.200% | 77.300% | 3.100% | 80.400% | N/A | N/A | N/A | N/A | N/A | 0.380% | 0.020% | N/A |
| WCM57 | 6.700% | 90.000% | 2.400% | 92.400% | N/A | N/A | N/A | N/A | N/A | 0.032% | 0.030% | N/A |
| WCM59 | N/A | N/A | N/A | N/A | N/A | N/A | N/A | N/A | N/A | N/A | N/A | N/A |
| WCM60 | 12.500% | 77.200% | 4.300% | 81.500% | N/A | N/A | N/A | N/A | N/A | N/A | N/A | N/A |
| WCM61 | N/A | N/A | N/A | N/A | N/A | N/A | N/A | N/A | N/A | N/A | N/A | N/A |
| WCM62 | 7.300% | 73.200% | 2.900% | 76.100% | N/A | N/A | N/A | N/A | N/A | 1.040% | N/A | N/A |
| WCM64 | 3.800% | 89.800% | 3.500% | N/A | N/A | N/A | N/A | N/A | N/A | 0.238% | N/A | N/A |
| WCM65 | 7.200% | 87.000% | 3.500% | 90.500% | 0.600% | N/A | N/A | N/A | N/A | 0.825% | 0.100% | 0.000% |
| WCM66 | 10.800% | 83.200% | 3.300% | 86.500% | 0.200% | N/A | N/A | 1.100% | 0.100% | 0.336% | 0.169% | 0.001% |
| WCM67 | 5.100% | 87.700% | 5.000% | 92.700% | N/A | N/A | N/A | N/A | N/A | 0.415% | 0.046% | N/A |
| WCM68 | N/A | N/A | N/A | N/A | N/A | N/A | N/A | N/A | N/A | N/A | N/A | N/A |
| WCM69 | N/A | N/A | N/A | N/A | N/A | N/A | N/A | N/A | N/A | N/A | N/A | N/A |
| WCM70 | 2.900% | 90.900% | 1.900% | 92.800% | 0.414% | N/A | N/A | N/A | N/A | 0.589% | N/A | 0.000% |
| WCM71 | 4.600% | 10.000% | 8.400% | 18.400% | 4.900% | 3.100% | 21.700% | 57.400% | 17.300% | | 0.200% | 0.300% |
| WCM72 | 34.600% | 51.000% | 8.800% | 59.800% | 2.000% | N/A | N/A | N/A | N/A | 0.200% | 0.300% | 0.100% |
| WCM73 | N/A | N/A | N/A | N/A | N/A | N/A | N/A | N/A | N/A | N/A | N/A | N/A |
| WCM77 | 23.700% | 63.000% | 4.900% | 67.900% | 1.000% | N/A | N/A | 0.400% | 0.100% | 0.368% | N/A | 0.021% |

**Supplemental Table 3. Initial and longitudinal therapy, clinical response data, and overall outcome for MDS patients.**

| Case ID | Initial Therapy | DMT | PB CR | Time to CR | BM CR (B<5%) | Time to BM CR | CG remission (CGR) | Time to CGR | Blast Count Progression? | Prog to AML | FU time | A/D |
|---|---|---|---|---|---|---|---|---|---|---|---|---|
| WCM2 | Revlimid (lenalidomide) | 1 | 1 | 109 | N/A | N/A | 1 | 581 | 0 | 0 | 986 | 0 |
| WCM3 | azacytidine | 1 | 1 | 103 | N/A | N/A | 0 | N/A | 0 | 0 | 1002 | 0 |
| WCM4 | transfusion | 0 | N/A | N/A | N/A | N/A | N/A | N/A | N/A | 0 | 1047 | 0 |
| WCM6 | EPO/transfusion | 0 | N/A | N/A | N/A | N/A | N/A | N/A | N/A | 0 | 281 | 0 |
| WCM7 | EPO >1yr then Luspatercept | 1 | 0 | N/A | N/A | N/A | N/A | | N/A | 0 | 1046 | 0 |
| WCM12 | Decitabine | 1 | 0 | N/A | N/A | N/A | N/A | N/A | N/A | 0 | 65 | 1 |
| WCM13 | Decitabine | 1 | 1 | 109 | 1 | 53 | 1 | 53 | 0 | 0 | 251 | 0 |
| WCM14 | HMA/ven --> CPX-351 --> SCT | 1 | 0 | N/A | 0 | N/A | 0 | N/A | 1 | 1 | 428 | 1 |
| WCM15 | EPO --> aza/magrolimab | 1 | 0 | N/A | N/A | N/A | 0 | N/A | 1 | 0 | 445 | 0 |
| WCM16 | Aza/Ven | 1 | 1 | 76 | 1 | 34 | 0 | N/A | 0 | 0 | 352 | 1 |
| WCM17 | azacytidine | 1 | 1 | 153 | N/A | N/A | N/A | N/A | N/A | 0 | 1161 | 0 |
| WCM19 | Aza/Magrolimab | 1 | 0 | N/A | 0 | N/A | 0 | N/A | 1 | 0 | 331 | 0 |
| WCM20 | EPO | 0 | N/A | N/A | N/A | N/A | N/A | N/A | N/A | 0 | 47 | 1 |
| WCM21 | Aza/Ven | 1 | 0 | N/A | 1 | 63 | N/A | N/A | N/A | 0 | 539 | 1 |
| WCM22 | Aza/Ven | 1 | 0 | N/A | 1 | 29 | 0 | N/A | 1 | 0 | 115 | 1 |
| WCM23 | azacytidine then Aza/Ven | 1 | 1 | 406 | 1 | 392 | 1 | 392 | 0 | 0 | 658 | 1 |
| WCM24 | No intervention to date | 0 | N/A | N/A | N/A | N/A | N/A | N/A | N/A | 0 | 293 | 0 |
| WCM25 | EPO --> azacytidine | 1 | 0 | N/A | N/A | N/A | 0 | N/A | 1 | 0 | 296 | 0 |
| WCM26 | transfusions/EPO --> Decitabine --> Luspatercept | 1 | 1 | 109 | 0 | N/A | 0 | N/A | 1 | 0 | 913 | 0 |
| WCM27 | EPO >6mo then luspatercept | 1 | 0 | N/A | N/A | N/A | N/A | N/A | N/A | 0 | 815 | 1 |
| WCM29 | eltrombopag then decitabine | 1 | 0 | N/A | N/A | N/A | N/A | N/A | N/A | 0 | 672 | 0 |
| WCM44 | azacytidine/decitabine --> eltanexor --> venetoclax, LDAC, GO | 1 | 1 | 539 | 1 | 431 | N/A | N/A | 1 | 1 | 1387 | 1 |
| WCM45 | Neupogen/Aranesp then Decitabine | 1 | 0 | N/A | N/A | N/A | 0 | N/A | 0 | 0 | 1104 | 1 |
| WCM46 | azacytidine | 1 | 0 | N/A | N/A | N/A | 1 | 245 | 0 | 0 | 1807 | 0 |
| WCM49 | EPO --> OPN-305 --> SCT | 1 | 0 | N/A | N/A | N/A | N/A | N/A | 1 | 0 | 1764 | 0 |
| WCM50 | EPO | 0 | N/A | N/A | N/A | N/A | N/A | N/A | N/A | 0 | 109 | 1 |
| WCM51 | OPN-305 --> Azacytidine | 1 | 0 | N/A | N/A | N/A | 0 | N/A | 0 | 0 | 1073 | 1 |
| WCM52 | EPO | 0 | N/A | N/A | N/A | N/A | N/A | | N/A | 0 | 232 | 0 |
| WCM72 | transfusion --> APR-246 --> SCT | 1 | 0 | N/A | N/A | N/A | 1 | 196 | 1 | 0 | 411 | 1 |
| WCM73 | Aza/Ven | 1 | 0 | N/A | 1 | 47 | 0 | N/A | 0 | 1 | 453 | 1 |
| WCM77 | EPO/Revlimid | 1 | 1 | 89 | N/A | N/A | N/A | N/A | N/A | 0 | 447 | 0 |

DMT, disease modifying therapy; PB, peripheral blood; CR, complete remission; BM, bone marrow; CG, cytogenetic; FU, follow up; A, alive; D, deceased
EPO, erythropoietin; HMA, hypomethylating agent; Aza/Ven, azacitidine/venetoclax; LDAC, low dose Ara-C; SCT, stem cell transplant
0, event did not occur; 1, event occurred
All time values listed in days.

**Supplemental Table 4. Next-generation sequencing data.**

| Case ID | Variant count | Gene | Variant | VAF |
|---|---|---|---|---|
| WCM1 | None | | | |
| WCM2 | None | | | |
| WCM2_2 | None | | | |
| WCM3 | 1 | ASXL1 | c.2321_2322delGA; p.R774Ifs*12 | 38.0% |
| WCM3 | 2 | U2AF1 | c.101C>T; p.S34F | 43.0% |
| WCM4 | 1 | DNMT3A | c.2567_2568delAG; p.E856Gfs*7 | 52.0% |
| WCM4 | 2 | SF3B1 | c.2098A>G; p.K700E | 37.0% |
| WCM5 | 1 | SF3B1 | c.1998G>T; p.K666N | 26.3% |
| WCM5 | 2 | DNMT3A | c.2207G>A; pArg736His | 26.3% |
| WCM6 | 1 | SF3B1 | c.2098A>G, p.K700E | 41.7% |
| WCM6 | 2 | TET2 | c.5360_5361insG, p.N1787Kfs*2 | 20.8% |
| WCM6 | 3 | ASXL1 | c.2113delG, p.E705Sfs*20 | 4.5% |
| WCM6 | 4 | DOT1L | c.367G>A, p.E123K | 16.1% |
| WCM6 | 5 | TET2 | c.3820C>G, p.Q1274E | 2.7% |
| WCM6 | 6 | DNMT3A | c.2321A>G, p.E774G | 41.8% |
| WCM7 | 1 | SF3B1 | c.2098A>G; p.K700E | 21.6% |
| WCM7 | 2 | DNMT3A | c.2339T>G; p.Ile780Ser | 21.8% |
| WCM8 | 1 | DNMT3A | c.2339T>C; p.I780T | 4.2% |
| WCM11 | None | | | |
| WCM12 | 1 | TP53 | c.716A>G; p.Asn239Ser | 10.7% |
| WCM12 | 2 | TP53 | c.493C>T; p.Gln165* | 3.8% |
| WCM12 | 3 | TP53 | c.319T>G; p.Tyr107Asp | 5.0% |
| WCM13 | 1 | SF3B1 | c.1998G>T; p.Lys666Asn | 38.1% |
| WCM13_2 | None | | | |
| WCM13_3 | 1 | SF3B1 | c.1998G>T; p.Lys666Asn | 31.1% |
| WCM14 | 1 | TP53 | c.722C>T; p.Ser241Phe | 37.4% |
| WCM14 | 2 | TP53 | c.592G>T; p.Glu198* | 31.9% |
| WCM14 | 3 | DNMT3A | c.2023G>A; p.Val675Met | 30.4% |
| WCM14_2 | 1 | TP53 | c.722C>T; p.Ser241Phe | 34.6% |
| WCM14_2 | 2 | TP53 | c.592G>T; p.Glu198* | 38.1% |
| WCM14_2 | 3 | DNMT3A | c.2023G>A; p.Val675Met | 35.9% |
| WCM14_3 | 1 | TP53 | c.722C>T; p.Ser241Phe | 15.5% |
| WCM14_3 | 2 | TP53 | c.592G>T; p.Glu198* | 15.9% |
| WCM14_3 | 3 | DNMT3A | c.2023G>A; p.Val675Met | 19.5% |
| WCM14_4 | 1 | TP53 | c.722C>T, p.S241F | 6.3% |
| WCM14_4 | 2 | TP53 | c.592G>T, p.E198* | 6.9% |
| WCM14_4 | 3 | DNMT3A | c.2023G>A, p.V675M | 6.8% |
| WCM15 | None | | | |
| WCM15_2 | 1 | TP53 | c.536A>T; p.His179Leu | 16.5% |
| WCM15_3 | 1 | TP53 | c.536A>T, p.H179L | 2.0% |
| WCM15_3 | 2 | ZRSR2 | c.1338_1343dup6, p.S447_R448dup | 23.7% |
| WCM16 | 1 | TP53 | c.711G>A; p.Met237Ile | 45.0% |
| WCM16 | 2 | TP53 | c.626_627delGA; p.Arg209Lysfs*6 | 45.3% |
| WCM16_2 | 1 | TP53 | c.711G>A, p.M237I | 14.1% |
| WCM16_2 | 2 | TP53 | c.626_627delGA, p.R209Kfs*6 | 14.2% |
| WCM17 | 1 | SF3B1 | c.1998G>T, p.K666N | 14.3% |
| WCM17 | 2 | SF3B1 | c.1873C>T, p.R625C | 17.7% |
| WCM17 | 3 | BCOR | c.2341delA, p.T781Pfs*5 | 8.0% |
| WCM17 | 4 | BCOR | c.599_603del5, p.T200Sfs*11 | 9.3% |

| | | | | |
|---|---|---|---|---|
| WCM17_2 | 1 | SF3B1 | c.1998G>T; p.K666N | 5.0% |
| WCM17_3 | None | | | |
| WCM18 | 1 | SF3B1 | c.2098A>G; p.Lys700Glu | 42.3% |
| WCM18 | 2 | TET2 | c.2429A>G; pGln810Arg | 50.2% |
| WCM19 | 1 | DNMT3A | c.2645G>A; p.Arg882His | 36.3% |
| WCM19 | 2 | SRSF2 | c.284C>G; p.Pro95Arg | 51.5% |
| WCM19 | 3 | SETBP1 | c.2608G>A; p.Gly870Ser | 35.6% |
| WCM19 | 4 | ASXL1 | c.1934_1935insG; p.Gly646Trpfs*12 | 25.2% |
| WCM20 | 1 | TP53 | c.527G>A; p.C176Y | 14.2% |
| WCM20 | 2 | TP53 | c.393del; p.N131Kfs*39 | 13.4% |
| WCM21 | 1 | ASXL1 | c.2555C>A; p.S852* | 13.0% |
| WCM21 | 2 | RUNX1 | c.779delA; p.N260tfs*51 | 15.0% |
| WCM21 | 3 | SH2B3 | c.622G>C; p.E208Q | 42.0% |
| WCM21_5 | None | | | |
| WCM22_2 | 1 | KRAS | c.35G>T, p.G12V | 2.7% |
| WCM22_2 | 2 | TP53 | c.817C>T, p.R273C | 9.1% |
| WCM23 | 1 | ETV6 | c.592C>T, p.Q198* | 36.7% |
| WCM23 | 2 | SRSF2 | c.284C>A, p.P95H | 52.1% |
| WCM23 | 3 | SETBP1 | c.2608G>A, p.G870S | 35.2% |
| WCM23 | 4 | RUNX1 | c.109delA, p.S37Afs*11 | 39.0% |
| WCM23 | 5 | ASXL1 | c.1900_1922del23, p.E635Rfs*15 | 57.1% |
| WCM23 | 6 | ASXL1 | c.1900_1927delinsCGGA, p.E635_G643delinsR | 3.5% |
| WCM23 | 7 | GATA2 | c.71C>G, p.S24* | 2.0% |
| WCM23_2 | 1 | ETV6 | c.592C>T, p.Q198* | 44.0% |
| WCM23_2 | 2 | SRSF2 | c.284C>A, p.P95H | 61.9% |
| WCM23_2 | 3 | SETBP1 | c.2608G>A, p.G870S | 44.4% |
| WCM23_2 | 4 | RUNX1 | c.109delA, p.S37Afs*11 | 45.5% |
| WCM23_2 | 5 | ASXL1 | c.1900_1922del23, p.E635Rfs*15 | 63.0% |
| WCM23_2 | 6 | NRAS | c.38G>T, p.G13V | 4.5% |
| WCM24 | 1 | IDH2 | c.419G>A; p.Arg140Gln | 38.0% |
| WCM24 | 2 | SRSF2 | c.284C>A; p.Pro95His | 38.5% |
| WCM24 | 3 | ASXL1 | c.1934_1935insG; p.Gly646Trpfs*12 | 33.2% |
| WCM24 | 4 | STAG2 | c.1057_1058insCAAG; p.Gly353Alafs*6 | 18.5% |
| WCM24 | 5 | STAG2 | c.1758_1759insT; p.Asn587* | 34.0% |
| WCM24 | 6 | STAG2 | c.2284A>T; p.Lys762* | 2.5% |
| WCM25 | 1 | ASXL1 | c.1934_1935insG; p.Gly646Trpfs*12 | 40.6% |
| WCM25 | 2 | U2AF1 | c.470A>C; p.Gln157Pro | 47.6% |
| WCM25 | 3 | STAG2 | c.1195C>T; p.Gln399* | 6.5% |
| WCM25 | 4 | TET2 | c.5618T>C; p.Ile1873Thr | 47.1% |
| WCM25 | 5 | SMC3 | c.2535+1G>A; p.? (splice donor variant) | 25.6% |
| WCM25 | 6 | CSF3R | c.1853C>T, p.T618I | 2.7% |
| WCM25 | 7 | NF1 | c.5376C>A, p.C1792* | 31.2% |
| WCM25 | 8 | CUX1 | c.1778_1779insAG, p.K594Afs*7 | 1.5% |
| WCM25_2 | 1 | ASXL1 | c.1934_1935insG; p.Gly646Trpfs*12 | 35.1% |
| WCM25_2 | 2 | U2AF1 | c.470A>C; p.Gln157Pro | 45.1% |
| WCM25_2 | 3 | STAG2 | c.1195C>T; p.Gln399* | 57.4% |
| WCM25_2 | 4 | TET2 | c.5618T>C; p.Ile1873Thr | 41.0% |
| WCM25_2 | 5 | SMC3 | c.2535+1G>A; p.? (splice donor variant) | 10.1% |
| WCM25_2 | 6 | RUNX1 | c.508+1_508+2insGTCGAAGTGGAAGAGG (splice donor variant | 4.5% |
| WCM26 | 1 | EZH2 | c.221delT; p.Val74Glyfs*12 | 38.5% |
| WCM27 | 1 | SF3B1 | c.2098A>G; p.Lys700Glu | 36.2% |

| | | | | |
|---|---|---|---|---|
| WCM27 | 2 | MPL | c.1544G>T; p.Trp515Leu | 1.4% |
| WCM27 | 3 | DNMT3A | c.2083-2A>G; p.? (splice donor variant) | 41.2% |
| WCM28 | 1 | TP53 | c.916C>T, p.R306* | 30.3% |
| WCM28 | 2 | TP53 | c.375+1G>A, p.? | 28.8% |
| WCM28 | 3 | PIK3R3 | c.8A>G, p.N3S | 30.7% |
| WCM28 | 4 | ZFHX3 | c.9524_9594delins123, p.S3175Cfs*6 | 8.9% |
| WCM29 | 1 | SF3B1 | c.1997A>C; p.Lys666Thr | 34.6% |
| WCM29 | 2 | ASXL1 | c.4402G>A; p.Ala1468Thr | 36.2% |
| WCM31 | None | | | |
| WCM32 | 1 | TET2 | c.3803+1G>C, p.? | 2.7% |
| WCM32 | 2 | ASXL1 | c.1900_1922del23, p.E635Rfs*15 | 1.0% |
| WCM32 | 3 | CREBBP | c.6609_6611delACA, p.Q2216del | 3.8% |
| WCM33 | 1 | DNMT3A | c.1105A>T; p.I369F | 10.7% |
| WCM33 | 2 | GATA1 | c.337C>T; p.R113C | 16.5% |
| WCM34 | None | | | |
| WCM35 | 1 | DNMT3A | p.799T | 0.8% |
| WCM36 | 1 | ALK | c.1288T>A, p.S430T | 2.0% |
| WCM36 | 2 | CREBBP | c.6743_6745delAGC, p.Q2248del | 1.6% |
| WCM36 | 3 | CTCF | c.1054A>C, p.K352Q | 2.2% |
| WCM36 | 4 | TSC1 | c.2626-2dupA, p.? | 2.1% |
| WCM36 | 5 | SDHA | c.1274T>G, p.V425G | 3.5% |
| WCM37 | 1 | SETBP1 | c.2602G>A; p.D868N | 23.5% |
| WCM37 | 2 | SRSF2 | c.284_307del24; p.P95_R102del | 69.6% |
| WCM37 | 3 | ASXL1 | c.2155G>T; p.E719* | 47.2% |
| WCM37 | 4 | TET2 | c.651delC; p.V218Wfs*32 | 47.5% |
| WCM37 | 5 | TET2 | c.1842dupG; p.L615Afs*23 | 45.2% |
| WCM37 | 6 | NPM1 | c. 847-5_*20delins108; p.? | *** |
| WCM39 | 1 | ABL1 | c.1786C>T; p.Arg596* | 3.2% |
| WCM40 | 1 | ASXL1 | c.1748G>A; p.Trp583* | 4.5% |
| WCM40 | 2 | U2AF1 | c.101C>T; p.Ser34Phe | 7.8% |
| WCM40 | 3 | DNMT3A | c.1238_1259delinsC; p.Gly413_Lys420delinsAla | 6.9% |
| WCM41 | 1 | IDH2 | c419G>A; p.Arg140Gln | 41.0% |
| WCM41 | 2 | SRSF2 | c.284C>G; p.Pro95Arg | 40.0% |
| WCM42 | 1 | ASXL1 | c.1934dup; p.Gly646Trpfs*12 | 27.0% |
| WCM42 | 2 | SRSF2 | c.284C>T | 36.0% |
| WCM44 | 1 | BCOR | c.2428C>T; p.R810* | 21.0% |
| WCM44 | 2 | NRAS | c.35G>T; p.G12V | 18.0% |
| WCM44 | 3 | RUNX1 | c.921_925delCAGCGinsT; p.S308Afs*2 | 36.0% |
| WCM44 | 4 | SRSF2 | c.284C>T; p.P95L | 35.0% |
| WCM44 | 5 | STAG2 | c.385+2T>C | 34.0% |
| WCM44_2 | 1 | BCOR | c.2428C>T; p.R810* | 28.0% |
| WCM44_2 | 2 | NRAS | c.35G>T; p.G12V | 32.0% |
| WCM44_2 | 3 | RUNX1 | c.921_925delCAGCGinsT; p.S308Afs*2 | 41.0% |
| WCM44_2 | 4 | SRSF2 | c.284C>T; p.P95L | 34.0% |
| WCM44_2 | 5 | STAG2 | c.385+2T>C | 30.0% |
| WCM44_3 | 1 | BCOR | c.2428C>T; p.R810* | 15.0% |
| WCM44_3 | 2 | NRAS | c.35G>T; p.G12V | 11.0% |
| WCM44_3 | 3 | RUNX1 | c.921_925delCAGCGinsT; p.S308Afs*2 | 12.0% |
| WCM44_3 | 4 | SRSF2 | c.284C>T; p.P95L | 15.0% |
| WCM44_3 | 5 | STAG2 | c.385+2T>C | 14.0% |
| WCM44_3 | 6 | ASXL1 | c.1772dupA; p.Y591* | 23.0% |

| | | | | | |
|---|---|---|---|---|---|
| WCM44_5 | None | | | | |
| WCM44_6 | None | | | | |
| WCM44_7 | | 1 | BCOR | c.2428C>T; p.R810* | 12.0% |
| WCM44_7 | | 2 | NRAS | c.35G>T; p.G12V | 9.0% |
| WCM44_7 | | 3 | RUNX1 | c.921_925delCAGCGinsT; p.S308Afs*2 | 14.0% |
| WCM44_7 | | 4 | SRSF2 | c.284C>T; p.P95L | 10.0% |
| WCM44_7 | | 5 | STAG2 | c.385+2T>C; splice-site | 12.0% |
| WCM45 | | 1 | SRSF2 | c.284C>T; p.P95L | 41.0% |
| WCM45_2 | | 1 | SRSF2 | c.284C>T; p.P95L | 41.0% |
| WCM46 | | 1 | DNMT3A | c.1627G>T; p.G543C | 22.0% |
| WCM46 | | 2 | TET2 | c.4642C>T; p.Q1548* | 25.0% |
| WCM46 | | 3 | U2AF1 | c.470A>G; p.Q157R | 27.0% |
| WCM46_2 | | 1 | DNMT3A | c.1627G>T; p.G543C | 36.2% |
| WCM46_2 | | 2 | TET2 | c.4642C>T; p.Q1548* | 36.6% |
| WCM46_2 | | 3 | U2AF1 | c.470A>G; p.Q157R | 34.7% |
| WCM46_3 | | 1 | DNMT3A | c.1627G>T; p.G543C | 8.0% |
| WCM46_3 | | 2 | U2AF1 | c.470A>G; p.Q157R | 4.0% |
| WCM46_4 | None | | | | |
| WCM47 | | 1 | PPM1D | c.1349delT, p.L450* | 7.2% |
| WCM49 | | 1 | RUNX1 | c.790C>T; p.Q264* | 32.0% |
| WCM49 | | 2 | SF3B1 | c.2098A>G; p.K700E | 31.0% |
| WCM49_2 | | 1 | SF3B1 | c.2098A>G; p.K700E | 38.3% |
| WCM49_2 | | 2 | RUNX1 | c.790C>T, p.Q264* | 40.1% |
| WCM49_2 | | 3 | RUNX1 | c.329A>G, p.K110R | 3.8% |
| WCM49_2 | | 4 | TET2 | c.2771A>G, p.H924R | 47.3% |
| WCM49_2 | | 5 | ZRSR2 | c.1338_1343dup6, p.S447_R448dup | 19.8% |
| WCM50 | | 1 | SF3B1 | c.2098A>G; p.K700E | 45.0% |
| WCM50 | | 2 | TP53 | c.659A>G; p.Y220C | 45.0% |
| WCM51 | | 1 | SF3B1 | c.2098A>G; p.K700E | 27.0% |
| WCM52 | | 1 | SF3B1 | c.2098A>G; p.K700E | 40.2% |
| WCM52 | | 2 | DNMT3A | c.2252T>C, p.F751S | 38.0% |
| WCM53 | | 1 | TET2 | c.2080_2081delCT, p.L694Yfs*17 | 13.7% |
| WCM53 | | 2 | CBL | c.1238G>A, p.G413D | 5.4% |
| WCM54 | None | | | | |
| WCM55 | | 1 | DNMT3A | c.1952_1961del10; p.K651Tfs*51 | 1.5% |
| WCM55 | | 2 | RIT1 | c.200A>G; p.E67G | 1.4% |
| WCM56 | None | | | | |
| WCM57 | | 1 | DNMT3A | c.1058_1059insT, p.F354Vfs*39 | 13.6% |
| WCM57 | | 2 | DIS3 | c.1653A>C, p.L551F, c.1653A>C | 15.2% |
| WCM71 | | 1 | TP53 | c.711G>A; p.M237I | 40.0% |
| WCM71 | | 2 | TP53 | c.733G>A; p.G245S | 36.0% |
| WCM72 | | 1 | PPM1D | c.1450_1451dupTT; p.L484Ffs*2 | 14.0% |
| WCM72 | | 2 | TP53 | c.584T>C; p.I195T | 38.0% |
| WCM73 | | 1 | TP53 | c.818G>A; p.Arg273His | 20.0% |
| WCM73 | | 2 | TP53 | c.524G>A; p.Arg175His | 30.7% |
| WCM73 | | 3 | TP53 | c.422G>A; p.Cys141Tyr | 32.0% |
| WCM73 | | 4 | SUZ12 | c.1079G>C; p.Arg360Pro | 4.6% |
| WCM77 | | 1 | SF3B1 | c.2098A>G; p.K700E | 7.7% |

**Supplemental Table 5. Summary of model architectures, training, and validation**

| Domain | Network Architecture | Prediction Method | Data Set | Validation Details | Metrics |
|---|---|---|---|---|---|
| Cell Detection Segmentaion | Mesmer: ResNet50 backbone with a multi-resolution feature pyramid. Two semantic heads for pixelwise and inner distance transforms. Model weights pretrained on TissueNet. | 256x256 overlapping patches MxIF channels | 1036 annotated 256x256 patches amounting to around 100,000 individual cell segmentations derived from training set of 6 BM WSI samples. Split 80%/20% into training and validation sets. | 340 annotated 256x256 image patches amounting to around 28,500 individual cell segmentations derived from hold-out set of 3 BM WSI samples. | Jaccard IoU: .71<br>Recall: .77<br>Precision: .91<br>F1-score: .83 |
| Membrane Marker Classificaton | Resnet18 with global average pooling layer connected to single neuron binary prediction output. Model weights pretrained on ImageNet. | 128x128 cropped patches centered on the cell of interest. Three channel images: cell segmentation, DAPI, and membrane marker channel of interest. | 872 single cell membrane images annotated as positive or negative expression. Split into 75%/25% training and validation sets. | 155 annotated single cell membrane images. | Accuracy: .97<br>Precision: 1.<br>Recall: .94<br>F1-score: .97 |
| TP53 Classification | Resnet18 with global average pooling layer connected to three neuron classification output. Model weights pretrained on Membrane Marker Classification dataset and then fine-tuned for nuclear expression classification. | 128x128 cropped patches centered on the cell of interest. Three-channel images: cell segmentation, DAPI, and TP53 channel. | 76 single cell TP53 nuclear expression images annotated as No Expression, Low Expression, Moderate/High Expression. Split into 80%/20% training and validation sets. | 25 annotated singe cell TP53 nuclear expression images. | Accuracy: .95<br>Precision: 1.<br>Recall: .86<br>F1-score: .92 |

**Supplemental Table 6. MDS-MAPS Additive Weight ( + Weight = MDS feature)**

| Feature | Weight |
| --- | --- |
| Myeloblast_Proportion | 1.000 |
| Megakaryocyte_Proportion | 0.981 |
| Megakaryocyte_Area | -0.959 |
| Erythroid_Area | 0.944 |
| Proerythro_Proportion | 0.925 |
| Erythroid_spatial_density | -0.914 |
| Hspc_endothelial_distance_ratio | 0.900 |
| Promyelocyte_endothelial_distance_ratio | 0.900 |
| Erythroid_K_ratio | -0.860 |
| Fat_to_cell_ratio | -0.852 |
| FatElements_per_mm2 | -0.817 |
| ALIP_Clusters | 0.760 |
| Megakaryocyte_Num_Nuclei | -0.757 |
| MMC_Proportion | -0.756 |
| Mast_Proportion | 0.750 |
| Myeloblast_endothelial_distance_ratio | 0.750 |
| Bcell_Proportion | -0.742 |
| Bcell_Area | 0.654 |
| Erythroid_Proportion | 0.635 |
| Promyelocyte_Proportion | 0.618 |
| HSPC_Proportion | 0.582 |
| Plasma_Proportion | 0.572 |
| Myeloblast_Area | -0.566 |
| Megakaryocyte_endothelial_distance_ratio | 0.500 |
| Promyelocyte_Area | -0.490 |
| Mean_fat_eccentricity | -0.486 |
| Endothelial_Proportion | 0.469 |
| Erythroid_Nuclei_Eccentricity | 0.440 |
| Hspc_25um_trabeculae | -0.360 |

#### SUPPLEMENTARY DATA FIGURES

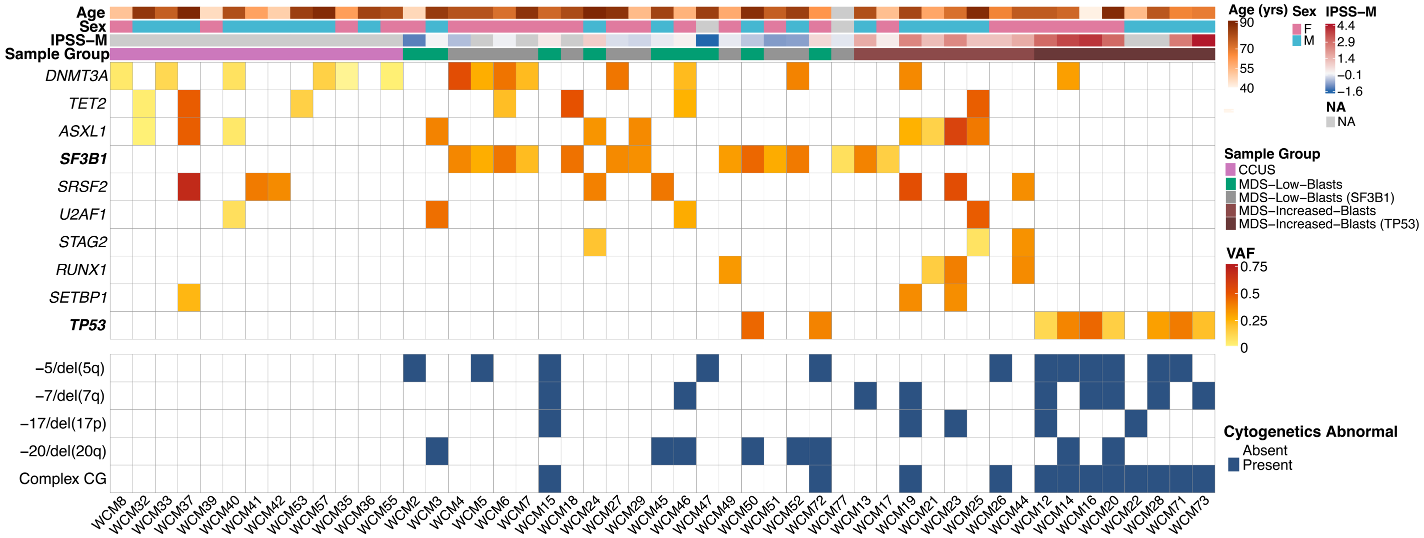

**Supplementary Fig. 1. Somatic mutation profiles and cytogenetic abnormalities in CCUS and MDS patients.** Stacked heatmap of patient metadata as each column. Top row of heatmap details patient age, sex, IPSS-M score (if applicable), sample group designated for each patient sample according to color legend on right-hand side. Patient sample groups listed are, in order: CCUS, MDS-LB and MDS-LB-SF3B1m, MDS-IB, MDS-IB-TP53m. Below patient metadata are results of targeted next-generation sequencing (only genes mutated in ≥10% samples are shown) with quantification of clonal mutation burden colored by variant allele frequency according to color bar on right-hand side. Lowest segment of heatmap are blue squares denoting cytogenetic abnormalities.

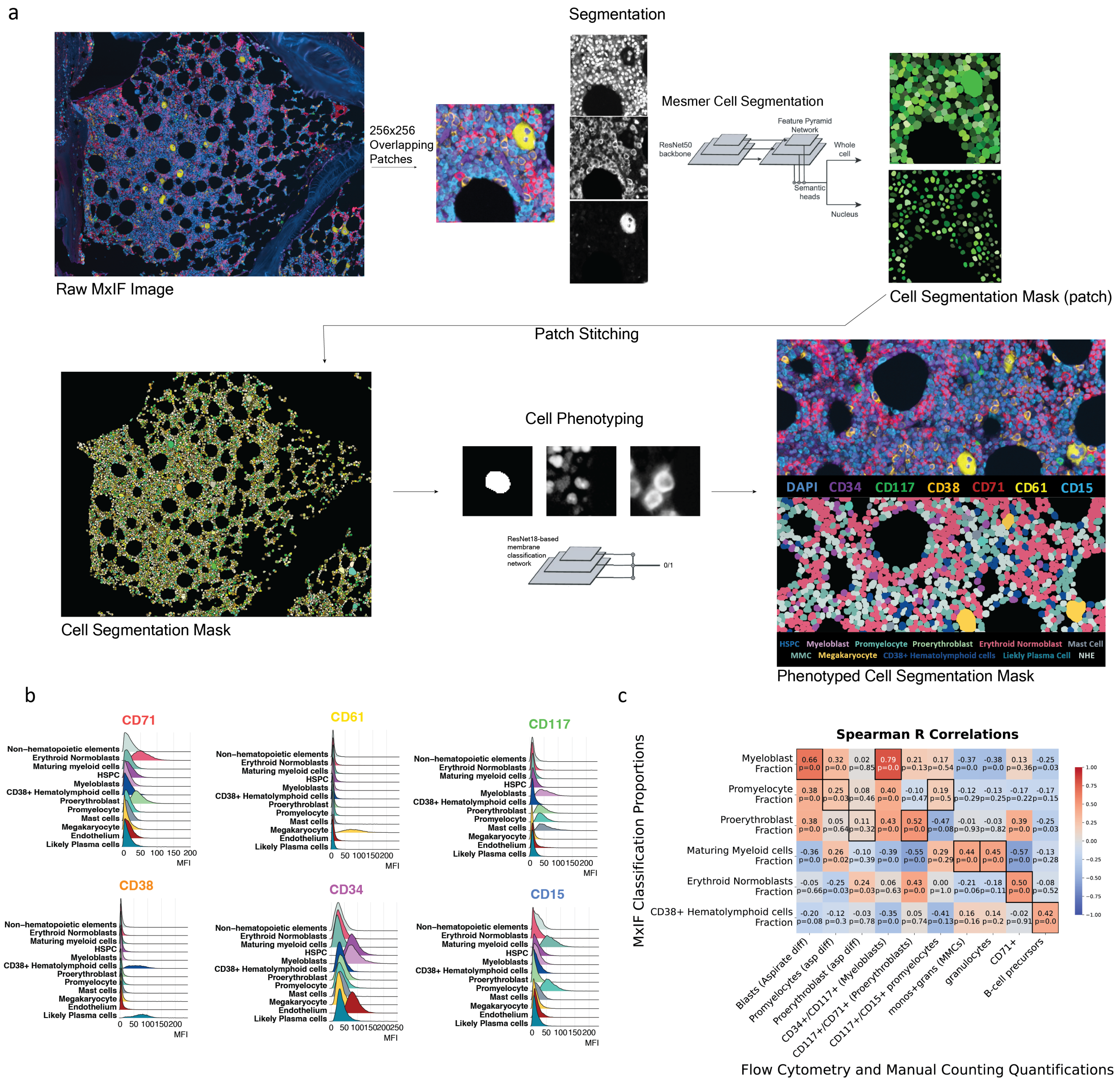

**Supplementary Fig. 2. Single-cell segmentation, phenotyping, and quantitative validation of the multiplex immunofluorescence analysis pipeline.**

**(a)** Schematic of the artificial intelligence–driven digital image analysis workflow. Shown are representative raw multiplex immunofluorescence (MxIF) image tiles (top left), patch-level Mesmer-based segmentation masks (top right), stitched whole-slide segmentation masks generated from overlapping 256 × 256 μm patches (bottom left), and convolutional neural network (CNN)–based cell phenotype classification masks used for downstream spatial analyses (bottom right). **(b)** Ridge plots showing distribution of mean fluorescence intensity (MFI) values across cell types classified by the CNN-based binary marker labeling algorithm, demonstrating concordance between predicted phenotypes and marker signal intensity distributions. **(c)** Correlation heatmap comparing MxIF-derived cell-type proportions with orthogonal measurements from manual differential counts and multiparameter flow cytometry. Spearman correlation coefficients (ρ) and corresponding P values are shown for each comparison.

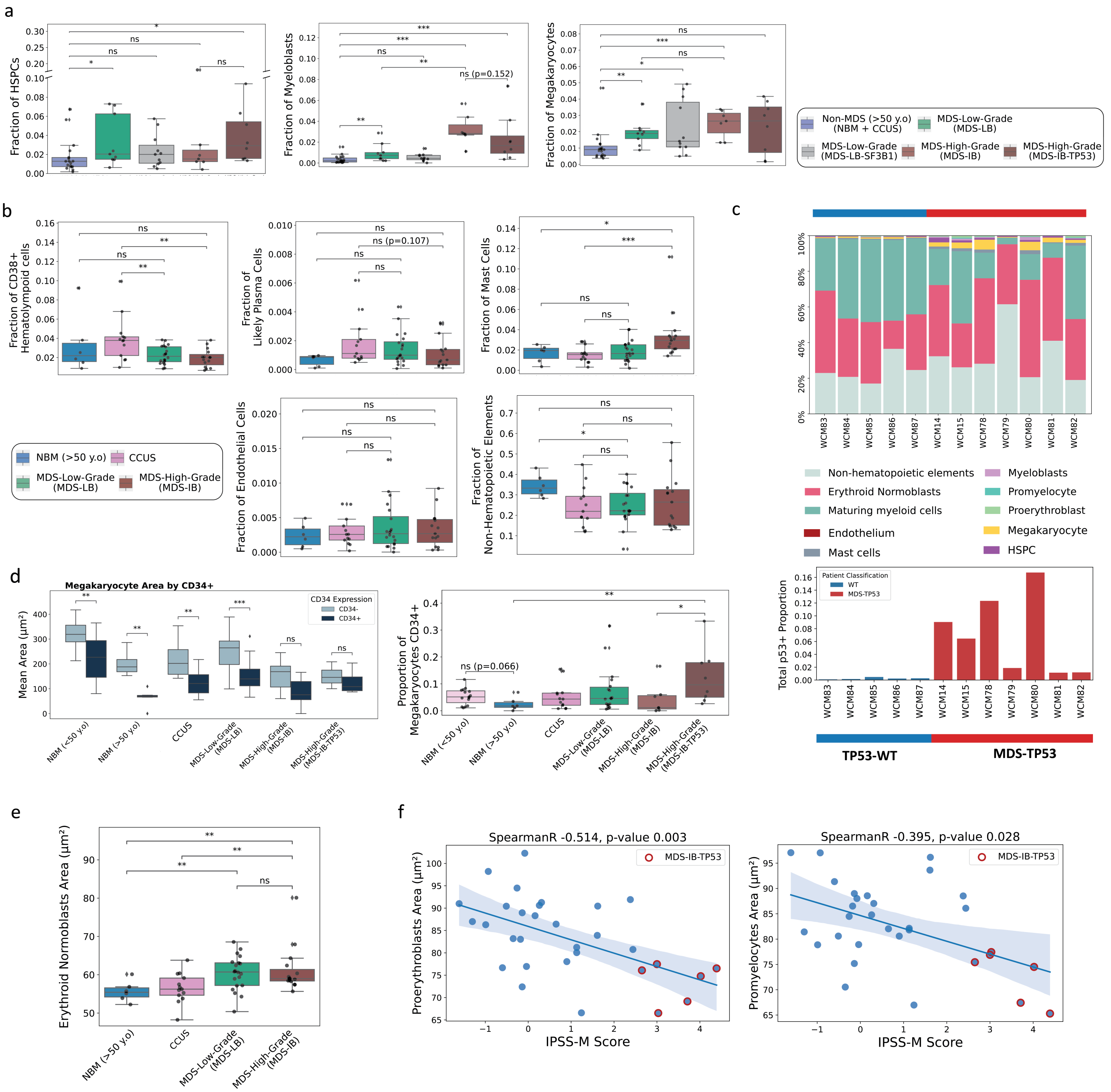

**Supplementary Fig. 3. Cell-type frequencies and morphologic characteristics across disease and genotype-defined groups.**

**(a)** Box plots showing proportions of hematopoietic stem and progenitor cells (HSPCs), myeloblasts, and megakaryocytes across genotype-defined sample groups. Statistical comparisons were performed using the Mann–Whitney U test (*$P \leq 0.05$, **$P \leq 0.01$, ***$P \leq 0.001$).

**(b)** Box plots showing proportions of $CD38^+$ hematolymphoid cells (non-plasma cells), mast cells, endothelial cells, and non-hematopoietic elements across sample groups (Mann–Whitney U test; *$P \leq 0.05$, **$P \leq 0.01$, ***$P \leq 0.001$).

**(c)** *Top*: Stacked bar plots showing cell-type proportion distributions in tissue microarray clot section samples. *Bottom*: Proportion of total cells exhibiting high p53 expression (putative *TP53*-mutated cells) within each sample.

**(d)** *Left*: Box plots comparing megakaryocyte cell area across sample groups stratified by CD34 expression status. *Right*: Proportion of $CD34^+$ megakaryocytes (mean fluorescence intensity >60) by sample group (Mann–Whitney U test; *$P \leq 0.05$, **$P \leq 0.01$, ***$P \leq 0.001$).

**(e)** Box plots comparing erythroid normoblast cell area across sample groups (Mann–Whitney U test; *$P \leq 0.05$, **$P \leq 0.01$).

**(f)** Scatter plot demonstrating significant inverse correlations between proerythroblast and promyelocyte cell area and IPSS-M risk score. TP53-mutated MDS-IB samples are highlighted (red circles).

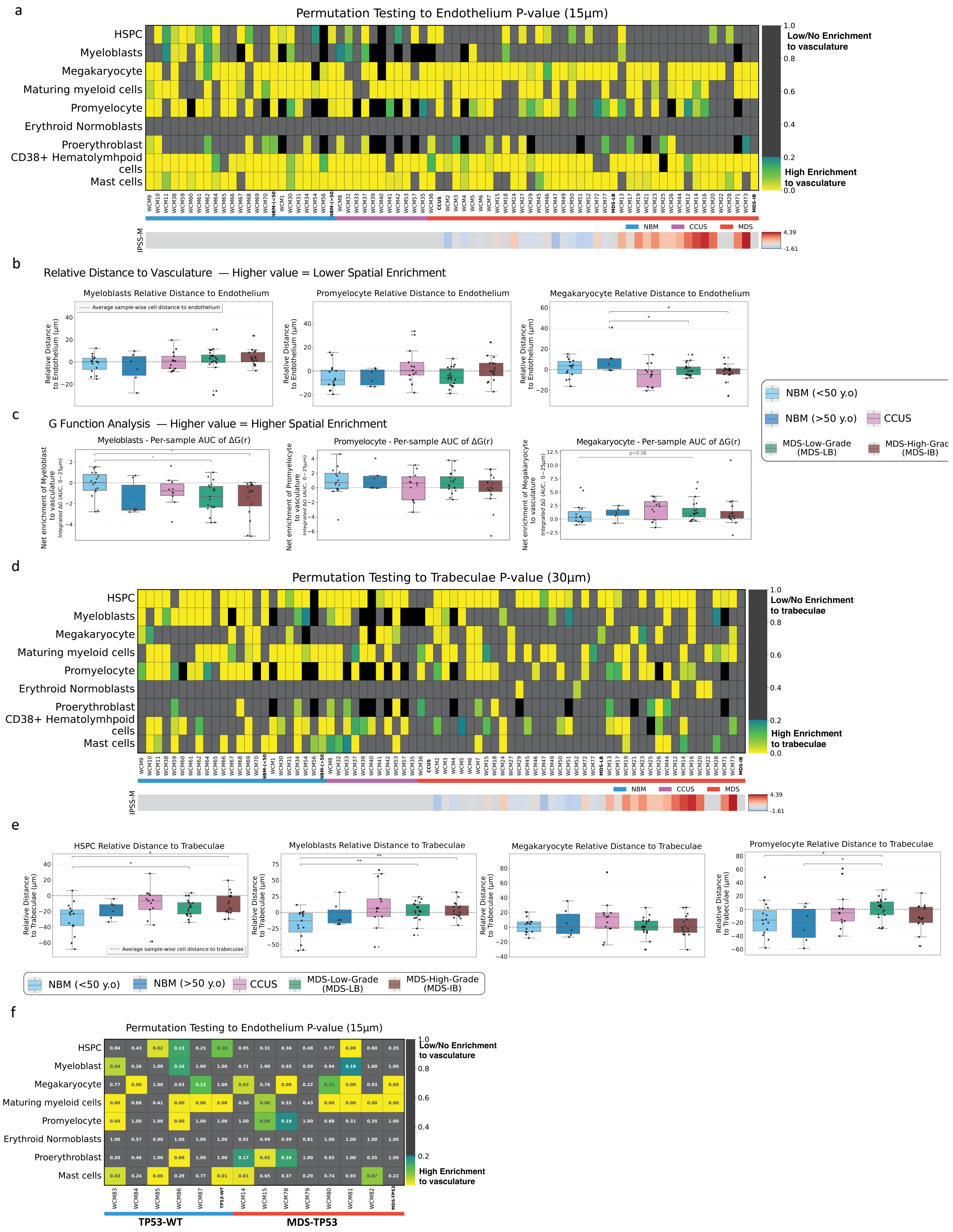
a
Permutation Testing to Endothelium P-value (15µm)
HSPC
Myeloblasts
Megakaryocyte
Maturing myeloid cells
Promyelocyte
Erythroid Normoblasts
Proerythroblast
CD38+ Hematolymhpoid cells
Mast cells
Low/No Enrichment to vasculature
High Enrichment to vasculature
NBM
CCUS
MDS
IPSS-M
b
Relative Distance to Vasculature — Higher value = Lower Spatial Enrichment
Myeloblasts Relative Distance to Endothelium
Promyelocyte Relative Distance to Endothelium
Megakaryocyte Relative Distance to Endothelium
Average sample-wise cell distance to endothelium
Relative Distance to Endothelium (µm)
NBM (<50 y.o)
NBM (>50 y.o)
CCUS
MDS-Low-Grade (MDS-LB)
MDS-High-Grade (MDS-IB)
c
G Function Analysis — Higher value = Higher Spatial Enrichment
Myeloblasts - Per-sample AUC of ΔG(r)
Promyelocyte - Per-sample AUC of ΔG(r)
Megakaryocyte - Per-sample AUC of ΔG(r)
Net enrichment of Myeloblast to vasculature
Net enrichment of Promyelocyte to vasculature
Net enrichment of Megakaryocyte to vasculature
Integrated ΔG (AUC, 0−25µm)
p=0.08
d
Permutation Testing to Trabeculae P-value (30µm)
Low/No Enrichment to trabeculae
High Enrichment to trabeculae
e
HSPC Relative Distance to Trabeculae
Myeloblasts Relative Distance to Trabeculae
Megakaryocyte Relative Distance to Trabeculae
Promyelocyte Relative Distance to Trabeculae
Relative Distance to Trabeculae (µm)
Average sample-wise cell distance to trabeculae
f
Permutation Testing to Endothelium P-value (15µm)
HSPC
Myeloblast
Megakaryocyte
Maturing myeloid cells
Promyelocyte
Erythroid Normoblasts
Proerythroblast
Mast cells
Low/No Enrichment to vasculature
High Enrichment to vasculature
TP53-WT
MDS-TP53

**Supplementary Fig. 4. Spatial proximity of hematopoietic cell types to vasculature and trabecular bone.**
**(a)** Heatmap of permutation-based P values showing enrichment of selected cell types within 15 μm of endothelial structures. Lower P values (yellow) indicate significant spatial enrichment relative to phenotype-label permutations. Individual samples are annotated by diagnostic group (NBM, CCUS, MDS-LB, MDS-IB); IPSS-M scores are shown for MDS samples. Bold column labels represent group-level combined P values calculated using Stouffer's method.
**(b)** Box plots showing relative distance to vasculature for selected cell types, defined as the mean cell-type distance to endothelium normalized to the mean distance of all other cell types within the same sample. Higher values indicate relative displacement (reduced enrichment). Statistical comparisons were performed using the Mann–Whitney U test (*P ≤ 0.05, **P ≤ 0.01, ***P ≤ 0.001).
**(c)** Box plots showing G-function based spatial enrichment test to vasculature for selected cell types. The value plotted is the AUC of G function curve representing the net enrichment of each cell type to vasculature, quantified by normalizing the proportion of each cell type within radius of vasculature to the expected proportion by random label shuffle and integrating this value over the range from 0 to 25 μm. Higher values indicate increased spatial enrichment. Statistical comparisons were performed using the Mann–Whitney U test (*P ≤ 0.05, **P ≤ 0.01, ***P ≤ 0.001)
**(d)** Heatmap of permutation-based P values showing enrichment of selected cell types within 30 μm of trabecular bone. Lower P values (yellow) indicate significant spatial enrichment.
**(e)** Box plots showing relative distance to trabeculae, defined as mean cell-type distance to trabecular mask normalized to the mean distance of all other cell types within the same sample (Mann–Whitney U test; *P ≤ 0.05, **P ≤ 0.01, ***P ≤ 0.001).
**(f)** Heatmap of permutation-based P values for enrichment within 15 μm of endothelium in the tissue microarray cohort containing *TP53*-mutated MDS and *TP53*–wild-type NBM samples. Columns represent individual samples; rows represent indicated cell types. Bold column labels denote group-level combined P values (Stouffer's method). HSPCs remain relatively enriched within 15 μm of endothelium in clot section material from NBMs represented on the tissue microarray.

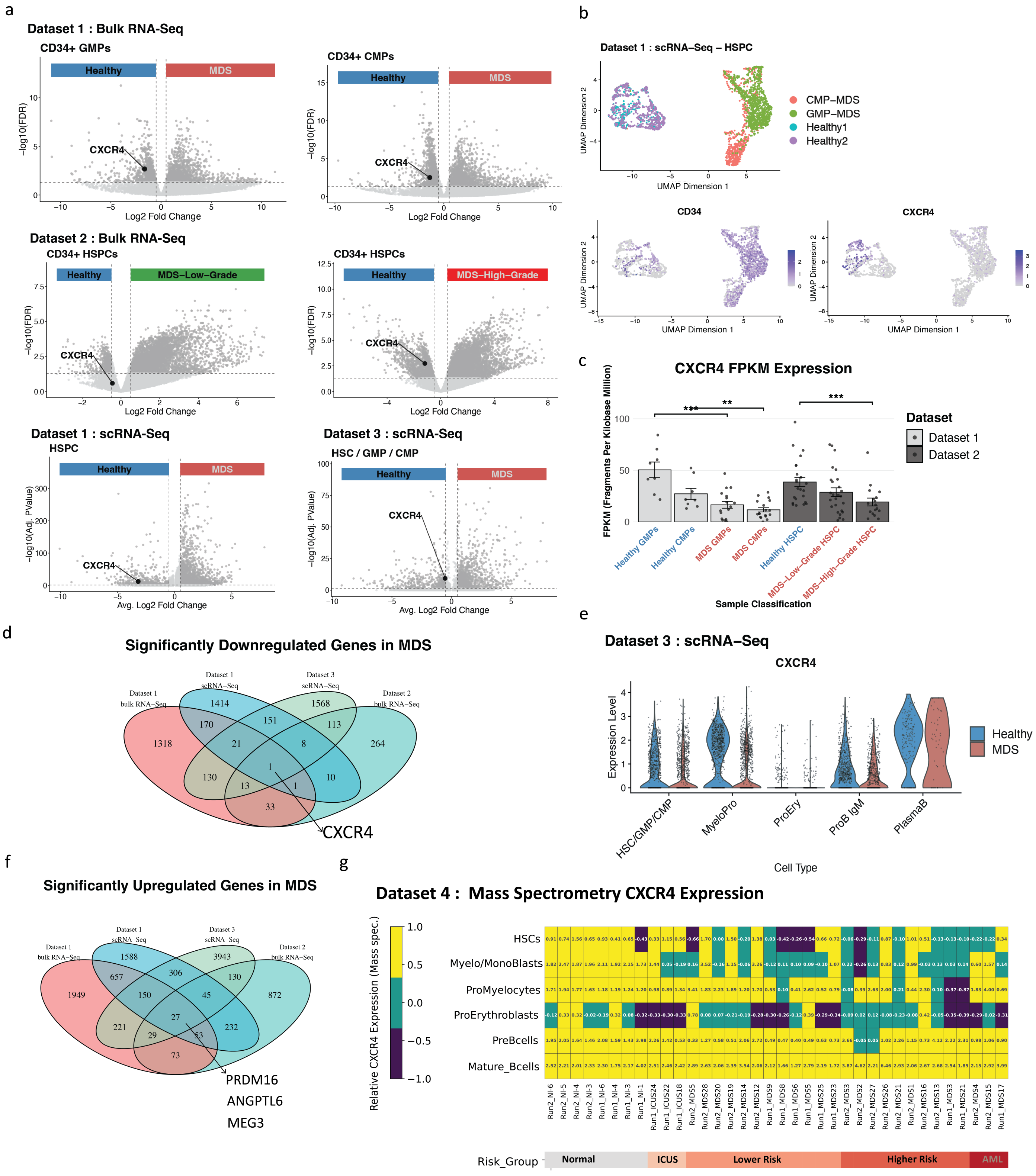

**Supplementary Fig. 5. Reduced CXCR4 expression in CD34+ progenitors across publicly available MDS transcriptomic and proteomic datasets.**

**(a)** Volcano plots showing differential gene expression between healthy and MDS CD34+ progenitors in four independent datasets from three separate publications. CXCR4 is highlighted as significantly down-regulated (FDR-adjusted P < 0.05 & abs(Log2 Fold Change) > .5) in Dataset 1 (GSE136816; bulk RNA-seq of GMPs and CMPs), Dataset 2 (GSE111085; bulk RNA-seq of CD34+ HSPCs, high-grade MDS), Dataset 3 (EGAS00001007568; single-cell RNA-seq of CD34+ HSPCs), and Dataset 1 single-cell RNA-Seq of HSPCs.

**(b)** UMAP embedding of CD34+ HSPCs from Dataset 1 single-cell RNA-seq, annotated by sample condition, with feature plots showing CD34 and CXCR4 expression.

**(c)** CXCR4 expression (FPKM) in bulk RNA-seq datasets comparing healthy versus MDS CD34+ progenitors: healthy vs MDS GMPs (adjusted P = 0.008), healthy vs MDS CMPs (adjusted P = 0.02), healthy vs MDS–low-grade HSPCs (adjusted P = 0.17), and healthy vs MDS–high-grade HSPCs (adjusted P = 0.008).
**(d)** Integration of differential expression results across four datasets (two bulk RNA-seq and two single-cell RNA-seq) identified CXCR4 as the only gene significantly and consistently down-regulated (FDR-adjusted P < 0.05) in all datasets.
**(e)** Violin plots from Dataset 3 single-cell RNA-seq showing CXCR4 expression across annotated CD34+ progenitor subsets in healthy versus MDS samples.
**(f)** Venn diagram illustrating overlap of significantly differentially expressed genes across datasets comparing healthy versus MDS CD34+ progenitors; 27 genes were consistently overexpressed in MDS, with selected genes labeled.
**(g)** Mass spectrometry–based quantification of CXCR4 protein abundance (Dataset 4) across defined progenitor populations. Columns represent individual samples ordered from healthy to high-risk MDS/AML (left to right); rows represent cell types. Higher values (yellow) indicate greater CXCR4 protein expression; lower values (gray/blue) indicate reduced expression.

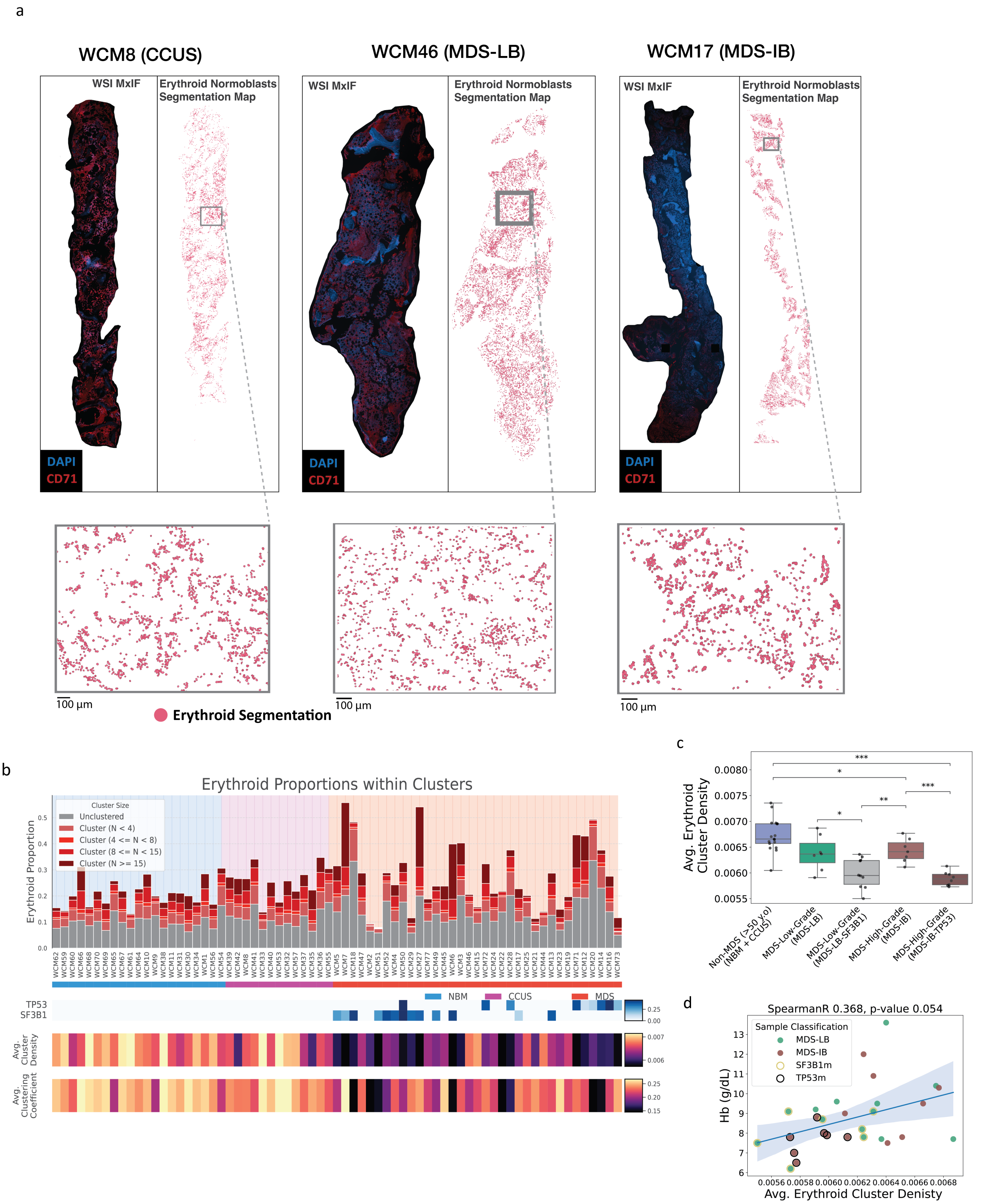

**Supplementary Fig. 6. Quantitative assessment of erythroid island architecture in normal and MDS bone marrow.**
**(a)** Representative whole-slide images displayed as merged DAPI and CD71 channels (left) and corresponding erythroid normoblast segmentation masks (right) for CCUS, MDS-LB, and MDS-IB samples. Insets show magnified regions of interest (ROIs) highlighting differences in erythroid spatial clustering.
**(b)** Stacked bar plots showing the distribution of erythroid normoblast cluster sizes per sample, derived using HDBSCAN unsupervised clustering. Bars represent the proportion of total erythroid normoblasts within each cluster size category. Cells not assigned to clusters or belonging to clusters of <4 cells are shown in gray; increasing cluster sizes are shown in red. Samples are

grouped as NBM (ordered by age), CCUS, and MDS (ordered by disease severity), as indicated by the color bar beneath the x-axis. *TP53* and *SF3B1* variant allele frequencies are annotated for each sample. For each patient, mean cluster density and clustering coefficient (calculated across clusters of >4 cells) are indicated.

**(c)** Box plots comparing mean erythroid cluster density across sample groups (Mann–Whitney U test; $^*P \leq 0.05$, $^{**}P \leq 0.01$, $^{***}P \leq 0.001$). Lower cluster density reflects dispersed, poorly formed erythroid aggregates; higher density indicates compact, well-formed erythroid islands.

**(d)** Scatter plot showing positive correlation between mean erythroid cluster density and peripheral hemoglobin concentration in MDS-LB and MDS-IB samples. *TP53*-mutated cases are denoted by black circles and SF3B1-mutated cases by yellow circles.

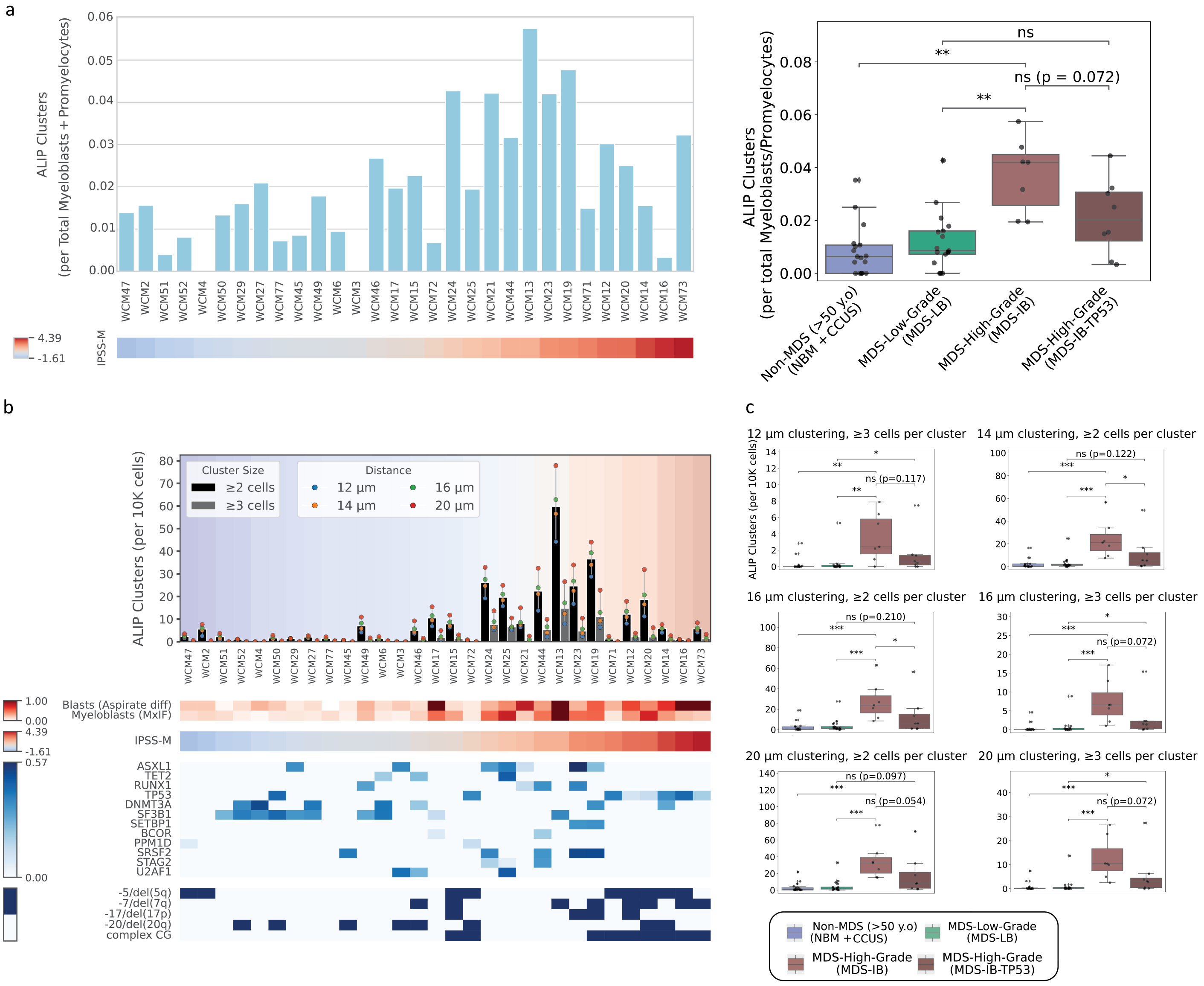

**Supplementary Fig. 7. Robustness analysis of atypical localization of immature precursors (ALIP).**
**(a)** Left: Bar plot showing ALIP cluster frequency per MDS sample (x-axis), normalized to the total number of promyelocytes and myeloblasts in each biopsy (y-axis). Right: Box plots comparing normalized ALIP cluster frequency across genotype-defined groups (Mann–Whitney U test; *P ≤ 0.05, **P ≤ 0.01, ***P ≤ 0.001).
**(b)** Sensitivity analysis of ALIP definition parameters. Bar plot showing ALIP cluster frequency per sample under eight parameter configurations, varying intercellular distance thresholds (12, 16, 18, and 20 µm) and minimum cluster size (≥2 or ≥3 cells). Black bars denote clusters of ≥2 cells; gray bars denote clusters of ≥3 cells. Colored dots indicate the corresponding distance threshold.
**(c)** Box plots comparing genotype-defined groups using six alternative ALIP parameter configurations (specified in subplot titles). Statistical comparisons were performed using Mann–Whitney U tests (*P ≤ 0.05, **P ≤ 0.01, ***P ≤ 0.001).

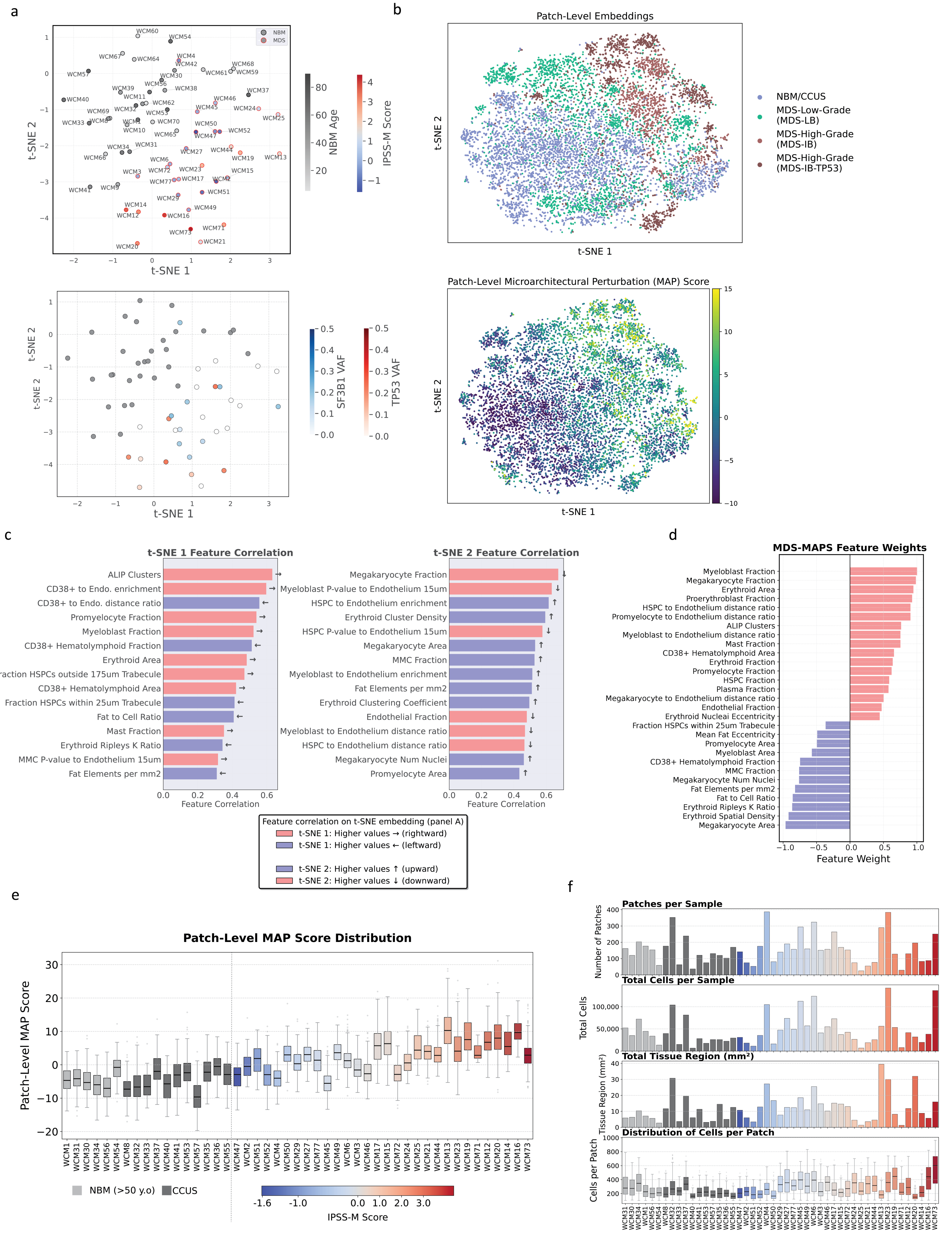

**Supplementary Fig. 8. Structure and quality control of the MDS Microarchitectural Perturbation Score (MDS-MAPS).**

**(a)** *Top*: t-distributed stochastic neighbor embedding (t-SNE) of sample-level features (corresponding to Fig. 6B), with individual samples labeled. *Bottom*: t-SNE plot annotated with *TP53* and *SF3B1* variant allele frequencies, highlighting a distinct MDS-IB-*TP53* cluster characterized by low t-SNE2 values.
**(b)** *Top*: t-SNE of 256 × 256 µm patch-level feature data. MDS-derived patches segregate from NBM and clonal hematopoiesis patches in patterns reflecting disease severity. *Bottom*: patch-level MDS-MAPS values (derived from additive weighting of features correlated with disease severity) overlaid as a color-coded feature map on the patch-level t-SNE projection.
**(c)** Bar plots showing features most strongly correlated with the t-SNE1 (left) and t-SNE2 (right) axes at the sample level. Red bars indicate features with strong positive correlation. Features positively correlated with t-SNE1 trend toward the MDS-IB cluster; features positively correlated with t-SNE2 trend toward the MDS-IB-*TP53* cluster.
**(d)** Bar plot of adjusted feature weights used in MAPS score computation. Correlative features from t-SNE sample stratification were curated by experts according to known MDS biology.
**(e)** Box plots showing within-sample distributions of patch-level MDS-MAPS values. Samples are grouped by diagnosis (NBM/CCUS in gray; MDS color-coded by IPSS-M risk score).
**(f)** Quality-control metrics per sample, including total patch count, total cell count, total analyzed tissue area ($mm^2$), and distribution of cells per patch (minimum 100 cells per patch threshold applied).

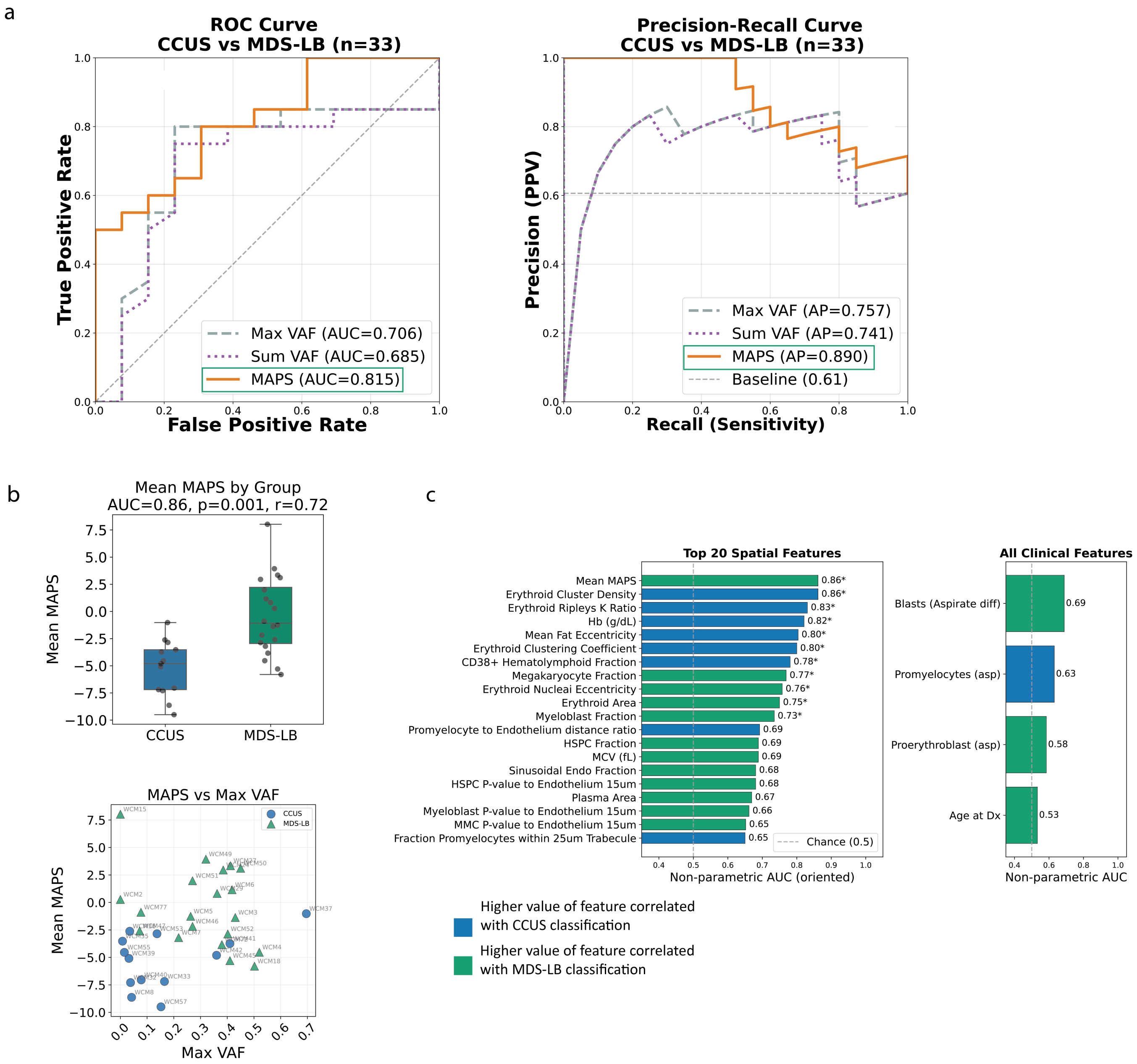

**Supplementary Fig. 9. MDS-MAPS discriminates CCUS and low-risk MDS trephine biopsies.**
**(a)** Leave-one-out cross-validation (LOOCV) for MDS-LB vs. CCUS classification (n = 33 samples). Receiver operating characteristic (ROC) and precision-recall analyses comparing MAPS against mutation burden features: Max VAF AUC = 0.706; Sum VAF AUC = 0.685; MAPS AUC = 0.815. Max VAF AP = 0.757; Sum VAF AP = 0.741; MAPS AP = 0.890. MAPS demonstrated superior discrimination relative to VAF-based mutation burden features across ROC and precision-recall metrics.
**(b)** *Top*: Box plot comparing baseline mean MAPS between CCUS and MDS-LB groups (CCUS median = -4.8; increase median = -1.1; Mann–Whitney U AUC = 0.86, P = 0.001; effect size r = 0.72). *Bottom*: Correlation between baseline MAPS and Max VAF from next-generation sequencing (n = 33 patients; Spearman ρ = 0.25, P = 0.17).
**(c)** Feature ranking by univariate discrimination capacity. *Left*: top 20 spatial features ranked by oriented nonparametric AUC. *Right*: clinical features ranked by oriented nonparametric AUC. Bar color indicates direction of association with MDS-LB classification.

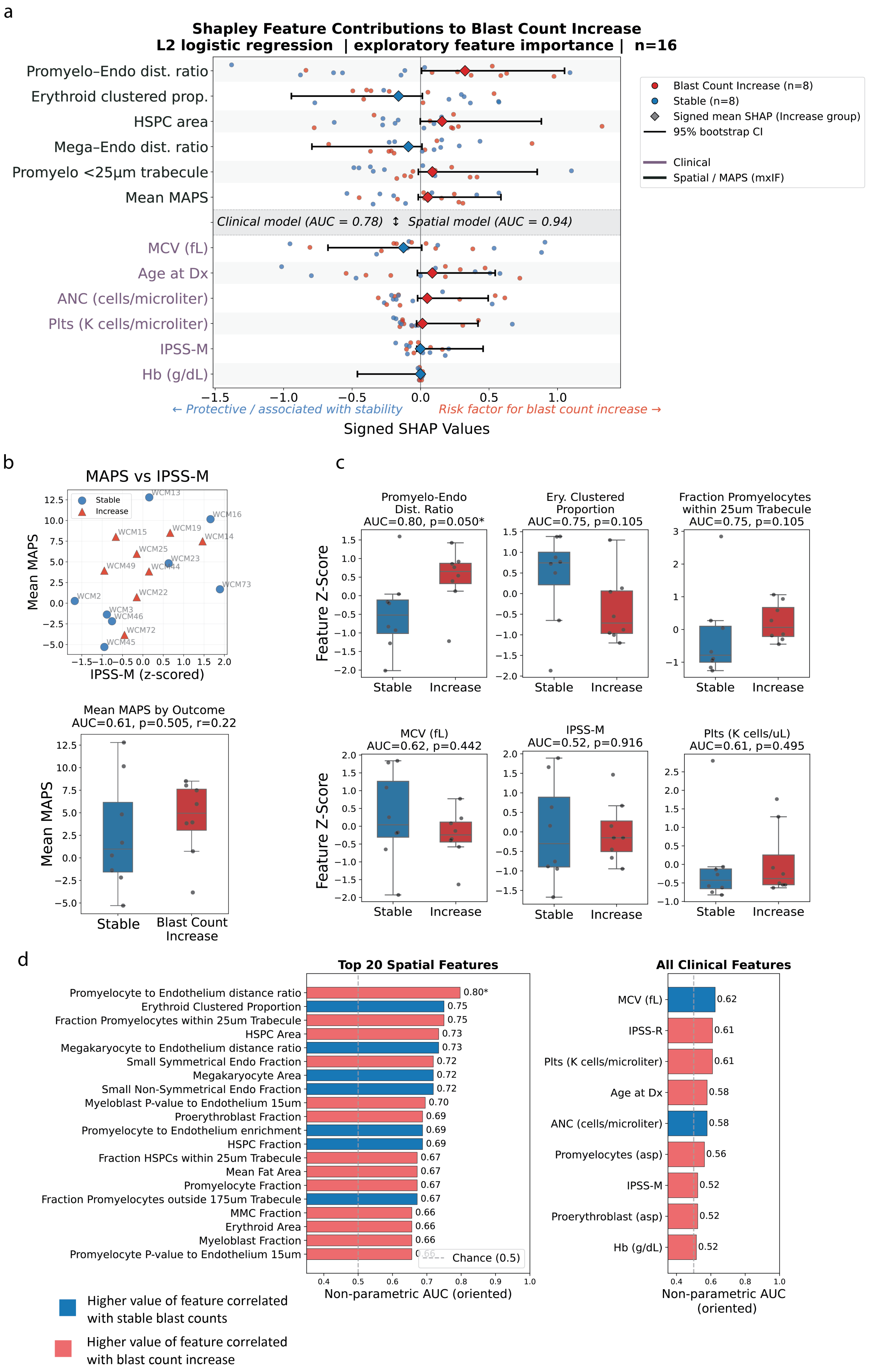

a
Shapley Feature Contributions to Blast Count Increase
L2 logistic regression | exploratory feature importance | n=16
Promyelo–Endo dist. ratio
Erythroid clustered prop.
HSPC area
Mega–Endo dist. ratio
Promyelo <25µm trabecule
Mean MAPS
Clinical model (AUC = 0.78) ↕ Spatial model (AUC = 0.94)
MCV (fL)
Age at Dx
ANC (cells/microliter)
Plts (K cells/microliter)
IPSS-M
Hb (g/dL)
Blast Count Increase (n=8)
Stable (n=8)
Signed mean SHAP (Increase group)
95% bootstrap CI
Clinical
Spatial / MAPS (mxIF)
← Protective / associated with stability
Risk factor for blast count increase →
Signed SHAP Values
b
MAPS vs IPSS-M
Stable
Increase
Mean MAPS
IPSS-M (z-scored)
Mean MAPS by Outcome
AUC=0.61, p=0.505, r=0.22
Mean MAPS
Stable
Blast Count Increase
c
Promyelo-Endo Dist. Ratio
AUC=0.80, p=0.050*
Ery. Clustered Proportion
AUC=0.75, p=0.105
Fraction Promyelocytes within 25um Trabecule
AUC=0.75, p=0.105
MCV (fL)
AUC=0.62, p=0.442
IPSS-M
AUC=0.52, p=0.916
Plts (K cells/uL)
AUC=0.61, p=0.495
Feature Z-Score
Stable
Increase
d
Top 20 Spatial Features
Promyelocyte to Endothelium distance ratio 0.80*
Erythroid Clustered Proportion 0.75
Fraction Promyelocytes within 25um Trabecule 0.75
HSPC Area 0.73
Megakaryocyte to Endothelium distance ratio 0.73
Small Symmetrical Endo Fraction 0.72
Megakaryocyte Area 0.72
Small Non-Symmetrical Endo Fraction 0.72
Myeloblast P-value to Endothelium 15um 0.70
Proerythroblast Fraction 0.69
Promyelocyte to Endothelium enrichment 0.69
HSPC Fraction 0.69
Fraction HSPCs within 25um Trabecule 0.67
Mean Fat Area 0.67
Promyelocyte Fraction 0.67
Fraction Promyelocytes outside 175um Trabecule 0.67
MMC Fraction 0.66
Erythroid Area 0.66
Myeloblast Fraction 0.66
Promyelocyte P-value to Endothelium 15um
Chance (0.5)
Non-parametric AUC (oriented)
All Clinical Features
MCV (fL) 0.62
IPSS-R 0.61
Plts (K cells/microliter) 0.61
Age at Dx 0.58
ANC (cells/microliter) 0.58
Promyelocytes (asp) 0.56
IPSS-M 0.52
Proerythroblast (asp) 0.52
Hb (g/dL) 0.52
Non-parametric AUC (oriented)
Higher value of feature correlated with stable blast counts
Higher value of feature correlated with blast count increase

**Supplementary Fig. 10. Exploratory modeling of baseline predictors of blast count increase.**
**(a)** SHAP (Shapley additive explanation) forest plots showing feature contributions to prediction of blast count increase from baseline biopsies (n = 16 patients; 8 increase, 8 stable). Two L2-regularized logistic regression models (C = 0.5) were evaluated: clinical features (purple; AUC = 0.78) and spatial/MxIF features (black; AUC = 0.94). Each dot represents a patient-level SHAP value (red = blast increase; blue = stable). Diamonds indicate sign-corrected mean SHAP values for the increase group; horizontal lines represent 95% bootstrap confidence intervals (1,000 resamples). Positive values indicate association with blast increase; negative values indicate association with stability. Features are ranked by absolute mean SHAP magnitude within each model.
**(b)** *Top*: Correlation between baseline MAPS and IPSS-M risk score (n = 13 patients; Spearman $\rho = 0.65$, P = 0.016). *Bottom*: Box plot comparing baseline mean MAPS between stable and blast-increase groups (stable median = 5.5; increase median = 7.5; Mann–Whitney U AUC = 0.61, P = 0.505; effect size r = 0.22).
**(c)** Box plots comparing z-scored feature values between stable and blast-increase groups for top discriminating spatial features. Nonparametric AUC was calculated as the normalized Mann–Whitney U statistic ($U / [n_0 \times n_1]$), representing the fraction of correctly ranked stable/increase patient pairs.
**(d)** Feature ranking by univariate discrimination capacity. *Left*: top 20 spatial features ranked by oriented nonparametric AUC. *Right*: clinical features ranked by oriented nonparametric AUC. Bar color indicates direction of association with blast increase.

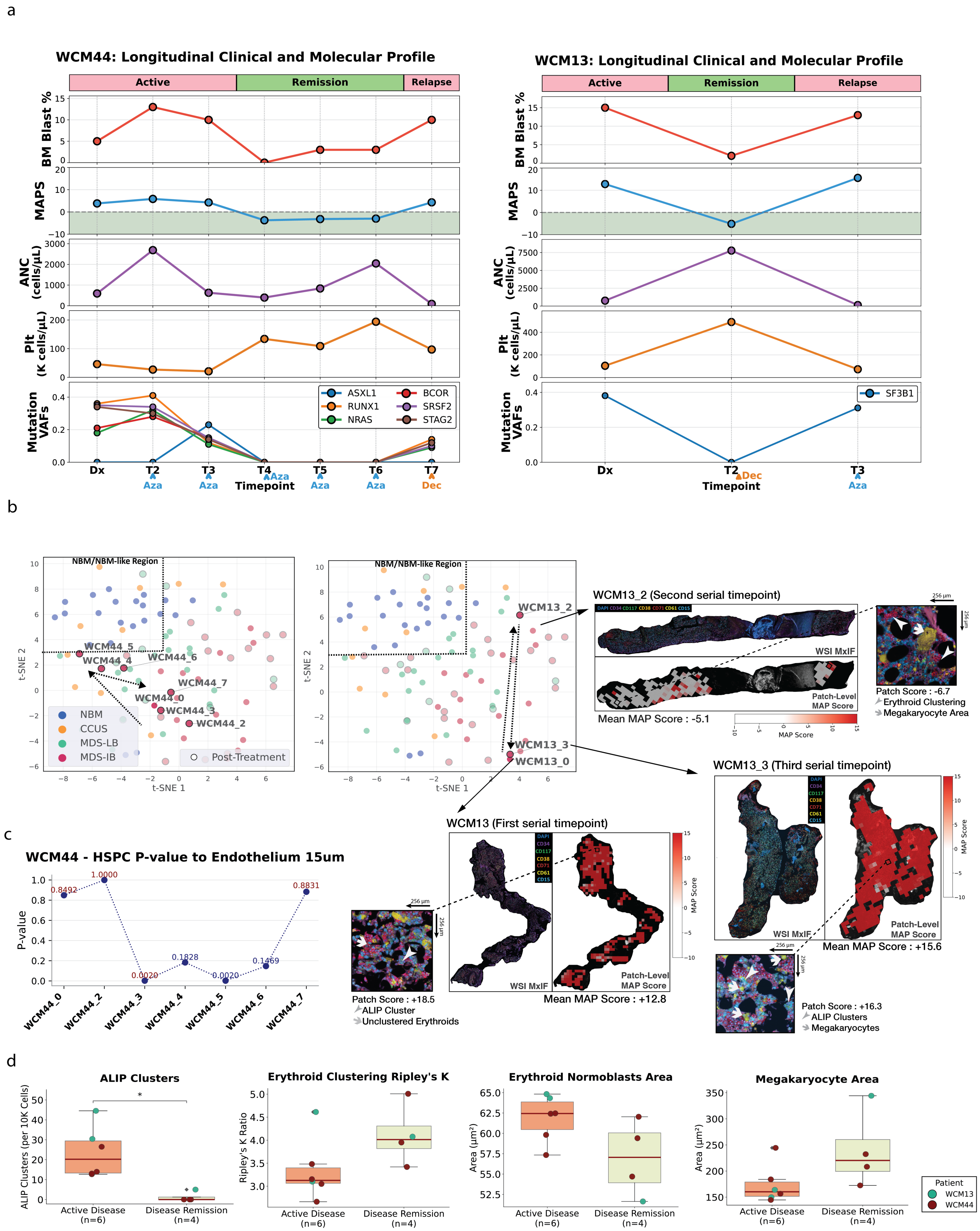

**Supplementary Fig. 11. Multimodal longitudinal assessment in patients with transient remission.**

**(a)** Longitudinal stacked line plots showing clinical parameters and MAPS over treatment course for two patients with transient remission: WCM44 (left; 7 biopsies) and WCM13 (right; 3 biopsies). Red bars denote active disease; light green bars denote remission. Treatment at biopsy timepoints is annotated (Aza, azacitidine; Dec, decitabine).
**(b)** t-SNE trajectories for WCM44 (left) and WCM13 (right), showing sample-level embeddings overlaid on the full cohort embedding of NBM and MDS cases. Post-treatment samples are highlighted with black circles. Regions enriched in Non-MDS samples are indicated. For WCM13, corresponding whole-slide MxIF images and MDS-MAPS heatmaps at each timepoint are shown, with arrows linking each biopsy to its t-SNE embedding position.
**(c)** Longitudinal assessment of HSPC spatial enrichment to endothelium in WCM44, shown as permutation testing p-values over time (low p-value indicates significant enrichment; high p-value indicates displacement).
**(d)** Box plots comparing sample-level quantifications of selected features between active disease and remission biopsies in WCM44 and WCM13. Individual points are color-coded by patient.
